# Single-Cell RNA Sequencing Reveals the Effects of Chemotherapy on Human Pancreatic Adenocarcinoma and its Tumor Microenvironment

**DOI:** 10.1101/2022.05.24.493132

**Authors:** Gregor Werba, Daniel Weissinger, Emily A. Kawaler, Ende Zhao, Despoina Kalfakakou, Surajit Dhara, Grace Oh, Xiaohong Jing, Nina Beri, Lauren Khanna, Tamas Gonda, Paul Oberstein, Cristina Hajdu, Cynthia Loomis, Adriana Heguy, Mara H. Sherman, Amanda W. Lund, Theodore H. Welling, Igor Dolgalev, Aristotelis Tsirigos, Diane M. Simeone

## Abstract

The tumor microenvironment (TME) in pancreatic ductal adenocarcinoma (PDAC) is a complex ecosystem that drives tumor progression; however, in-depth single cell characterization of the PDAC TME and its role in response to therapy is lacking. We performed single-cell RNA sequencing on freshly collected human PDAC samples either before or after chemotherapy. Overall, we found a heterogeneous mixture of basal and classical cancer cell subtypes, along with distinct cancer-associated fibroblast and macrophage subpopulations. Strikingly, classical and basal-like cancer cells exhibited similar transcriptional responses to chemotherapy, and did not demonstrate a shift towards a basal-like transcriptional program among treated samples. We observed decreased ligand-receptor interactions in treated samples, particularly TIGIT on CD8+ T cells and its receptor on cancer cells, and identified TIGIT as the major inhibitory checkpoint molecule of CD8+ T cells. Our results suggest that chemotherapy profoundly impacts the PDAC TME and may promote resistance to immunotherapy.

## Introduction

Pancreatic ductal adenocarcinoma (PDAC) is a highly lethal cancer with a five-year survival rate of approximately 11%^1^. With increasing incidence it is projected to become the second leading cause of cancer-related deaths in the United States by 2030^2^. Current treatment options are limited. Only 15% to 20% of patients qualify for upfront surgery with curative intent; the remaining patients present with unresectable locally advanced disease or distant metastases. Nearly all patients receive adjuvant, neoadjuvant, or palliative chemotherapy^3^. Recent advances in immunomodulating drugs, such as checkpoint inhibitors, have yet to show encouraging results in PDAC^4, 5^. A deeper understanding of pancreatic cancer biology is needed to develop better therapeutics and improve outcomes.

Although recent genomic and transcriptomic studies on bulk tumor samples identified key molecular drivers (notably KRAS, TP53, CDKN2A, and SMAD4) and cancer cell subtypes (basal and classical), these findings have yet to translate into treatment advances^6–9^. Recent investigations into the PDAC tumor microenvironment (TME) revealed that this complex ecosystem is a critical mediator of therapeutic resistance and tumor progression^10, 11^. Single-cell RNA sequencing (scRNA-seq) has emerged as a valuable tool in precision medicine by enabling in-depth characterization of the TME and serial assessment on scant amounts of tissue^12–17^. While recent single-cell studies have provided insights into PDAC, there is an unmet need to characterize the effects of chemotherapy on the TME^18–22^.

For a comprehensive view of the PDAC TME, and its response to chemotherapy in a real-time clinical context, we performed scRNA-seq on freshly collected tumor specimens from 27 PDAC patients – one of the largest scRNA-seq cohorts to date. We observed significant heterogeneity in the malignant epithelial compartment, with most tumors displaying a mixture of basal-like and classical-like cells. Furthermore, we examined the composition of CAFs and discrete groups of immune cells in the TME and executed comparative analyses on naive and chemotherapy-treated samples. Our data show that chemotherapy profoundly alters the PDAC TME, inducing features consistent with tumor progression and refractoriness to immunotherapy by downregulation of inhibitory checkpoint molecules in CD8+ T cells.

## Results

### Single-cell analysis reveals the transcriptomic landscape in PDAC

PDAC specimens were acquired via surgical resection of primary pancreatic lesions (n=10) or by endoscopic/interventional radiology–guided biopsies of either primary pancreatic lesions (n=7) or liver metastases (n=10) (Fig. 1A, Suppl. Table 1). Of the 27 total patients, seven received chemotherapy (FOLFIRINOX-based n=4, gemcitabine/abraxane-based n=3) prior to tissue collection, while 20 patients had not been treated at the time of specimen acquisition (Fig. 1A, Suppl. Table 1). We prepared single-cell suspensions from tumor specimens and sequenced them using the 10X Genomics Chromium system (Fig. 1B, see Methods for details). Histopathological assessment was done on each tumor specimen to verify malignancy, and samples underwent mutational profiling (Suppl. Table 2, Extended Data). After standard processing and quality filtering of the raw sequencing data (see Methods), a total of 139,446 cells were retained for analysis. Unsupervised, graph-based clustering and visualization via uniform manifold approximation and projection (UMAP) revealed ten distinct cell clusters that were annotated using canonical markers and gene expression profiles (Fig. 1C-D, Supp. Fig. 1, Methods). Three major clusters were identified as epithelial, T/natural killer (NK), and myeloid cells. We further identified CAFs, endothelial cells, B/plasma cells, and a small mast cell population among our clusters. The proportional distribution of these major cell types varied widely between samples (Fig. 1E).

**Figure 1.**
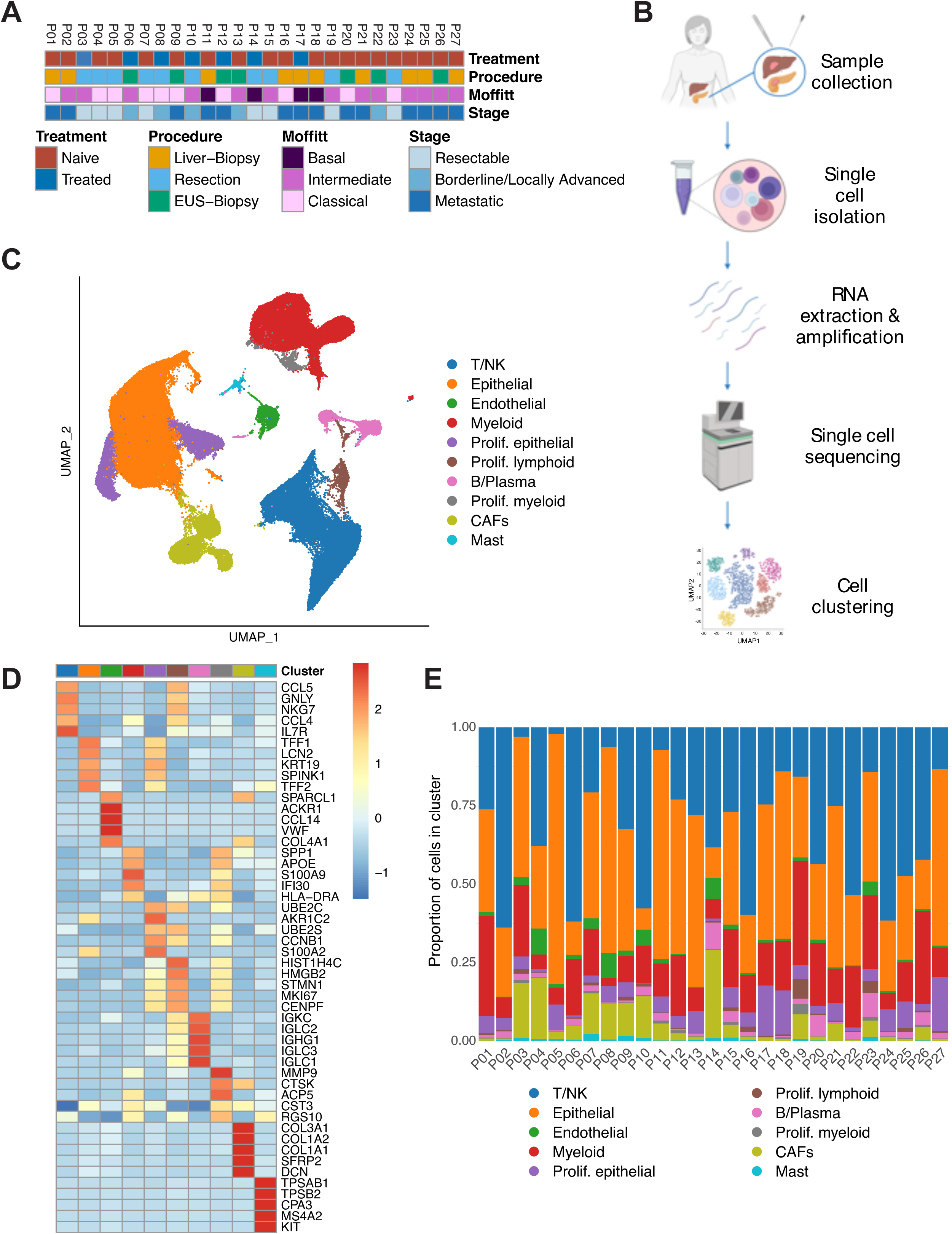
Single-cell analysis reveals the transcriptomic landscape in PDAC. A. Clinicopathologic characteristics for each patient sample including treatment status, stage, Moffitt subtype, and procedure. EUS, endoscopic ultrasound. B. Workflow depicting sample acquisition, processing, and analysis. Designed with BioRender**©**. C. UMAP embedding of the expression profile of 139,446 cells that passed quality control from n=27 patient samples. Distinct clusters are annotated and color-coded. CAF, cancer-associated fibroblast; NK, natural killer; UMAP, uniform manifold approximation and projection. D. Heatmap of the five most differentially expressed genes within each cluster. Colors correspond to the cluster colors in Figure 1C. E. Proportional distribution of each cell cluster by individual sample.

### The epithelial compartment reveals heterogeneous malignant subtype composition

The epithelial compartment evinced sample-specific differences while the mesenchymal (CAF and endothelial) and immune (T/NK and myeloid) compartments clustered together across all samples (Suppl. Fig. 2A). This suggests that non-epithelial cells of the PDAC TME share more common transcriptional signatures across all patients than their malignant counterparts within the same tumor. Since some epithelial cells also clustered across samples, we further assessed the proportion of malignant cells in the total epithelial compartment (Fig. 2A). To this end, we performed copy number variation (CNV) analysis. The vast majority of epithelial cells showed several CNV events, indicating malignancy and distinguishing them from normal epithelial cells (Fig. 2B, Suppl. Fig. 2B). We further classified malignant cells as either basal or classical based on their Moffitt subtype signature expression^7^. All samples but one were comprised of a mixture of basal and classical cancer cells in varying proportions (Fig. 2C-D). Tumor subtype composition was not correlated with the amount of any major non-malignant cell types (Suppl. Fig. 2C).

**Figure 2.**
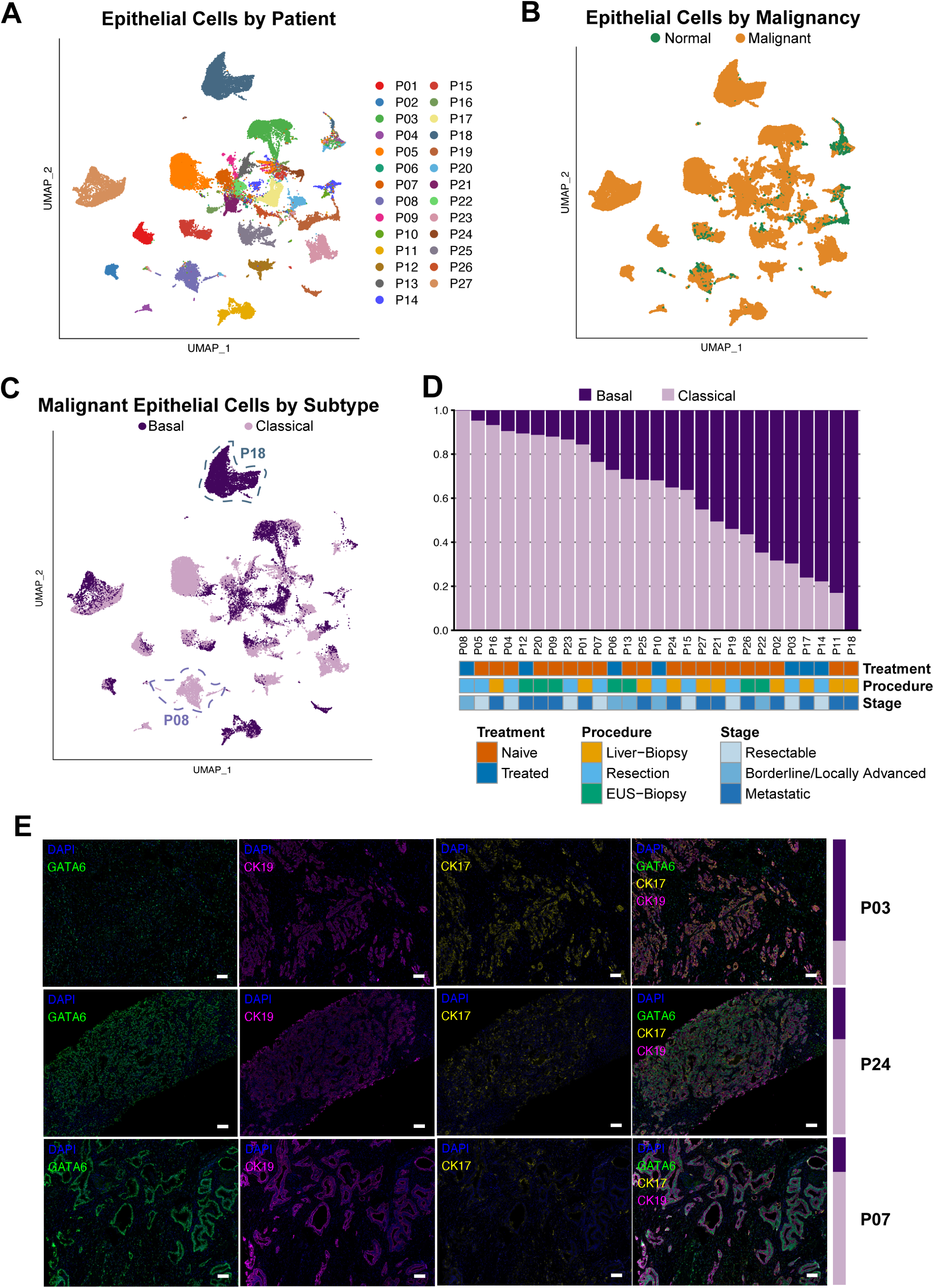
The epithelial compartment reveals heterogeneous malignant subtype composition. A. Non-batch corrected UMAP of all epithelial cells, showing clustering by patient sample of most epithelial cells and clustering of some epithelial cells across samples. UMAP, uniform manifold approximation and projection. B. Non-batch corrected UMAP of malignant and normal epithelial cells as determined by InferCNV. C. UMAP embedding of malignant epithelial cells labeled with Moffitt subtypes. Exemplary samples high in basal (P18) and classical (P08) cells are annotated. D. Proportion of cells of each Moffitt subtype by sample. EUS, endoscopic ultrasound. E. Representative images of multiplex immunofluorescence from cases high in basal (upper panel, P03), mixed (middle panel, P24), and classical (lower panel, P07) subtype tumors. Basal to classical ratio for each sample by scRNA-seq transcriptional analysis is shown on the right as a colored bar (dark = basal, light = classical). Channels (always including DAPI (blue)): GATA6 (green), CK19 (cytokeratin 19) (violet), CK17 (cytokeratin 17) (yellow), and merged. Scale bar = 100 μm.

We validated the Moffitt transcriptional subtypes on formalin-fixed, paraffin-embedded tissue (FFPE) sections using multiplex immunohistochemistry (IHC). We used the canonical markers: cytokeratin 17 (CK17) for the basal subtype, GATA6 for the classical subtype, and cytokeratin 19 (CK19) as a pan-cancer cell marker. Representative samples with a higher percentage of basal or classical transcriptional subtypes were strongly positive for CK17 or GATA6, respectively (Fig. 2E, upper and lower panels), while a sample with a more even mixture of both subtypes showed an interspersed, more equal expression of GATA6 and CK17 (Fig. 2E, middle panels). Another sample with a mixture of both subtypes showed two distinct growth patterns with a CK17^high^ region next to a GATA6^high^ region (Suppl. Fig. 2D). In general, GATA6 staining was more omnipresent, and CK17^high^ cells often stained positive for GATA6. Of note, most samples included cells staining positive for both CK17 and GATA6 (Suppl. Fig. 2E), consistent with a previous report that hypothesized an intermediate cell state^23^.

To test this intermediate cell state hypothesis in our dataset, we applied the basal, classical, and intermediate gene signatures from that report to malignant cells from our treatment-naive PDAC liver metastatic samples. Although their basal and classical signatures matched up closely with ours, we did not see clear evidence of the intermediate signature (Suppl. Fig. 2F). Further characterization of basal and classical cells by differential gene expression showed typical basal and classical genes upregulated in the respective subtypes (Suppl. Fig. 2G). Pathway enrichment analysis revealed enrichment of EMT, angiogenesis, and extracellular matrix (ECM) interaction—features related to migration and invasion—in basal cells, while classical cells were enriched for metabolic pathways, especially lipid metabolism (Suppl. Fig. 2H). These results emphasize the heterogeneity of malignant epithelial cells in PDAC and suggest that single-marker analysis may be insufficient to accurately identify cancer cell subtypes within a patient’s tumor.

### iCAF and myCAF subpopulations are not linked to Moffitt tumor subtype distribution

We next analyzed the mesenchymal compartment of the PDAC TME. Mesenchymal cell reclustering revealed seven distinct clusters (Fig. 3A). The two main clusters consisted of myofibroblastic CAFs (myCAFs) and inflammatory CAFs (iCAFs) (Fig. 3A-B)^24^. A third proposed CAF subtype, antigen-presenting CAFs (apCAFs), was not included among our clusters. Therefore, we instead looked specifically for cells with strong apCAF signatures^19^. While iCAFs and myCAFs matched with their respective signatures, we did not find a strong antigen-presenting CAF (apCAF) signature (Fig. 3C). Notably, the apCAF signature was derived from mouse CAFs and may not translate to the human CAFs in our cohort. As apCAF features in human CAFs have been linked to co-expression of CD74 and human leukocyte antigen (HLA) molecules, we checked for expression of these genes in our CAF populations; a small number of cells in both the iCAF and myCAF subpopulations co-expressed CD74 and HLA genes (Fig. 3C, Suppl. Fig. 3A)^19^. Other clusters were identified as pericytes and Schwann cells, while one sample provided a cluster of chondrocyte-like cells. Additionally, we detected a small proportion of cells expressing epithelial markers (Fig. 3A-B). We ruled out these cells being cancer cells undergoing EMT due to a lack of CNVs, low expression of EMT-related transcription factors (TFs) SNAI1 and TWIST1, and low values of an EMT-TF independent partial EMT (pEMT) program signature^25^. We labeled this cluster “epithelial-like” (Suppl. Fig. 3B).

**Figure 3.**
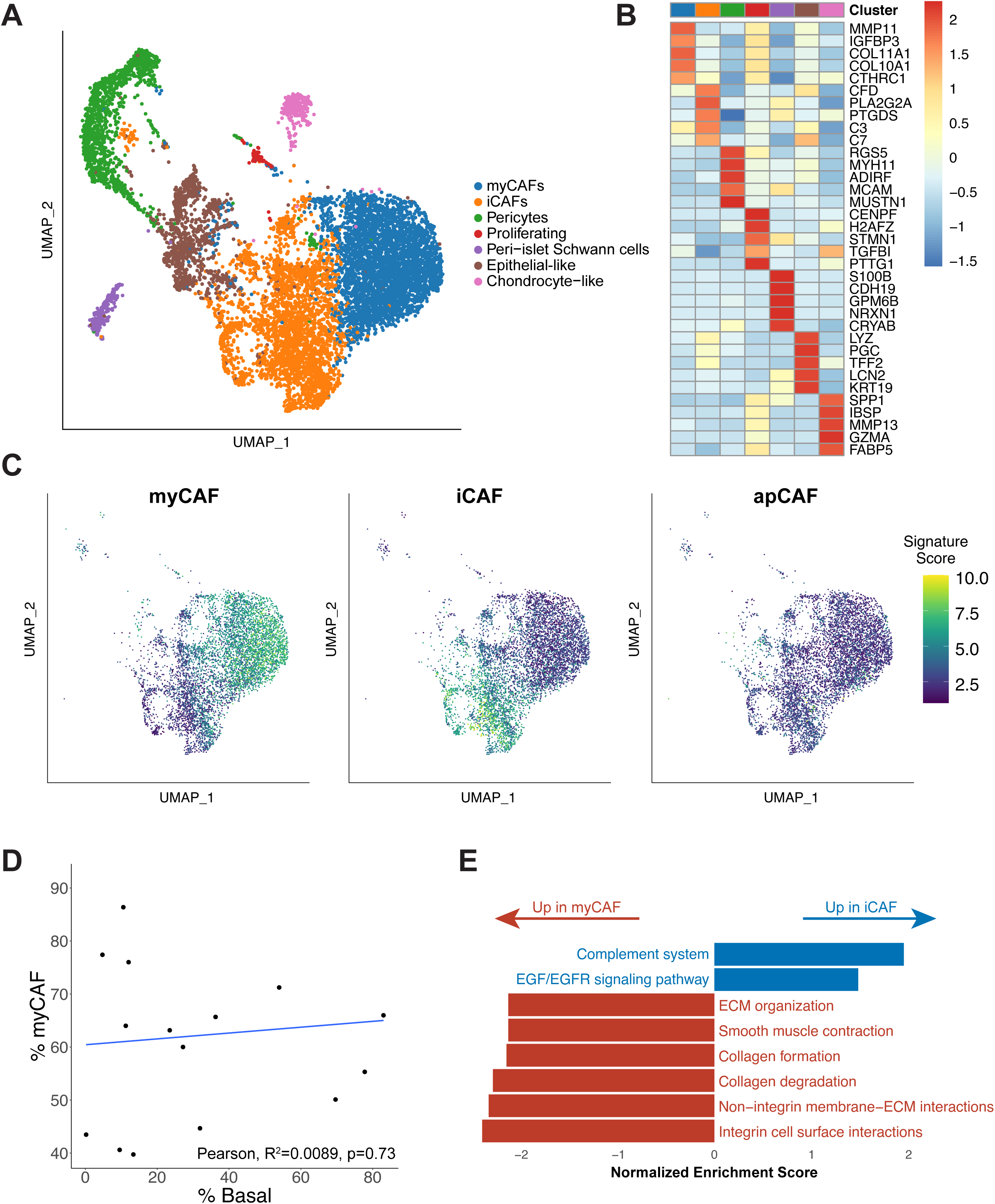
iCAF and myCAF subpopulations are not linked to Moffitt tumor subtype distribution. A. UMAP overview of the mesenchymal compartment. CAF, cancer-associated fibroblast; iCAF, inflammatory CAF; myCAF, myofibroblastic CAF; UMAP, uniform manifold approximation and projection. B. Heatmap of the five most differentially expressed genes within each cluster. Colors correspond to the cluster colors in panel A. C. iCAF, myCAF, and apCAF signatures from Elyada *et al.* overlaid on the UMAP embedding of our mesenchymal compartment. The iCAF and myCAF signatures are derived from human fibroblasts; the apCAF signature is derived from KPC mouse fibroblasts. apCAF, antigen-presenting CAF. D. Assessment of correlation between myCAF/iCAF composition and basal/classical composition among our samples. Samples are plotted as a function of the percentage of their CAF cells that are myCAFs and the percentage of their malignant epithelial cells that are basal. Samples with fewer than 40 total CAF cells were excluded; all but one of these samples were biopsies, which were not expected to pick up many mesenchymal cells. E. Selected gene set enrichment analysis results from a comparison between the cells in the iCAF and myCAF compartments. ECM, extracellular matrix.

The myCAF to iCAF ratio varied across samples, while the proportional distribution of both subpopulations in relation to Moffitt subtype distribution was not significantly different (Fig. 3D). In myCAFs, SDC1^26^, a gene linked to CAF-mediated regulation of breast cancer cell migration, was upregulated, while in iCAFs, CCL-19^27^, a gene linked to cytotoxic T cell recruitment by CAFs in lung cancer, was highly expressed (Suppl. Fig. 3C). Additionally, iCAFs showed enrichment for genes involved in the complement cascade and EGF signaling, whereas myCAFs demonstrated enrichment for ECM organization and smooth muscle contraction, supporting the notion of CAF heterogeneity and distinct functional characteristics of CAFs in the TME (Fig. 3E).

### Charting the T/NK Landscape in the PDAC Tumor Microenvironment

To better characterize tumor-infiltrating lymphocytes, we analyzed the T/NK population and identified 13 distinct clusters (Fig. 4A). CD4+ T cells clustered into four distinct subtypes: i) CCR7+CD4+ (high expression of naïve/central memory markers), ii) IL7R+CD4+ (high expression of activation markers), iii) FOXP3+CD4+ (high expression of regulatory T cell genes), and iv) CXCL13+CD4+ (high expression of follicular helper genes) (Fig. 4B). CD8+ T cells clustered into three main subpopulations: i) GZMH+CD8+ (high in cytotoxic gene expression), ii) GZMK+CD8+ (expressing effector memory genes), and iii) ITGA1+CD8+ (expression of tissue residency and exhaustion markers). We also identified a small ISG15+ cluster, shared between CD4+ and CD8+ T cells, with high interferon (IFN)-related gene expression. NK cells separated into a GNLY+ cluster highly expressing cytotoxic genes and an XCL1+ cluster with lower cytotoxic gene expression and higher expression of genes previously linked to tissue-resident NK cells^28, 29^. Smaller clusters were identified as mast cells and plasma cells (Fig. 4A-B). T/NK subpopulations were heterogeneously distributed across samples and did not correlate with basal/classical subtyping (Suppl. Fig. 4A-B).

**Figure 4.**
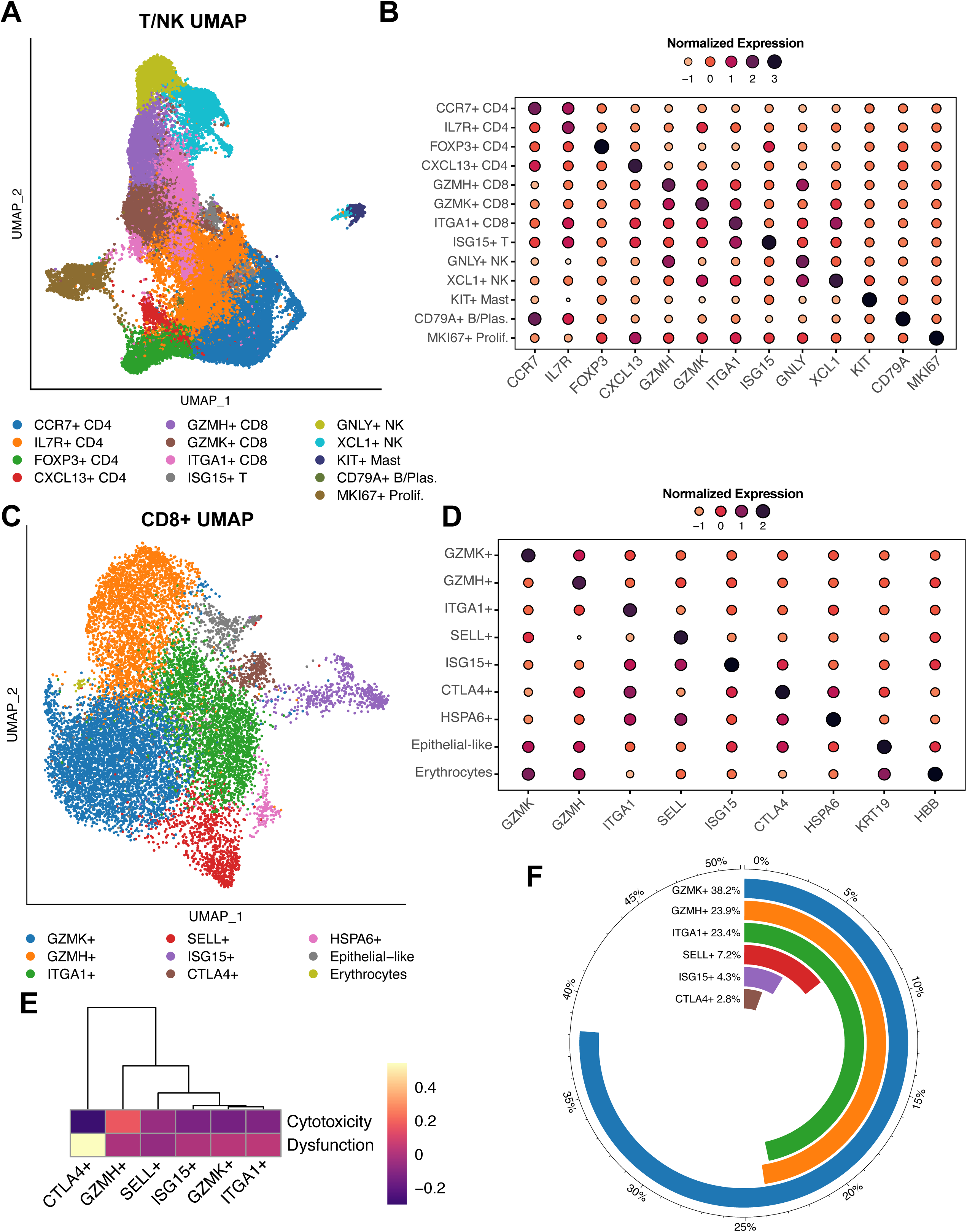
Charting the T/NK Landscape in the PDAC Tumor Microenvironment. A. UMAP overview of the T/NK compartment. NK, natural killer; UMAP, uniform manifold approximation and projection. B. The most differentially expressed gene within each T/NK cluster. C. UMAP of the more granular CD8+ T cell reclustering reveals additional subpopulations. D. The most differentially expressed gene within each CD8+ T cell cluster. E. Hierarchical clustering of the CD8+ T cell subsets based on dysfunction and cytotoxicity scores. F. Composition of the CD8+ T cell subset (excluding the two non-T cell clusters and the single patient cluster).

Because our initial T/NK clustering did not identify distinct exhausted T cell subpopulations, we refined the resolution of the CD8+ T cell clustering (Fig. 4C). At this finer granularity, we discerned a small CTLA4+CD8+ T cell cluster with a high level of exhaustion markers (CTLA4, HAVCR2, LAG3, TIGIT). Additional distinct CD8+ T cell clusters included i) GZMK+CD8+ (high expression of effector memory genes), ii) GZMH+CD8+ (high expression of cytotoxic genes), iii) ITGA1+CD8+ (high expression of tissue residency genes), iv) SELL+CD8+ (expression of naive/central memory genes), and v) ISG15+CD8+ (expressing IFN activation genes) (Fig. 4C-D). We also found an epithelial-like cluster, an erythrocyte-like cluster, and an HSPA6+ cluster; the latter was mainly derived from one patient and did not show other distinct CD8+ subpopulation features (Fig. 4C-D).

To further assess exhaustion in our CD8+ T cell subpopulations, we scored each subset on a dysfunction and cytotoxicity scale^30^. As expected, GZMH+CD8+ T cells scored highest in cytotoxicity, and CTLA4+CD8+ T cells highest in exhaustion (Fig. 4E). Except for CTLA4+CD8+ T cells, the exhaustion state was similar between subsets (Fig. 4E). Exhausted CD8+ T cells represented only 2.8% of all CD8+ T cells (Fig. 4F). Moreover, exhausted T cells, along with most other CD8+ T cell subsets, were heterogeneously distributed across samples, independent of the Moffitt subtype composition of their epithelial compartments (Suppl. Fig. 4C). Of note, ITGA1+CD8+ T cells were the only subpopulation that was significantly negatively correlated with the basal subtype (Suppl. Fig. 4D).

### C1QC+ and SPP1+ TAMs comprise two distinct subpopulations within the myeloid cell compartment

Myeloid cells are an important component of the larger TME. We identified twelve distinct myeloid clusters (Fig. 5A). TAMs, representing the majority of myeloid cells, clustered into SPP1+ and C1QC+ subpopulations (Fig. 5A-B). We also identified myeloid-derived suppressor cells (MDSCs), monocytes, and dendritic cells (DCs), which further clustered into conventional DCs 1-3 (cDC1-3) and plasmacytoid DCs (pDCs) (Fig. 5A-B). We did not detect any trends in myeloid composition across samples or in relation to Moffitt subtypes (Suppl. Fig. 5A-B).

**Figure 5.**
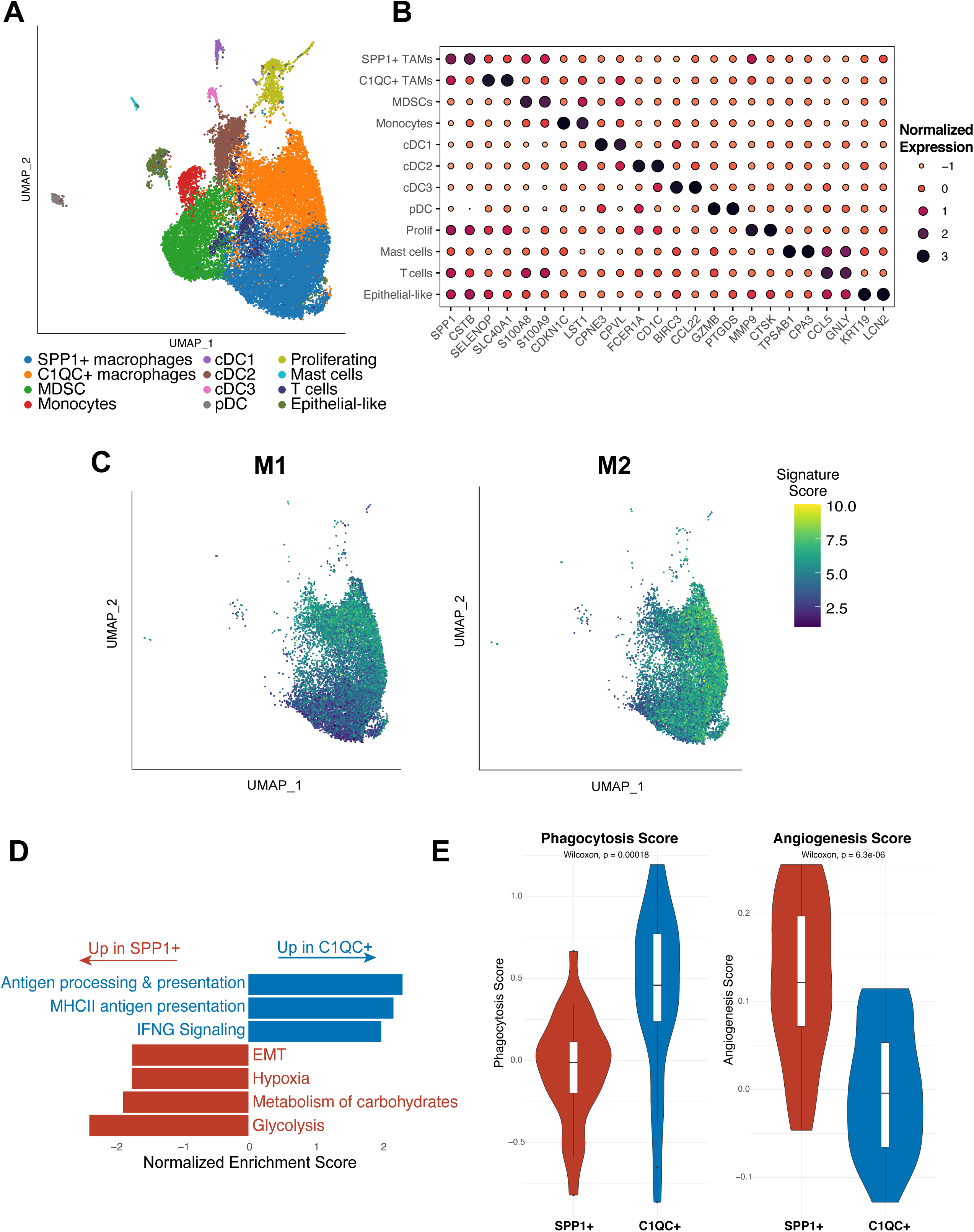
C1QC+ and SPP1+ TAMs comprise two distinct subpopulations within the myeloid cell compartment. A. UMAP overview of the myeloid compartment. DC, dendritic cell; MDSC, myeloid-derived suppressor cell; UMAP, uniform manifold approximation and projection. B. For each myeloid cluster, the two genes most differentially expressed between that cluster and the rest of the myeloid compartment. TAM, tumor-associated macrophages. C. M1 (left) and M2 (right) polarization signatures in the macrophage populations. D. Selected gene set enrichment analysis results from a comparison between SPP1+ and C1QC+ TAMs. EMT, epithelial-mesenchymal transition. E. Phagocytosis and angiogenesis scores for SPP1+ and C1QC+ TAMs. One sample with fewer than 40 total TAM cells was excluded.

To better understand what distinguishes SPP1+ and C1QC+ TAMs, we further characterized these two populations. Historically, TAMs have been classified as either M1 (pro-inflammatory) or M2 (pro-tumorigenic), so we assessed M1 and M2 signature expression in our TAMs^13^. While C1QC+ TAMs showed higher M1 signature expression than SPP1+TAMs, both SPP1+ and C1QC+ subsets displayed strong M2 signature expression and therefore could not be clearly distinguished based on the M1/M2 classification (Fig. 5C). SPP1+ TAMs showed increased expression of S100A8 and S100A9, genes previously linked to immunosuppressive macrophages^31^, while C1QC+ TAMs had increased expression of HLA-DRB5 and CXCL9, which is involved in the anticancer immune response^32^ (Suppl. Fig. 5C). In general, SPP1+ TAMs were enriched for genes involved in EMT, metabolism of carbohydrates, and hypoxia, while C1QC+ TAMs showed enrichment for IFN signaling and antigen presentation (Fig. 5D). Consistent with the gene set enrichment analysis (GSEA) results and a recent colon cancer study^33^, SPP1+ TAMs were enriched for angiogenesis-related genes and C1QC+ TAMs for phagocytosis (Fig. 5E).

To estimate whether the TAM subtype may affect overall survival in PDAC, we evaluated SPP1+ and C1QC+ TAM signature expression in the Cancer Genome Atlas (TCGA) PDAC bulk tumor dataset^9^. While a similar analysis of the colorectal cancer (CRC) TCGA dataset implied worse overall survival for patients with a SPP1+^high^ /C1QC+^low^ TAM combination^33^, we found that estimated TAM composition did not correlate with overall survival in the TCGA PDAC data (Suppl. Fig. 5D).

### Chemotherapy-treatment induces transcriptional changes in cancer cells independent of subtype

To analyze the effect of chemotherapy on the PDAC TME, we focused on primary pancreatic tumor specimens collected before treatment (treatment-naive, n=11) and after treatment (n=6). Specimens collected from liver biopsies were excluded from this part of the analysis in order to minimize potential confounding effects of mixing tumors from multiple sites. As our results have shown that most samples are a mixture of basal and classical cancer cells, which could imply subtype plasticity, we investigated the effect of chemotherapy on subtype signature. We used representative genes for both Moffitt subtypes^7^ to create a basal and classical signature score which allowed us to place each cell on a basal-classical spectrum rather than assigning discrete phenotypes. Overall, treated samples showed no shift towards a more basal-like expression pattern (Fig. 6A), although the treated sample P03 was distinctly more basal than any other sample (Fig. 6B) (P03 was, notably, the only treated sample with poorly differentiated histology). Generally, cancer cells clustered by sample on this subtype spectrum, highlighting intertumoral heterogeneity (Fig. 6B). Looking beyond Moffitt signature genes, we measured the heterogeneity of a sample’s expression profile by calculating the correlations between every pair of its cancer cells. By this metric, treated and naïve samples displayed no difference in intratumor heterogeneity, either per sample (Fig. 6C) or per cell (Fig. 6D).

**Figure 6.**
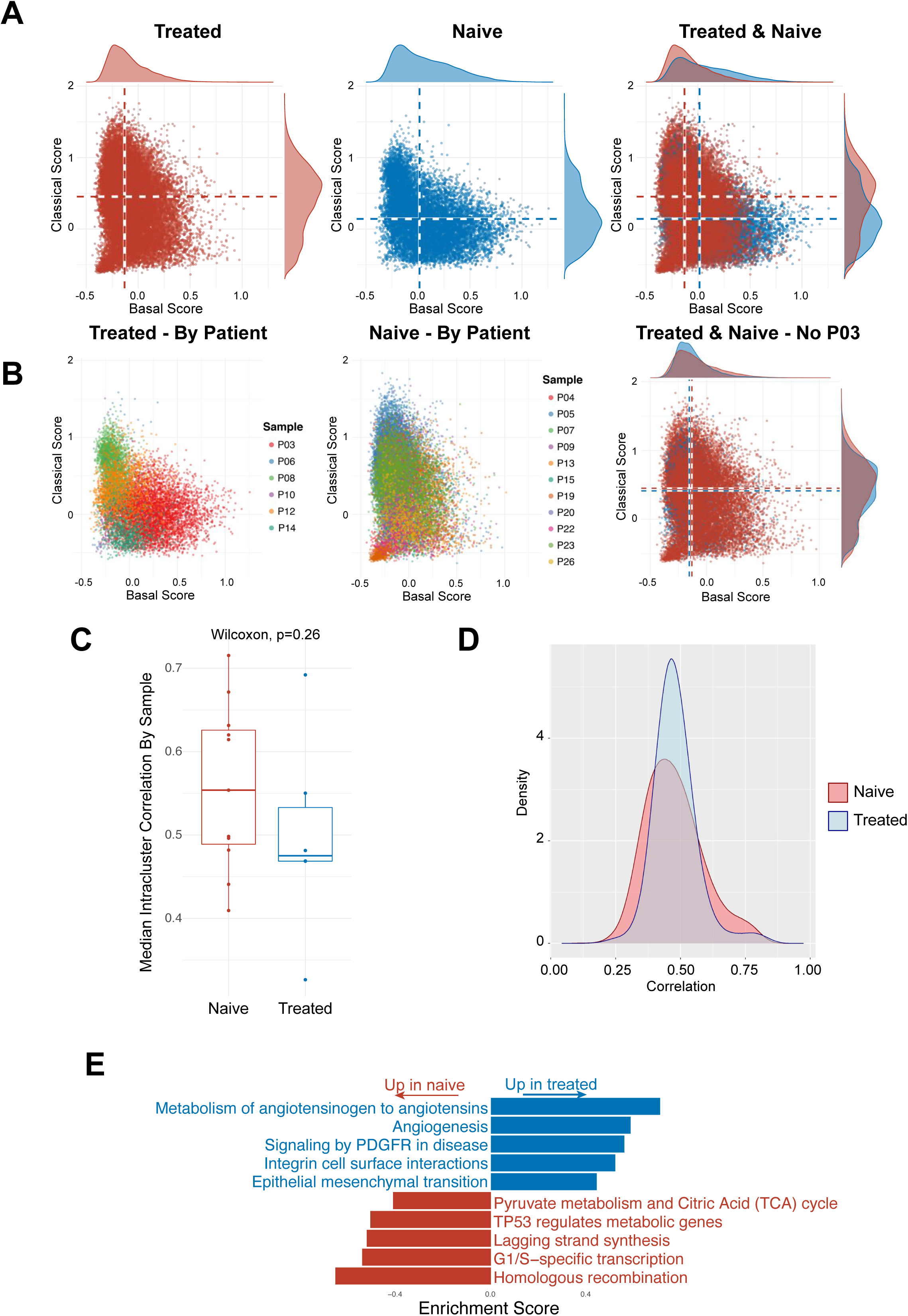
Chemotherapy-treatment induces transcriptional changes in cancer cells independent of subtype. A. Basal and classical signature expression matrix of cancer cells: treated, naive and combined. B. Basal and classical signature expression matrix of cancer cells by sample: treated, naïve, and combined without sample P03. C. Box plot showing the median intracluster correlation of cancer cells by sample for treated and naive groups. D. Density plots of cancer cell correlation within samples for treated and untreated groups. E. Gene set enrichment analysis of overall cancer cells, treated and untreated.

We also used GSEA to check for common gene expression patterns of cancer cells in each sample. Cancer cells from treated samples were enriched in angiogenesis and EMT as compared to naive samples (Fig. 6E), processes that can contribute to tumor recurrence and progression. We did not detect significant changes in CAF and TAM subpopulation distribution between naive and treated groups (Suppl. Fig. 6A).

### Chemotherapy reduces expression of inhibitory checkpoint molecules and ligand-receptor interactions in PDAC

We applied CellPhoneDB^34^ to infer ligand-receptor interactions between malignant epithelial cells, TAM and CAF subpopulations, CD8+ and CD4+ T cells, MDSCs, and DCs, and found a general decrease of cell-cell interactions in the chemotherapy-treated group (Fig. 7A). We noted an unexpectedly high volume of interactions between CAFs and TAMs, which is highlighted in Fig. 7B. As CAFs are known to recruit and polarize macrophages, we checked for recruitment-associated ligand-receptor interactions between CAFs and TAMs^35^. Only two ligand-receptor pairs were detected in this analysis: low levels of the CSF1-CSF1R interaction that did not change with treatment, and an interaction between CXCL12 (on iCAFs but not myCAFs) and CXCR4 (on both TAM subpopulations). CXCL12-CXCR4, the strongest ligand-receptor interaction pair in naive samples, significantly weakened with treatment (Fig. 7C). The CXCL12-CXCR4 interaction between perivascular CAFs and TAMs has recently been linked to metastasis^36^ and may be one way in which conventional chemotherapy affects crosstalk in the TME.

**Figure 7.**
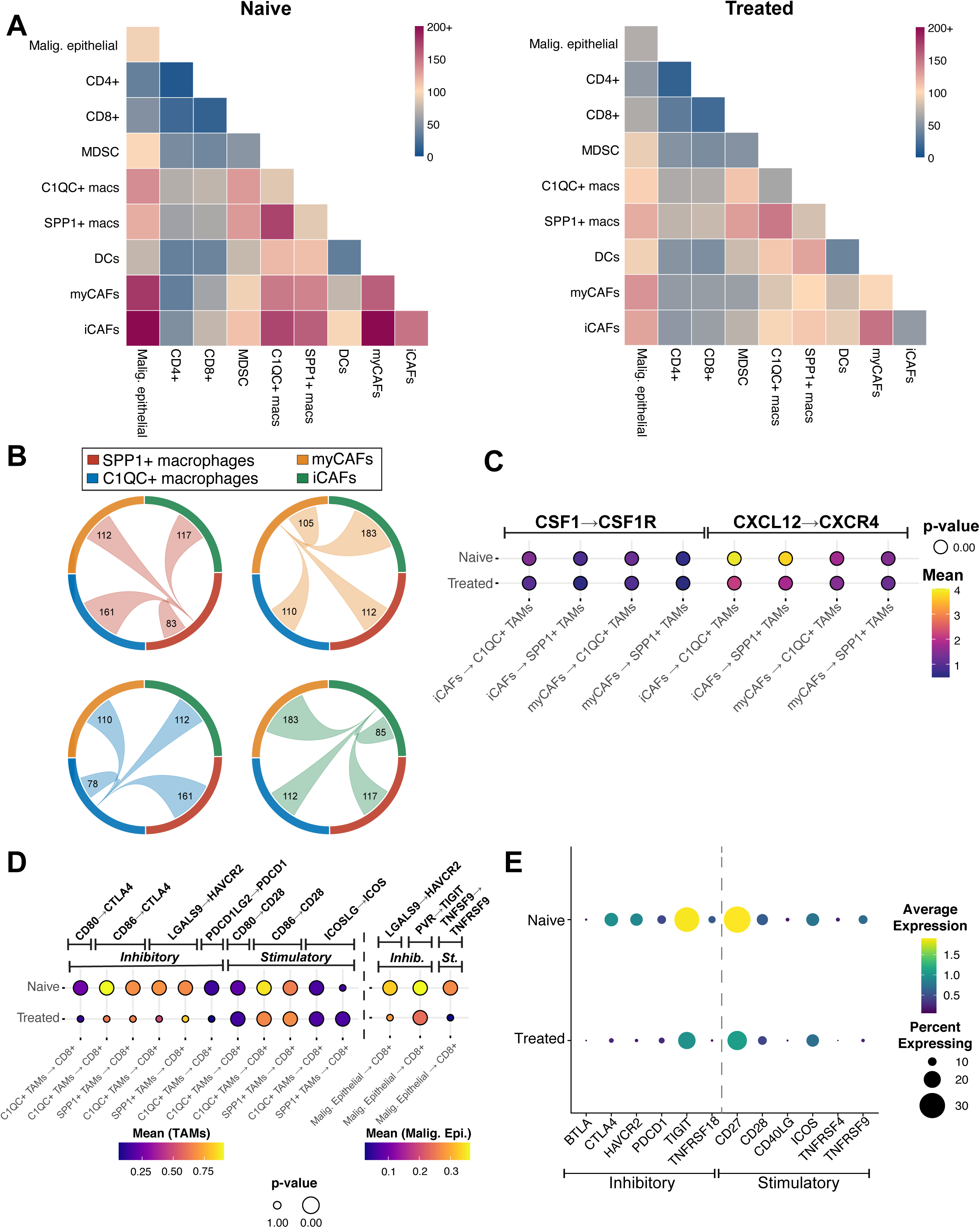
Chemotherapy reduces expression of inhibitory checkpoint molecules and ligand-receptor interactions in PDAC. A. Heatmap displaying an overview of all inferred LRIs between displayed cell types (cancer cells, MDSCs, DCs, C1QC+ TAMs, SPP1+ TAMs, iCAFs, myCAFs, CD8+ T cells, CD4+ T cells). Left: untreated samples, right: treated samples. CAF, cancer-associated fibroblast; DC, dendritic cell; iCAF, inflammatory CAF; LRI, ligand-receptor interaction; MDSC, myeloid-derived suppressor cell; myCAF, myofibroblastic CAF; TAM, tumor-associated macrophage. B. Interaction network for SPP1+ and C1QC+ TAMs and iCAFs and myCAFs. C. Dot plot of CellphoneDB output for recruitment-associated LRIs between iCAFs/myCAFs and SPP1+/C1QC+ TAMs comparing untreated and treated samples. D. Dot plot of CellphoneDB output for checkpoint molecule LRIs between CD8+ T cells, SPP1+ or C1QC+ TAMs, and cancer cells comparing untreated and treated samples. E. Differential gene expression of checkpoint molecules in CD8+ T cells between treated and untreated samples.

Since checkpoint inhibitory drugs have yet to show benefit in PDAC clinical trials (except in the 1% of PDAC cases with mutations in mismatch repair genes^37^), we assessed whether treatment alters checkpoint molecule interactions between CD8+ T cells, TAMs, and cancer cells. We evaluated all CD8+ T cells due to the scarcity of exhausted T cells in our dataset. Moreover, CD8+ T cells can exhibit different states of exhaustion, and the larger CD8+ population is more representative of response regarding checkpoint molecule interactions with chemotherapy^38^. We observed more ligand receptor interactions between TAMs and CD8+ T cells than between cancer cells and CD8+ T cells (Fig. 7D). Comparing treated and naive samples, we noticed that all of the observed inhibitory ligand receptor interactions were only significant in naive samples, while immune stimulatory interactions were significant in both treated and naive samples between TAMs and CD8+ T cells (Fig. 7D). The only significant PD-1 (PDCD-1) interaction occurred in naive samples, between PD-1 on C1QC+ TAMs and PD-L2 on CD8+ T cells (Fig. 7D). We detected only three significant interactions between CD8+ T cells and cancer cells, with TIGIT-PVR being the strongest. TIGIT-PVR is the only one that was significant in naive and treated samples, though the effect was weakened in treated samples (Fig. 7D). Of note, we did not detect PD-1 related interactions between CD8+ T cells and cancer cells.

Given the clinical relevance of these findings, we further evaluated checkpoint molecule expression in CD8+ T cells. We found TIGIT to be the highest and most broadly expressed inhibitory checkpoint molecule, while PD-1 was expressed at low levels and in a minority of CD8+ T cells (Fig. 7E). Expression of TIGIT, PD-1, and other checkpoint molecules was lower in CD8+ T cells of chemotherapy-treated samples than naive samples (Fig. 7E).

## Discussion

In this study we systematically characterized the composition of human PDAC at the single-cell transcriptomic level and assessed the effects of chemotherapy on the TME. Individual samples demonstrated significant heterogeneity in Moffitt subtype and TME composition. Analysis of the TME revealed discrete cell subpopulations with distinct features, further highlighting the complexity of the PDAC TME. We found that chemotherapy profoundly alters the PDAC TME and might lead to further resistance to immunotherapy by reduced inhibitory checkpoint molecule expression and interactions involving CD8+ T cells.

Consistent with previous reports on subtype heterogeneity, our initial per-sample analysis of cancer cell composition revealed a heterogeneous mixture of cells with classical or basal transcriptional signatures in most samples and a variety of subclonal growth patterns^22, 23, 39, 40^. A recent study demonstrating plasticity of pancreatic cancer cells and subtype switching dependent on culture conditions *in vitro* also identified a new intermediate subtype state^23^. Although we did not observe their specific intermediate state gene signature (the number of samples in each dataset may limit the generalizability of such a signature), we also discovered cancer cells that expressed both basal and classical histological markers to varying degrees. Since both subtypes exhibit different biology and metabolism^41, 42^, using scSeq to evaluate individual subtype composition prior to therapy might help understand how these elements dictate responses to different therapeutic approaches.

In recent years, investigation of CAFs has led to the identification of distinct CAF subpopulations in human and mouse PDAC^19^. While iCAFs and myCAFs were present in our dataset, neither unsupervised clustering nor signature analysis revealed a defined apCAF population. Nevertheless, a selection of iCAFs and myCAFs did co-express CD74 and HLA genes, consistent with another recent report^22^. Plasticity between CAF subtypes, as hypothesized by multiple groups, might explain the differences in our CAF subpopulation findings^19, 24^.

Myeloid cell analysis led to the identification of SPP1+ and C1QC+ TAMs as the major TAM subpopulations in human PDAC. In our data, M1/M2 polarization did not fully describe these recently described subpopulations, suggesting that M1/M2 classification does not capture the complexity of TAM subpopulations in the human TME^43–45^. Further characterization showed enrichment for EMT and a high angiogenesis score in SPP1+ TAMs, while C1QC+ TAMs were enriched for antigen presentation and phagocytosis. These findings are consistent with recent scRNA-seq studies in CRC^33, 46^. A SPP1+^high^/C1QC+^low^ TAM signature has been associated with worse overall survival in the TCGA CRC dataset, and SPP1+ TAMs have been linked to CRC liver metastases^33, 47^. In our analysis of both TAM signatures in the TCGA PDAC dataset, TAM subtype combination did not correlate with worse overall survival. Since the TCGA PDAC dataset has only bulk transcriptomics and has a limited variety of clinical stages, the role of SPP1+ and C1QC+ TAMs in PDAC requires further evaluation in a larger, non-bulk RNA clinical cohort.

For a more unambiguous analysis of chemotherapy effects on the TME we compared naive and treated samples from primary tumors only. Recent studies have correlated the classical subtype with better therapy response than the more aggressive basal subtype and proposed that chemotherapy might shift the Moffitt subtype from classical to basal^48, 49^. While there was heterogeneity between tumors in regard to Moffitt subtype composition, they did not shift toward a basal subtype in treated samples or display significant differences in intratumor heterogeneity. However, there was a Moffitt-independent trend toward enrichment of angiogenesis and EMT in treated samples. EMT is a hallmark feature of tumor invasion, and different EMT programs have been linked to PDAC progression, dissemination, and resistance to chemotherapy^50–52^. While the complexity of EMT makes therapeutic approaches targeting it challenging, novel approaches would be highly impactful^53^. Tumor angiogenesis and microvessel density have been linked to metastasis in PDAC^54–56;^ while anti-angiogenic drugs have thus far not yielded clinical results, the question of how chemotherapy affects the microvasculature in PDAC warrants further investigation^57^.

The complexity of PDAC is not only determined by the cancer cell heterogeneity but also the intricate composition of and communication between cells in the TME. In our analysis, we found a high volume of ligand receptor interactions between TAMs and CAFs. There is limited data on crosstalk between CAFs and TAMs in the TME, even though these interactions likely play an important role in PDAC biology and may reveal potential targets for therapy^35^. We found CXCL12-CXCR4, specific to iCAFs and TAMs respectively, to be the most significant interaction. Notably, it has been demonstrated that CXCL12 secretion by CAFs in PDAC promotes immune evasion by T cell exclusion^58^, and CXCL12-CXCR4 interaction between perivascular CAFs and TAMs promotes metastasis in a breast cancer model^36^. In our dataset, CXCL12-CXCR4 interaction between iCAFs and TAMs was significantly lower in treated samples, exemplifying how conventional chemotherapy may alter the PDAC TME in ways we could not previously observe.

Although effective in a number of other malignancies, checkpoint inhibition therapy has performed poorly in PDAC. While the underlying biology is still being uncovered, few studies have explored checkpoint expression in human PDAC samples at the single cell level. In our analysis, exhausted CD8+ T cells constituted only 2.8% of all CD8+ T cells. This corresponds with previous studies showing PDAC to be a relatively immune-cold tumor with low PD-1 expression and might be a reason why checkpoint inhibition has failed in clinical trials for PDAC^4, 5, 59^. The strongest interaction we observed between CD8+ T cells and cancer cells was TIGIT-PVR, which significantly decreased with treatment. Recently, Steele et al. showed elevated TIGIT expression in tumor-infiltrating CD8+ T cells while PD-1 and LAG-3 were not significantly altered in human PDAC^21^. Functionally, Freed-Pastor et al. used a PDAC mouse model to show that the TIGIT-PVR axis is critical for maintaining immune evasion and that TIGIT blockage elicited an immune response to PDAC^60^. TIGIT was the most expressed inhibitory checkpoint molecule on CD8+ T cells and its expression, as with other checkpoint molecules, decreased with treatment. These findings imply that TIGIT may be a stronger immunotherapy candidate than PD-1 in PDAC. The higher TIGIT expression in treatment-naive samples further suggests that blocking TIGIT may be more effective given as first-line monotherapy or concurrently with chemotherapy, and that immune blockade as second-line treatment following chemotherapy may lessen its antitumor effects. While a recent phase III trial of tiragolumab as first line therapy in small cell lung cancer was unsuccessful^61^, the Morpheus-Pancreatic Cancer trial (NCT03193190) is investigating the effect of TIGIT blockage in metastatic PDAC and will hopefully provide more insight into the clinical relevance of TIGIT inhibition in PDAC. Our results provide an in-depth view into human PDAC composition, its TME, and the effects of chemotherapy; we believe these observations will be valuable bases for further clinical and biological studies of PDAC.

## Methods

### Patient cohort and data collection

A total of 27 PDAC patients were recruited for study participation at the Perlmutter Cancer Center at New York University (NYU) Langone Health between May 1, 2020, and June 30, 2021. Informed consent for the collection of blood, tissue, and clinical information was obtained from all patients using an institutional review board-approved study protocol. Standard clinicopathological variables including sex, age, tumor site, disease stage, type of procedure, tumor stage (TNM classification), and treatment type were collected for each patient as part of our prospective PDAC database. Clinical germline and somatic tumor analyses (Tempus xT 596 gene panel, Tempus Labs, Chicago, IL; Invitae 91 gene panel, Invitae Corporation, San Francisco, CA; FoundationOne CDx 324 gene panel, Foundation Medicine, Cambridge, MA) were performed. A supplement providing individual UMAP, H&E as well as clinical and mutational information is provided in the Extended Data.

### Sample acquisition and processing

Of the 27 samples, 10 were obtained following surgical resection, 7 were obtained from endoscopic ultrasound (EUS)-guided biopsies, and 10 were obtained by interventional radiology (IR) from metastatic liver lesions. For EUS- and IR-guided biopsies, 2-3 cores were used for scRNA-seq analysis. The pathologic diagnosis of PDAC was confirmed in all samples by a board-certified pancreatic pathologist. Within 4-6 h of tissue acquisition, the specimen was placed in ice-cold RPMI 1640 medium (Gibco, Cat. No. 11875101) and transported to the Center for Biospecimen Research & Development at NYU, where the sample was provided for tissue processing.

### Tissue processing and single cell isolation

Tissue samples were washed in ice-cold phosphate-buffered saline (PBS; Corning, Cat. No. 21-040-CM) and grossly assessed for adequacy (solid tissue component, free of large hemorrhagic or necrotic areas). Next, the tissue was manually minced with a scalpel blade and filtered through a 40 µm mesh filter (Falcon; Cat. No. 352340) to remove any excess debris or blood. The tissue was then transferred to a sterile gentleMACS C tube (Miltenyi Biotec, Cat. No. 130096334), resuspended in 3-5 mL of Miltenyi Tumor Dissociation Kit buffer (Cat. No. 130-095-929), and mechanically minced to submillimeter particles. The suspension was then digested utilizing the gentleMACS Octo Dissociator (Miltenyi Biotec, Cat. No. 130-095-937) for 25-30 min using the “37C_h_TDK_3’’ program for tumor tissue. Dissociated cells were then quenched with ice-cold RPMI 1640 containing 10% fetal bovine serum (FBS; Gibco, Cat. No. 10082147) and filtered through a 40 µm mesh filter. Red blood cells were removed by treatment with 3-5 mL of ACK lysis buffer (Gibco, Cat. No. A1049201) and a subsequent washing step in RPMI 1640 with 10% FBS. Final cell counts were established with 0.4% trypan blue solution and a Countess II Automated Cell Counter (Applied Biosystems, Cat. No. A27977). The final concentration was titrated to 300-500 viable cells/µL for further analysis.

### Single-cell library preparation

Single-cell suspensions were processed for 10x Genomics by the Genome Technology Center at the NYU School of Medicine per the manufacturer’s guidelines. The libraries were prepared using the following reagents: Chromium Single Cell 3′ Reagent Kits (v3)—the Single Cell 3′ Library & Gel Bead Kit v3 (PN-1000075), the Chromium Next GEM Chip G Single Cell Kit (PN-100120), and the i7 Multiplex Kit (PN-120262) (10x Genomics) —by following the Single-Cell 3′ Reagent Kits (v2) User Guide (manual part no. CG00052 Rev A) and the Single-Cell 3′ Reagent Kits (v3) User Guide (manual part no. CG000183 Rev C). DNA libraries were run using the NovaSeq 6000 system (Illumina, San Diego, CA) to achieve roughly 500 million paired-end reads per sample.

### Single-cell RNA sequencing and data processing

Sequencing results were de-multiplexed and converted to FASTQ format using Illumina bcl2fastq software. The 10x Genomics Cell Ranger 5.0.1 software suite^62^ was used to perform sample de-multiplexing, barcode processing, and single-cell 3’ gene counting. The cDNA insert was aligned to the hg38/GRCh38 reference genome. Only confidently mapped, non-PCR duplicates with valid barcodes and unique molecular identifiers were used to generate the gene-barcode matrix. Further analysis including the identification of highly variable genes, dimensionality reduction, standard unsupervised clustering algorithms, and the discovery of differentially expressed genes was performed using Seurat^63^ and scooter^64^.

Raw data were initially filtered to remove ambient RNA using SoupX version 1.5.2^65^; and cells were subsequently filtered to include only high-quality cells (as defined by >500 detectable genes, >1500 unique molecular identifiers, and <15% of transcripts coming from mitochondrial genes). Cells with >1% of transcripts representing erythroid genes (HBA1, HBA2, HBB, HBM, and ALAS2) were also excluded from the analysis. The data were normalized to the total expression, multiplied by a scaling factor of 10,000, and log-transformed. Likely doublets/multiplets were identified and removed using scDblFinder version 1.6.0^66^.

To account for biological and technical batch differences between individual patients and scRNA-seq libraries, the Seurat anchor-based integration method for merging datasets that identify pairwise correspondences between cell pairs across datasets^67^ was utilized to transform them into a shared space. The 2,000 most variable genes based on standardized variance were selected for canonical correlation analysis as an initial dimensional reduction. The integration anchors were then identified based on the first 30 dimensions and used to generate a new dimensional reduction for further analysis.

To visualize the data, the dimensionality of the scaled integrated data matrix was further reduced to project the cells in two-dimensional space using principal component analysis followed by UMAP^68^. The 30 nearest neighbors were used to define the local neighborhood size with a minimum distance of 0.3. The resulting principal components were also used as a basis for partitioning the dataset into clusters using a smart local moving community detection algorithm^69^. A range of resolutions (0.1-10) was utilized to establish a sufficient number of clusters to separate known populations based on the expression of established markers.

### Cell identification and clustering

Single-cell clusters were identified based on common marker genes for various cell types, the most prominent of which are listed here. Full dataset (Figure 1C): T/NK (CD3E), epithelial (KRT19), endothelial (VWF, PECAM1), myeloid (CD68), proliferating epithelial (KRT19, MKI67), proliferating lymphoid (CD3E, MKI67), B/plasma (CD79A), proliferating myeloid (CD68, MKI67), mesenchyme (DCN), mast (KIT). Mesenchyme (Figure 3A): myCAF (ACTA2, MMP11, COL10A1^-^), iCAF (C3, C7, CFD, PTGDS), pericyte (ACTA2^+^, RGS5^+^, CSPG4, NOTCH3), proliferating (MKI67), peri-islet Schwann cells (SOX10, S100B, PLP1), epithelial-like (KRT19, EPCAM), chondrocyte-like (SPP1, IBSP, MMP13). T/NK (Figure 4A): CCR7+ CD4+ T cells (CCR7, CD4), IL7R+ CD4+ T cells (IL7R, CD4), FOXP3+ CD4+ T cells (FOXP3, CD4), CXCL13+ CD4+ T cells (CXCL13, CD4), GZMH+ CD8+ T cells (GZMH, CD8), GZMK+ CD8+ T cells (GZMK, CD8), ITGA1+ CD8+ T cells (ITGA1, CD8), ISG15+ T cells (ISG15), GNLY+ NK cells (GNLY, NCAM1), XCL1+ NK cells (XCL1, NCAM1), KIT+ mast (KIT), CD79+ B/Plasma (CD79A), MKI67+ proliferating (MKI67). CD8+ T cells (Figure 4C): GZMK+ CD8+ T cells (GZMK), GZMH+ CD8+ T cell (GZMH), ITGA1+ CD8+ T cells (ITGA1), SELL+ CD8+ T cells (SELL), ISG15+ CD8+ T cells (ISG15), CTLA4+ CD8+ T cells (CTLA4), HSPA6+ CD8+ T cells (HSPA6), epithelial-like (KRT19), erythrocytes (HBB). Myeloid (Figure 5): SPP1+ macrophage (SPP1, MARCO), C1QC+ macrophage (C1QA, C1QB, C1QC), MDSC (S100A8, S100A9, S100A12), monocyte (FCGR3A, CDKN1C), cDC1 (CLEC9A, XCR1), cDC2 (CD1C, FCER1A), cDC3 (CCL19, CCL22, CCR7), pDC (LILRA4, PLD4), proliferating (MKI67), mast (KIT), T (CD3E), epithelial-like (KRT18, KRT19).

### Identification of malignant and pancreatic epithelial cells

To detect large-scale chromosomal CNVs, InferCNV version 1.8.1^70^ was run at a sample level with the parameters recommended for 10x Genomics data. For each sample, all epithelial cells were tested for copy number aberrations. Non-epithelial (CAF, endothelial, immune) cells were used as the reference normal set. For samples with over 1,000 non-epithelial cells, they were randomly subsampled to 1,000. Additionally, 10% of non-epithelial cells were used in the observation set. We used a 101-gene window in subclustering mode with HMM enabled and a six-group clustering of observations.

### Ligand receptor interaction

CellPhoneDB v.2.1.7^34^ was used to estimate ligand-receptor interaction. We used the statistical method with 10 iterations and no subsampling; other parameters were left as default.

### Gene signature scores

Gene signature scores were created using Seurat’s AddModuleScore function. Lists of genes used for each score can be found in Supplementary Table 4.

### Differential gene expression

Differential gene expression was analyzed using the Wilcoxon rank-sum test with Bonferroni correction that is included in Seurat’s FindMarkers function.

### Gene set enrichment analysis

Gene set enrichment analysis was performed using the GSEAPreranked function of GSEA version 4.2.0^71^. Genes were pre-ranked for each comparison using the output of our differential gene expression; first by taking the -log_10_(adjusted p-value) and then multiplying it by the sign of the fold change (for example, a gene with a fold change of -2.3 and an adjusted p-value of 1e-160 would have a rank metric of -160). GSEAPreranked parameters were set to the defaults. The gene chip annotation file used was Human_Gene_Symbol_with_Remapping_MSigDB.v7.4.chip, and genes were collapsed and remapped to gene symbols. Gene sets used included Hallmarks (h.all.v7.5.1), KEGG (c2.cp.kegg.v7.5.1), Reactome (c2.cp.reactome.v7.5.1), Canonical Pathways (c2.cp.v7.5.1), and GO (c5.go.v7.5.1).

### Survival analysis of TCGA data

The TCGA PDAC (PAAD) clinical and gene expression data^9^ were downloaded using the TCGABiolinks version 3.15^72^ to examine the prognostic utility of specific cell subpopulation signatures. Gene expression was z-score transformed. For each signature, two distinct groups (low and high expression) were formed, corresponding to mean expression values below the 45th and above the 55th quantile values, respectively. The survival analysis was performed using the R package survminer^73^. Cox proportional hazards models from the R package survival^74^ were used to adjust for tumor stage, age, and sex.

### Histology and multiplex IHC

For histopathological assessment, tumor sections were cut and slides prepared by routine processes, followed by staining with hematoxylin & eosin. For multiplex immunofluorescence and imaging, five-micron formalin fixed paraffin-embedded sections were stained with Akoya Biosciences® Opal™ multiplex automation kit reagents (Leica Cat #ARD1001EA) on a Leica BondRX® autostainer, according to the manufacturers’ instructions. In brief, slides underwent sequential epitope retrieval with Leica Biosystems Epitope Retrieval 2 solution (ER2, EDTA based, pH 9, Cat. AR9640), primary and secondary antibody incubation, and tyramide signal amplification with Opal® fluorophores as shown in Supplementary Table 3. Primary and secondary antibodies/polymers were removed during sequential epitope retrieval steps while the relevant fluorophores remained covalently attached to the antigen. Slides were counterstained with spectral DAPI (Akoya Biosciences, FP1490) and mounted with ProLong Gold Antifade (ThermoFisher Scientific, P36935). Semi-automated image acquisition was performed on a Vectra® Polaris multispectral imaging system. After whole slide scanning at 20x in the motif mode, regions of interest were selected for spectral unmixing and image processing using InForm® version 2.4.10 software from Akoya Biosciences.

## Data availability

scSeq data is currently being uploaded to dbGaP.

## Acknowledgments

We thank Valeria Mezzano Robinson, Shanmugapriya Selvaraj and Branka Brukner Dabovic from the Experimental Pathology Core at NYU Langone Health for support with multiplex IHC staining and imaging. We thank Peter Meyn and Yutong Zhang from the Genome Technology Center Core at NYU Langone Health. We thank the Center for Biospecimen Research and Development at NYU Langone Health for support with tissue acquisition. These shared resources are partially supported by the Cancer Center Support Grant P30CA016087 at the Laura and Isaac Perlmutter Cancer Center.

This project was supported by NIH R01CA245005 (DMS) and a research scholar grant from the German Research Foundation (WE7047/1-1; DW).

## Author contributions

G.W., D.W., E.A.K., I.D., A.S. and D.M.S. developed the study concept and were responsible for the study design. D.W., E.A.K. and G.W. wrote the manuscript, which was then edited by all co-authors. D.M.S., T.G., X.J. and C.H. helped acquire human biospecimens. G.W., D.W., E.Z., G.O. and A.H. processed samples for scSeq. D.W., E.A.K., G.W., I.D. and D.K. analyzed the scRNA-Seq data. E.A.K., I.D. and D.K. were responsible for the bioinformatics analysis pipelines. X.J., G.W., E.Z. and D.W. collected patient information and maintained the databases. C.H., C.L., D.W., G.W. and D.M.S. analyzed the H&E and multiplex IHC. S.D., T.G., N.B., L.K. and P.O. provided study guidance and feedback on the manuscript. M.H.S., A.W.L. and T.H.W. assisted with study design and analysis. D.M.S. and A.S. directed the study. All co-authors approved the final version of the manuscript before submission.

---

**Supplementary Figure 1.**
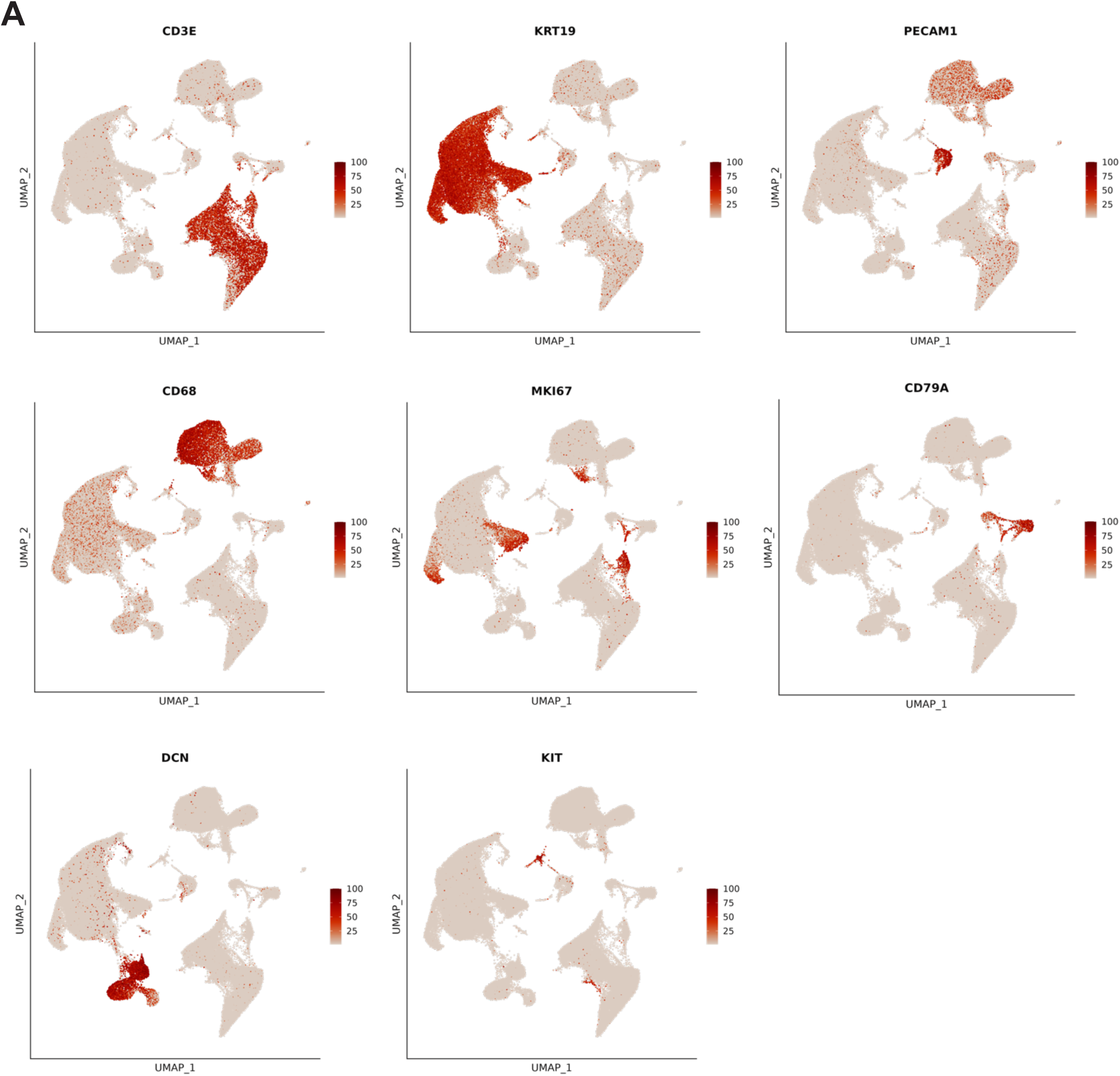
A. UMAP embeddings of canonical marker gene expression, utilized in the identification of clusters. UMAP, uniform manifold approximation and projection.

**Supplementary Figure 2.**
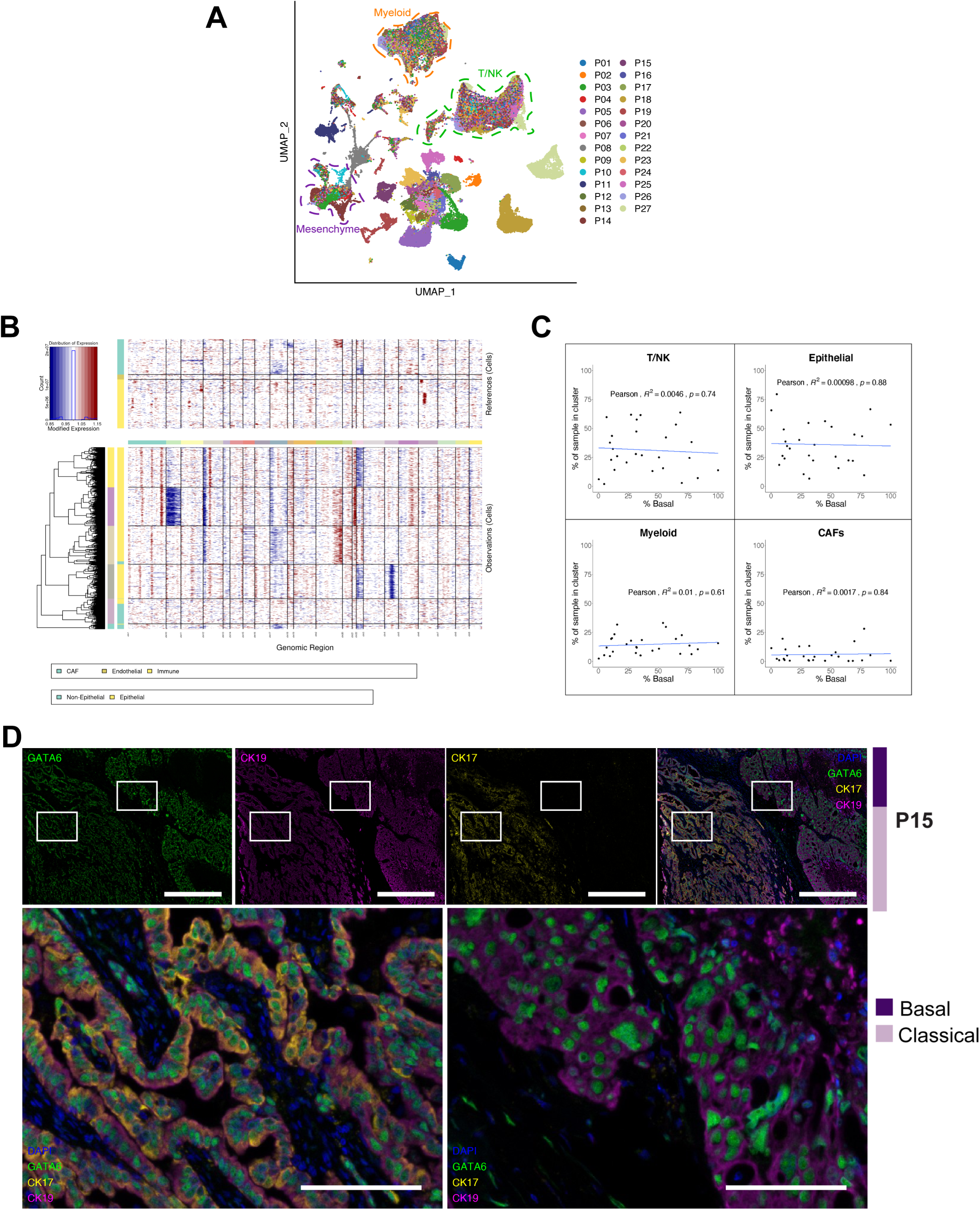
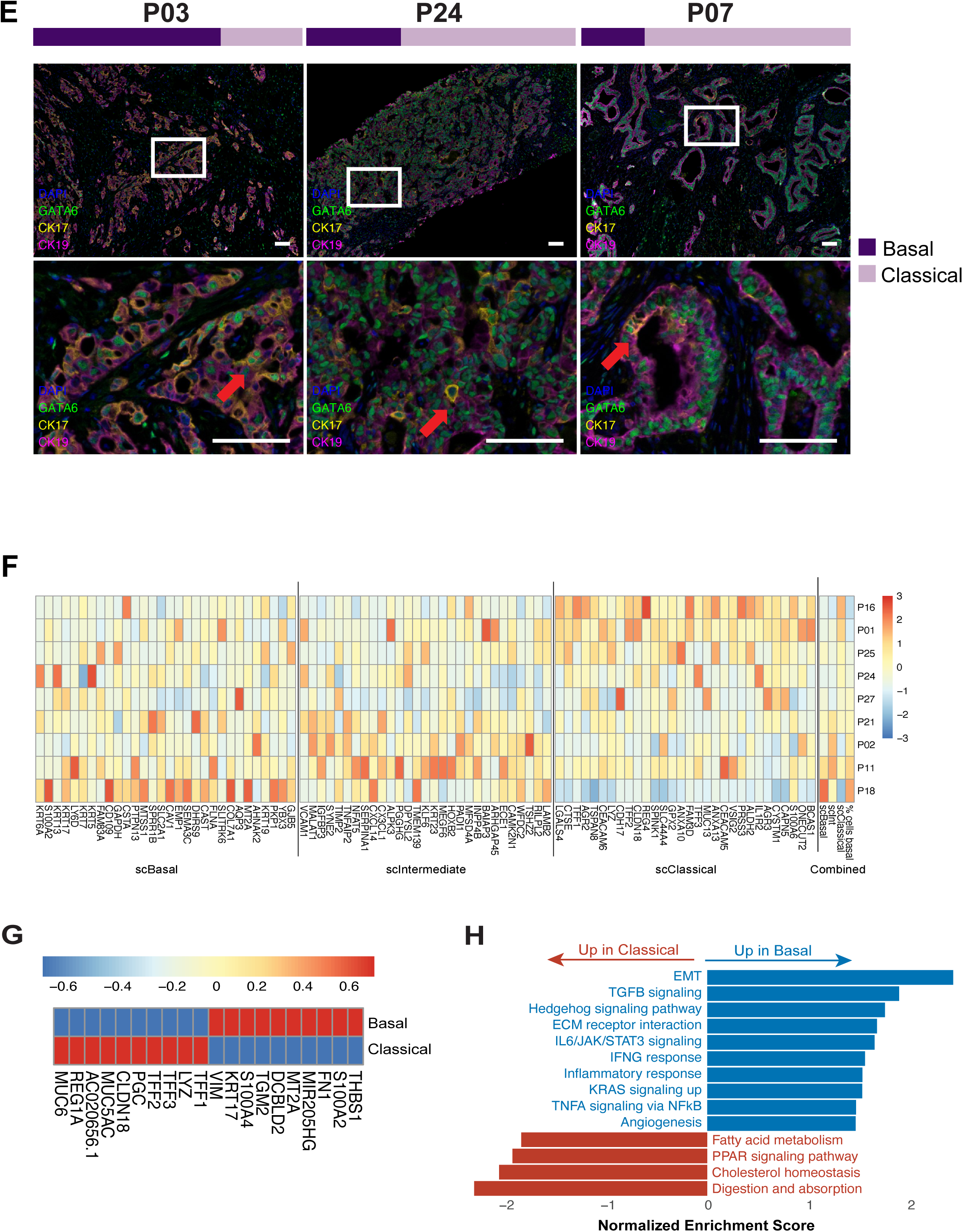
A. Non-batch corrected UMAP embedding reveals distinct transcriptomic landscapes in patient-specific tumor compartments, while cells of other lineages evince significant overlap. Clusters with non-epithelial cells are labeled with the majority cell type. UMAP, uniform manifold approximation and projection. B. inferCNV output for a patient sample (P19) with more than one subclone indicated in the CNV analysis. C. Correlation between proportion of malignant epithelial cells in a sample that are labeled as basal and the percentage of the sample that is in each of four major clusters. D. Representative images of multiplex immunofluorescence from a case with two distinct growth patterns (P15). Basal to classical ratio by scRNA-seq transcriptional analysis is shown on the right as colored bars (dark = basal, light = classical). Channels (always including DAPI (blue)): GATA6 (green), CK19 (cytokeratin 19) (violet), CK17 (cytokeratin 17) (yellow), and merged. Scale bar = 500 μm. Corresponding high power merged images of two distinct regions are indicated in the low power image by white boxes, highlighting the different growth pattern and marker expression (scale bar = 100 μm). E. Multiplex immunofluorescence images corresponding to main Fig. 2E. For each case, the merged low power image is shown (scale bar = 100 μm), along with the corresponding high power image of the region indicated in a white box in the low power image (scale bar = 100 μm). Red arrows indicate cells that express both GATA6 and CK17. F. Reproduction of the scBasal, scIntermediate, and scClassical subtypes from Raghavan *et al.* in our untreated liver samples (n=9). Samples are sorted from low to high by the percentage of cells determined to be basal using the Moffitt signature. G. The top ten differentially expressed genes between the basal and classical malignant epithelial cells. H. Selected gene set enrichment analysis results from a comparison between the classical and basal cancer cell subtypes.

**Supplementary Figure 3.**
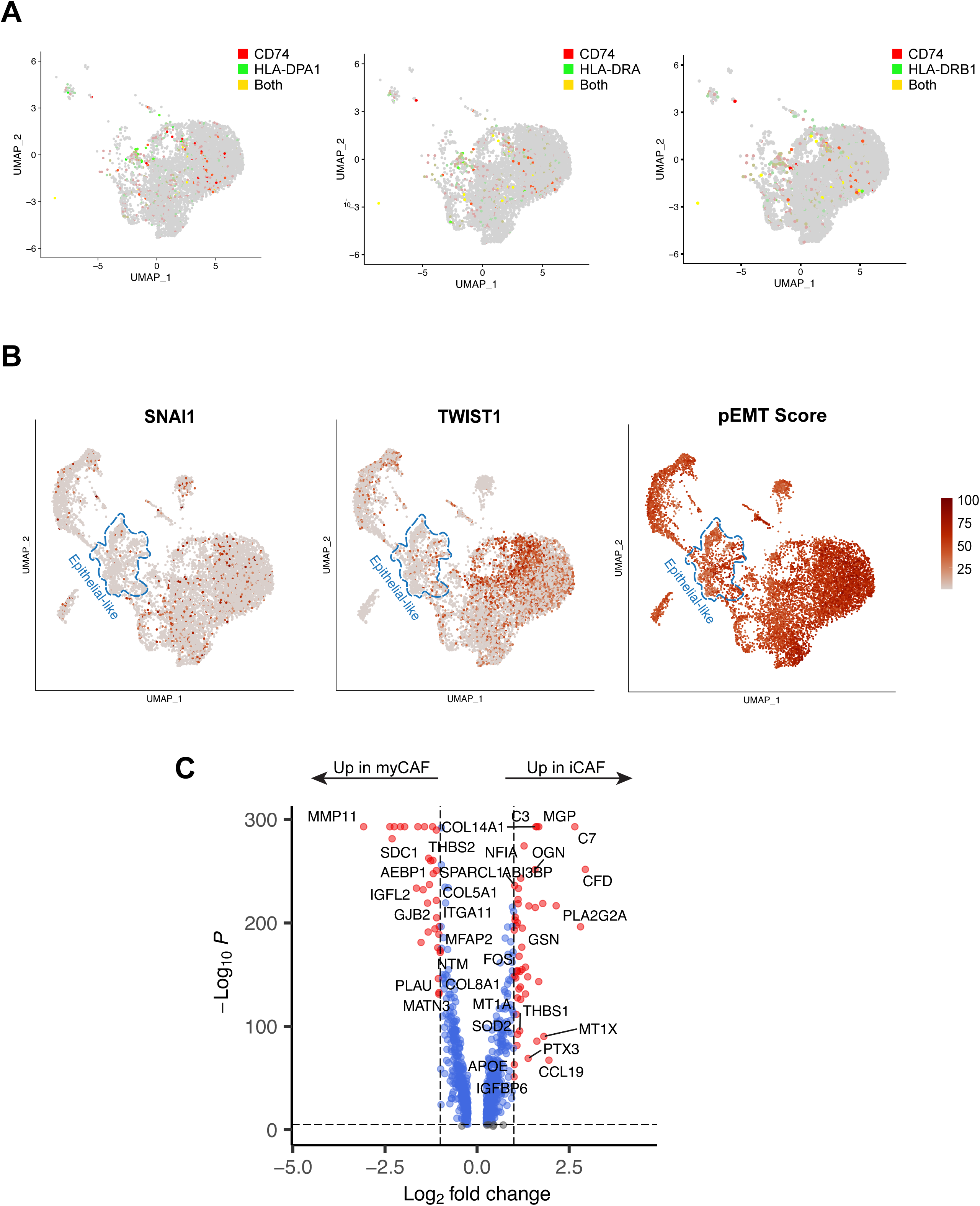
A. UMAP of CD74 and HLA-DPA1, HLA-DRA, and HLA-DRB1 co-expression in CAFs. UMAP, uniform manifold approximation and projection. B. Canonical EMT markers (SNAI1 and TWIST1) and pEMT score in the mesenchymal compartment suggest that the epithelial-like cluster does not consist of cells undergoing EMT. EMT, epithelial-mesenchymal transition. C. Differentially expressed genes between our iCAFs and myCAFs. CAF, cancer-associated fibroblast; iCAF, inflammatory CAF; myCAF, myofibroblastic CAF.

**Supplementary Figure 4.**
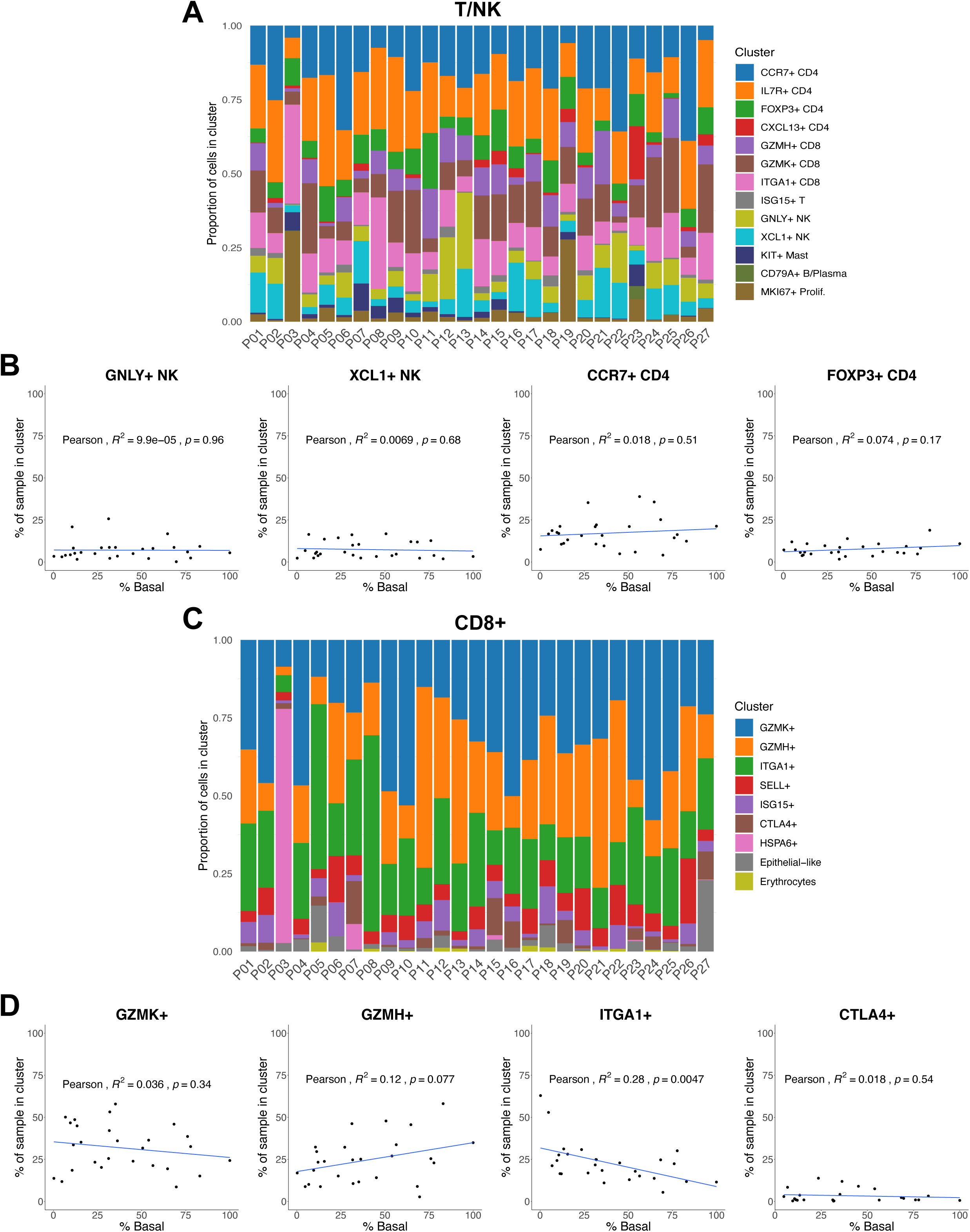
A. Proportional distribution of each T/NK cell cluster by individual sample. NK, natural killer. B. Correlation between proportion of malignant epithelial cells in a sample that are labeled as basal and the percentage of the sample that is in each of four exemplary T/NK clusters. C. Proportional distribution of each CD8+ T cell cluster by individual sample. D. Correlation between proportion of malignant epithelial cells in a sample that are labeled as basal and the percentage of the sample that is in each of four exemplary CD8+ T cell clusters.

**Supplementary Figure 5.**
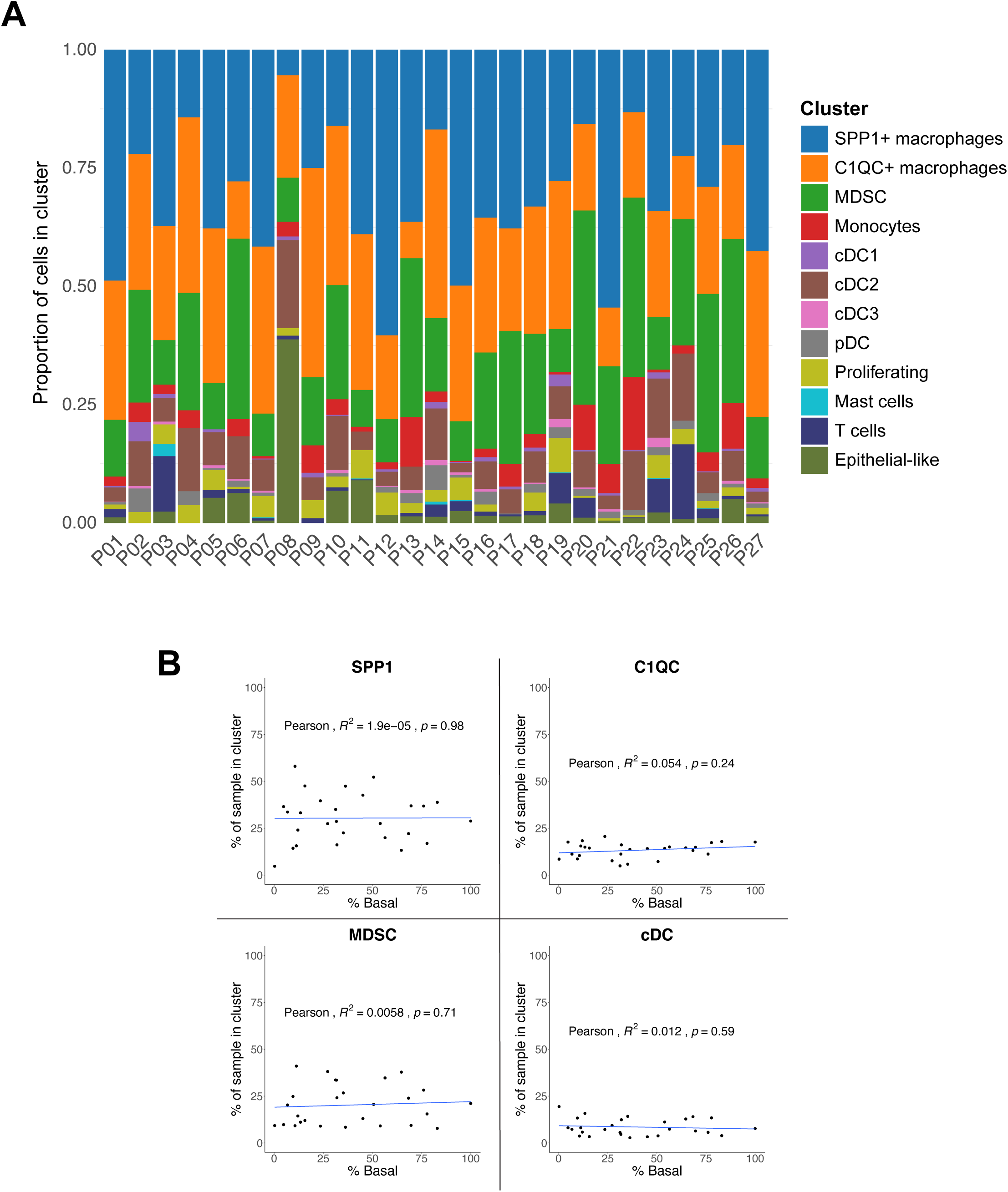
A. Proportional distribution of each myeloid cluster in each individual patient. DC, dendritic cell; MDSC, myeloid-derived suppressor cell. B. Correlation between the proportion of malignant epithelial cells in a sample that are labeled as basal and the percentage of the sample that is in each of the four exemplary myeloid populations. C. Differentially expressed genes between SPP1+ and C1QC+ TAMs. D. PDAC TCGA data on overall survival of patients with different combinations of C1QC+ and SPP1+ TAM signatures.

**Supplementary Figure 6.**
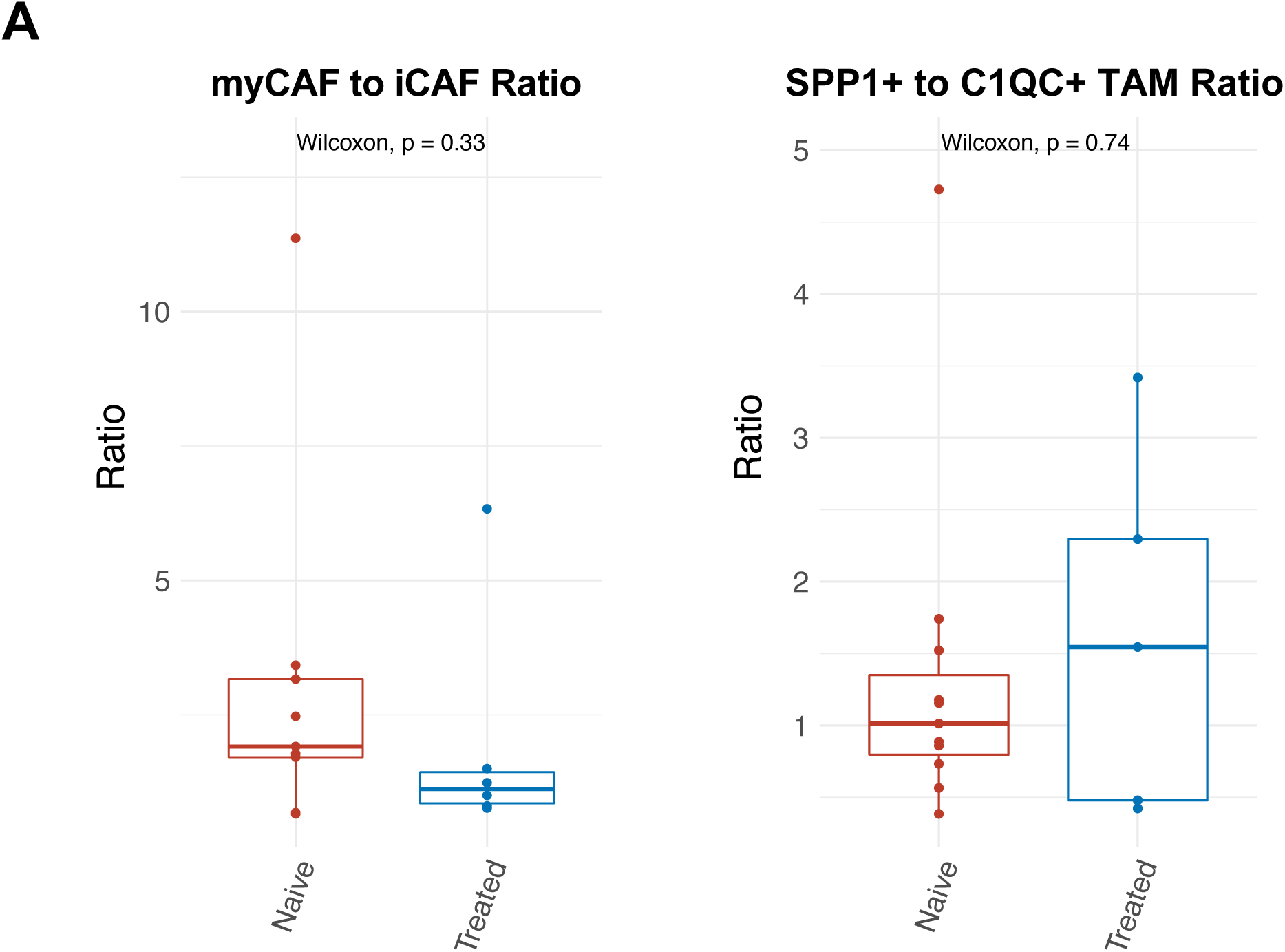
A. Ratio of CAF and TAM subpopulations in treated and untreated groups.

## References

1. Siegel, R. L., Miller, K. D., Fuchs, H. E. & Jemal, A. Cancer statistics, 2022. CA Cancer J. Clin. 72, 7–33 (2022).

2. Rahib, L., Wehner, M. R., Matrisian, L. M. & Nead, K. T. Estimated Projection of US Cancer Incidence and Death to 2040. JAMA Netw Open 4, e214708 (2021).

3. Vincent, A., Herman, J., Schulick, R., Hruban, R. H. & Goggins, M. Pancreatic cancer. Lancet 378, 607–620 (2011).

4. O’Reilly, E. M. et al. Durvalumab With or Without Tremelimumab for Patients With Metastatic Pancreatic Ductal Adenocarcinoma: A Phase 2 Randomized Clinical Trial. JAMA Oncol 5, 1431–1438 (2019).

5. Bear, A. S., Vonderheide, R. H. & O’Hara, M. H. Challenges and Opportunities for Pancreatic Cancer Immunotherapy. Cancer Cell 38, 788–802 (2020).

6. Collisson, E. A., Bailey, P., Chang, D. K. & Biankin, A. V. Molecular subtypes of pancreatic cancer. Nat. Rev. Gastroenterol. Hepatol. 16, 207–220 (2019).

7. Moffitt, R. A. et al. Virtual microdissection identifies distinct tumor- and stroma-specific subtypes of pancreatic ductal adenocarcinoma. Nat. Genet. 47, 1168–1178 (2015).

8. Wood, L. D., Canto, M. I., Jaffee, E. M. & Simeone, D. M. Pancreatic Cancer: Pathogenesis, Screening, Diagnosis and Treatment. Gastroenterology (2022) doi:10.1053/j.gastro.2022.03.056.

9. Cancer Genome Atlas Research Network. Integrated Genomic Characterization of Pancreatic Ductal Adenocarcinoma. Cancer Cell 32, 185–203.e13 (2017).

10. Ho, W. J., Jaffee, E. M. & Zheng, L. The tumour microenvironment in pancreatic cancer - clinical challenges and opportunities. Nat. Rev. Clin. Oncol. 17, 527–540 (2020).

11. Beatty, G. L., Werba, G., Lyssiotis, C. A. & Simeone, D. M. The biological underpinnings of therapeutic resistance in pancreatic cancer. Genes & Development vol. 35 940–962 (2021).

12. Moncada, R. et al. Integrating microarray-based spatial transcriptomics and single-cell RNA-seq reveals tissue architecture in pancreatic ductal adenocarcinomas. Nat. Biotechnol. 38, 333–342 (2020).

13. Azizi, E. et al. Single-Cell Map of Diverse Immune Phenotypes in the Breast Tumor Microenvironment. Cell 174, 1293–1308.e36 (2018).

14. Zhang, Y. et al. Single-cell analyses reveal key immune cell subsets associated with response to PD-L1 blockade in triple-negative breast cancer. Cancer Cell 39, 1578–1593.e8 (2021).

15. Chan, J. M. et al. Signatures of plasticity, metastasis, and immunosuppression in an atlas of human small cell lung cancer. Cancer Cell vol. 39 1479–1496.e18 (2021).

16. Wu, H. et al. Single-Cell Transcriptomics of a Human Kidney Allograft Biopsy Specimen Defines a Diverse Inflammatory Response. J. Am. Soc. Nephrol. 29, 2069–2080 (2018).

17. Kim, N. et al. Single-cell RNA sequencing demonstrates the molecular and cellular reprogramming of metastatic lung adenocarcinoma. Nat. Commun. 11, 2285 (2020).

18. Peng, J. et al. Single-cell RNA-seq highlights intra-tumoral heterogeneity and malignant progression in pancreatic ductal adenocarcinoma. Cell Research vol. 29 725–738 (2019).

19. Elyada, E. et al. Cross-Species Single-Cell Analysis of Pancreatic Ductal Adenocarcinoma Reveals Antigen-Presenting Cancer-Associated Fibroblasts. Cancer Discov. 9, 1102–1123 (2019).

20. Wang, Y. et al. Single-cell analysis of pancreatic ductal adenocarcinoma identifies a novel fibroblast subtype associated with poor prognosis but better immunotherapy response. Cell Discov 7, 36 (2021).

21. Steele, N. G. et al. Multimodal Mapping of the Tumor and Peripheral Blood Immune Landscape in Human Pancreatic Cancer. Nat Cancer 1, 1097–1112 (2020).

22. Lee, J. J. et al. Elucidation of Tumor-Stromal Heterogeneity and the Ligand-Receptor Interactome by Single-Cell Transcriptomics in Real-world Pancreatic Cancer Biopsies. Clin. Cancer Res. 27, 5912–5921 (2021).

23. Raghavan, S. et al. Microenvironment drives cell state, plasticity, and drug response in pancreatic cancer. Cell 184, 6119–6137.e26 (2021).

24. Öhlund, D. et al. Distinct populations of inflammatory fibroblasts and myofibroblasts in pancreatic cancer. J. Exp. Med. 214, 579–596 (2017).

25. Puram, S. V. et al. Single-Cell Transcriptomic Analysis of Primary and Metastatic Tumor Ecosystems in Head and Neck Cancer. Cell 171, 1611–1624.e24 (2017).

26. Yang, N., Mosher, R., Seo, S., Beebe, D. & Friedl, A. Syndecan-1 in breast cancer stroma fibroblasts regulates extracellular matrix fiber organization and carcinoma cell motility. Am. J. Pathol. 178, 325–335 (2011).

27. Cheng, H.-W. et al. CCL19-producing fibroblastic stromal cells restrain lung carcinoma growth by promoting local antitumor T-cell responses. J. Allergy Clin. Immunol. 142, 1257–1271.e4 (2018).

28. Crinier, A. et al. High-Dimensional Single-Cell Analysis Identifies Organ-Specific Signatures and Conserved NK Cell Subsets in Humans and Mice. Immunity 49, 971–986.e5 (2018).

29. de Andrade, L. F. et al. Discovery of specialized NK cell populations infiltrating human melanoma metastases. JCI Insight 4, (2019).

30. Li, H. et al. Dysfunctional CD8 T cells form a proliferative, dynamically regulated compartment within human melanoma. Cell 176, 775–789.e18 (2019).

31. Kwak, T. et al. Distinct Populations of Immune-Suppressive Macrophages Differentiate from Monocytic Myeloid-Derived Suppressor Cells in Cancer. Cell Rep. 33, 108571 (2020).

32. House, I. G. et al. Macrophage-Derived CXCL9 and CXCL10 Are Required for Antitumor Immune Responses Following Immune Checkpoint Blockade. Clinical Cancer Research vol. 26 487–504 (2020).

33. Zhang, L. et al. Single-Cell Analyses Inform Mechanisms of Myeloid-Targeted Therapies in Colon Cancer. Cell 181, 442–459.e29 (2020).

34. Efremova, M., Vento-Tormo, M., Teichmann, S. A. & Vento-Tormo, R. CellPhoneDB: inferring cell-cell communication from combined expression of multi-subunit ligand-receptor complexes. Nat. Protoc. 15, 1484–1506 (2020).

35. Gunaydin, G. CAFs Interacting With TAMs in Tumor Microenvironment to Enhance Tumorigenesis and Immune Evasion. Front. Oncol. 11, 668349 (2021).

36. Arwert, E. N. et al. A Unidirectional Transition from Migratory to Perivascular Macrophage Is Required for Tumor Cell Intravasation. Cell Rep. 23, 1239–1248 (2018).

37. Hu, Z. I. et al. Evaluating Mismatch Repair Deficiency in Pancreatic Adenocarcinoma: Challenges and Recommendations. Clin. Cancer Res. 24, 1326–1336 (2018).

38. Beltra, J.-C. et al. Developmental Relationships of Four Exhausted CD8 T Cell Subsets Reveals Underlying Transcriptional and Epigenetic Landscape Control Mechanisms. Immunity 52, 825– 841.e8 (2020).

39. Chan-Seng-Yue, M. et al. Transcription phenotypes of pancreatic cancer are driven by genomic events during tumor evolution. Nat. Genet. 52, 231–240 (2020).

40. Grünwald, B. T. et al. Spatially confined sub-tumor microenvironments in pancreatic cancer. Cell 184, 5577–5592.e18 (2021).

41. Espiau-Romera, P., Courtois, S., Parejo-Alonso, B. & Sancho, P. Molecular and Metabolic Subtypes Correspondence for Pancreatic Ductal Adenocarcinoma Classification. J. Clin. Med. Res. 9, (2020).

42. O’Kane, G. M. et al. GATA6 Expression Distinguishes Classical and Basal-like Subtypes in Advanced Pancreatic Cancer. Clin. Cancer Res. 26, 4901–4910 (2020).

43. Mills, C. D., Kincaid, K., Alt, J. M., Heilman, M. J. & Hill, A. M. M-1/M-2 macrophages and the Th1/Th2 paradigm. J. Immunol. 164, 6166–6173 (2000).

44. Anderson, C. F. & Mosser, D. M. A novel phenotype for an activated macrophage: the type 2 activated macrophage. J. Leukoc. Biol. 72, 101–106 (2002).

45. Italiani, P. & Boraschi, D. From Monocytes to M1/M2 Macrophages: Phenotypical vs. Functional Differentiation. Front. Immunol. 5, 514 (2014).

46. Cheng, S. et al. A pan-cancer single-cell transcriptional atlas of tumor infiltrating myeloid cells. Cell 184, 792–809.e23 (2021).

47. Liu, Y. et al. Immune phenotypic linkage between colorectal cancer and liver metastasis. Cancer Cell 40, 424–437.e5 (2022).

48. Aung, K. L. et al. Genomics-Driven Precision Medicine for Advanced Pancreatic Cancer: Early Results from the COMPASS Trial. Clin. Cancer Res. 24, 1344–1354 (2018).

49. Porter, R. L. et al. Epithelial to mesenchymal plasticity and differential response to therapies in pancreatic ductal adenocarcinoma. Proc. Natl. Acad. Sci. U. S. A. (2019) doi:10.1073/pnas.1914915116.

50. Aiello, N. M. et al. EMT Subtype Influences Epithelial Plasticity and Mode of Cell Migration. Dev. Cell 45, 681–695.e4 (2018).

51. Arumugam, T. et al. Epithelial to mesenchymal transition contributes to drug resistance in pancreatic cancer. Cancer Res. 69, 5820–5828 (2009).

52. Rhim, A. D. et al. EMT and dissemination precede pancreatic tumor formation. Cell 148, 349– 361 (2012).

53. Bulle, A. & Lim, K.-H. Beyond just a tight fortress: contribution of stroma to epithelial-mesenchymal transition in pancreatic cancer. Signal Transduct Target Ther 5, 249 (2020).

54. Bielenberg, D. R. & Zetter, B. R. The Contribution of Angiogenesis to the Process of Metastasis. Cancer J. 21, 267–273 (2015).

55. Hanahan, D. & Weinberg, R. A. Hallmarks of Cancer: The Next Generation. Cell 144, 646–674 (2011).

56. Takagi, K., Takada, T. & Amano, H. A high peripheral microvessel density count correlates with a poor prognosis in pancreatic cancer. J. Gastroenterol. 40, 402–408 (2005).

57. Annese, T., Tamma, R., Ruggieri, S. & Ribatti, D. Angiogenesis in Pancreatic Cancer: Pre- Clinical and Clinical Studies. Cancers 11, (2019).

58. Feig, C. et al. Targeting CXCL12 from FAP-expressing carcinoma-associated fibroblasts synergizes with anti-PD-L1 immunotherapy in pancreatic cancer. Proc. Natl. Acad. Sci. U. S. A. 110, 20212–20217 (2013).

59. Vonderheide, R. H. The Immune Revolution: A Case for Priming, Not Checkpoint. Cancer Cell 33, 563–569 (2018).

60. Freed-Pastor, W. A. et al. The CD155/TIGIT axis promotes and maintains immune evasion in neoantigen-expressing pancreatic cancer. Cancer Cell 39, 1342–1360.e14 (2021).

61. Mullard, A. Roche’s anti-TIGIT drug suffers a phase III cancer setback. Nat. Rev. Drug Discov. 21, 327 (2022).

62. Zheng, G. X. Y. et al. Massively parallel digital transcriptional profiling of single cells. Nat. Commun. 8, 14049 (2017).

63. Butler, A., Hoffman, P., Smibert, P., Papalexi, E. & Satija, R. Integrating single-cell transcriptomic data across different conditions, technologies, and species. Nat. Biotechnol. 36, 411–420 (2018).

64. Dolgalev, I. & Yeaton, A. scooter: Streamlined scRNA-Seq Analysis Pipeline. (2021).

65. Young, M. D. & Behjati, S. SoupX removes ambient RNA contamination from droplet-based single-cell RNA sequencing data. Gigascience 9, (2020).

66. Germain, P.-L., Lun, A., Macnair, W. & Robinson, M. D. Doublet identification in single-cell sequencing data using scDblFinder. F1000Research vol. 10 979 (2021).

67. Stuart, T. et al. Comprehensive Integration of Single-Cell Data. Cell vol. 177 1888–1902.e21 (2019).

68. Becht, E. et al. Dimensionality reduction for visualizing single-cell data using UMAP. Nat. Biotechnol. (2018) doi:10.1038/nbt.4314.

69. Waltman, L. & van Eck, N. J. A smart local moving algorithm for large-scale modularity-based community detection. Eur. Phys. J. B 86, (2013).

70. Tickle, T., Tirosh, I., Georgescu, C., Brown, M. & Haas, B. inferCNV of the Trinity CTAT Project. (Bioconductor, 2019). doi:10.18129/B9.BIOC.INFERCNV.

71. Subramanian, A. et al. Gene set enrichment analysis: a knowledge-based approach for interpreting genome-wide expression profiles. Proc. Natl. Acad. Sci. U. S. A. 102, 15545–15550 (2005).

72. Colaprico, A. et al. TCGAbiolinks: an R/Bioconductor package for integrative analysis of TCGA data. Nucleic Acids Research vol. 44 e71–e71 (2016).

73. Kassambara, A., Kosinski, M. & Biecek, P. survminer. (2021).

74. Therneau, T. M. A Package for Survival Analysis in R. https://CRAN.R-project.org/package=survival (2022).

