## Extended Data for "Single-Cell RNA Sequencing Reveals the Effects of Chemotherapy on Human Pancreatic Adenocarcinoma and its Tumor Microenvironment"

| | |
|---|---|
| **Patient Number** | P01 |
| **Age** | 78 |
| **Gender** | Male |
| **Stage at Diagnosis** | IV |
| **Treatment before tissue collection** | No |
| **Tissue site** | Liver |
| **Procedure** | Biopsy |
| **Pathology** | Moderately differentiated adenocarcinoma |
| **Mutations** | KRAS G12R, TP53 Y327* |

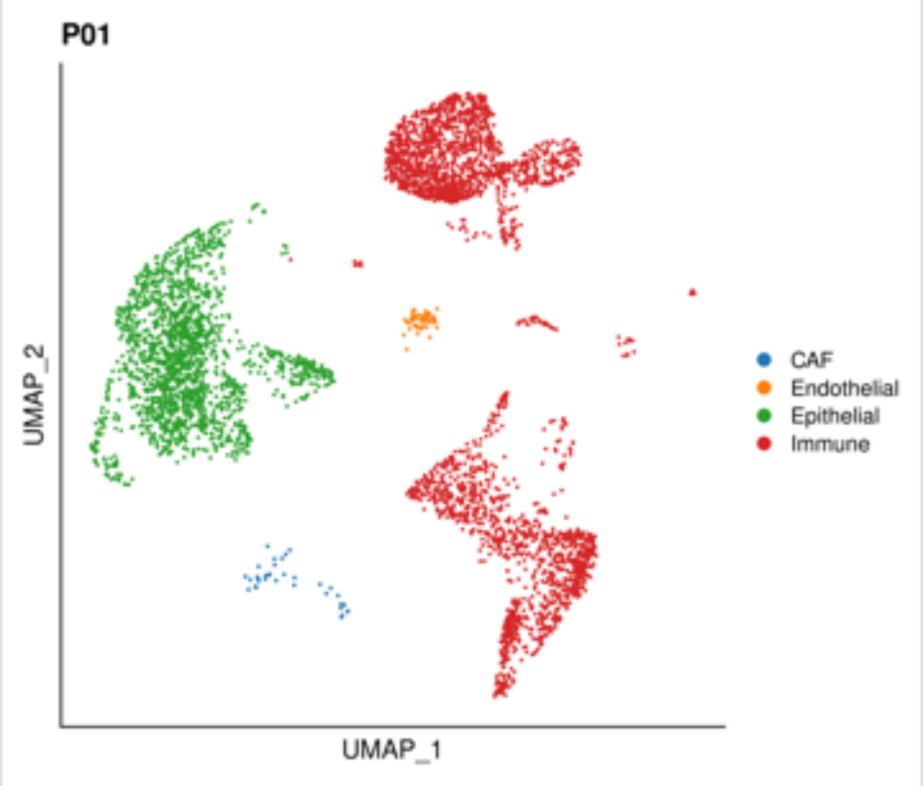


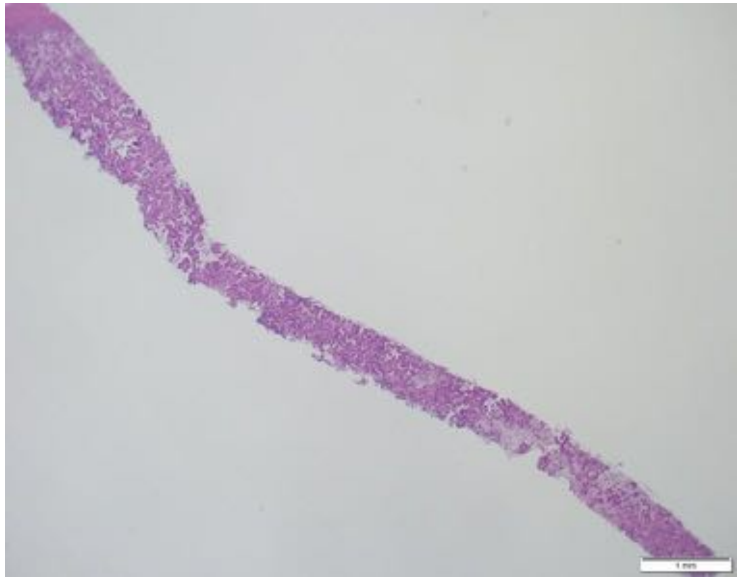

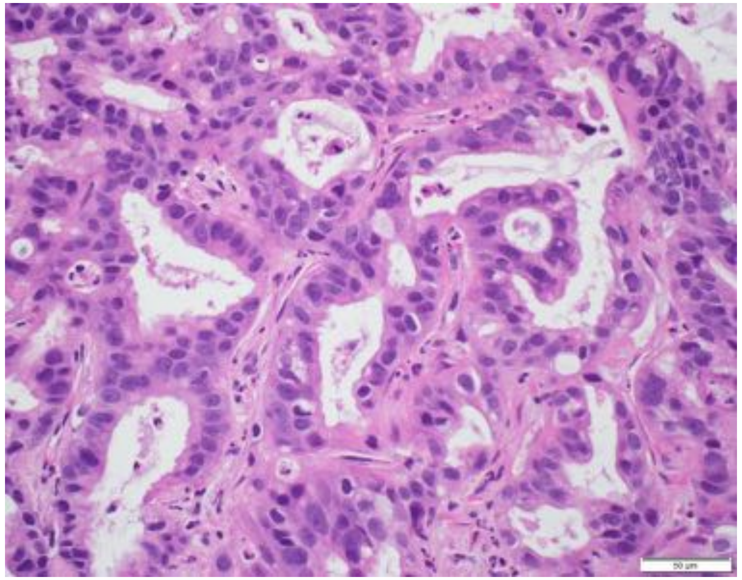

| | |
|---|---|
| **Patient Number** | P02 |
| **Age** | 74 |
| **Gender** | Male |
| **Stage at Diagnosis** | IV |
| **Treatment before tissue collection** | No |
| **Tissue site** | Liver |
| **Procedure** | Biopsy |
| **Pathology** | Moderately differentiated adenocarcinoma |
| **Mutations** | KRAS G12D, TP53 c.673-1G>A |

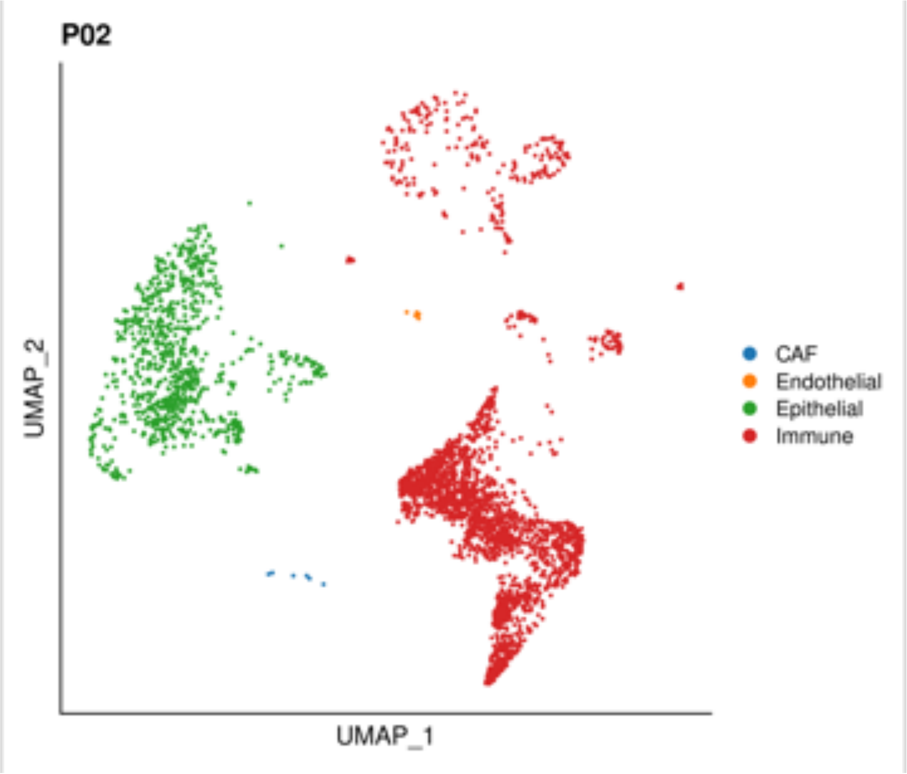


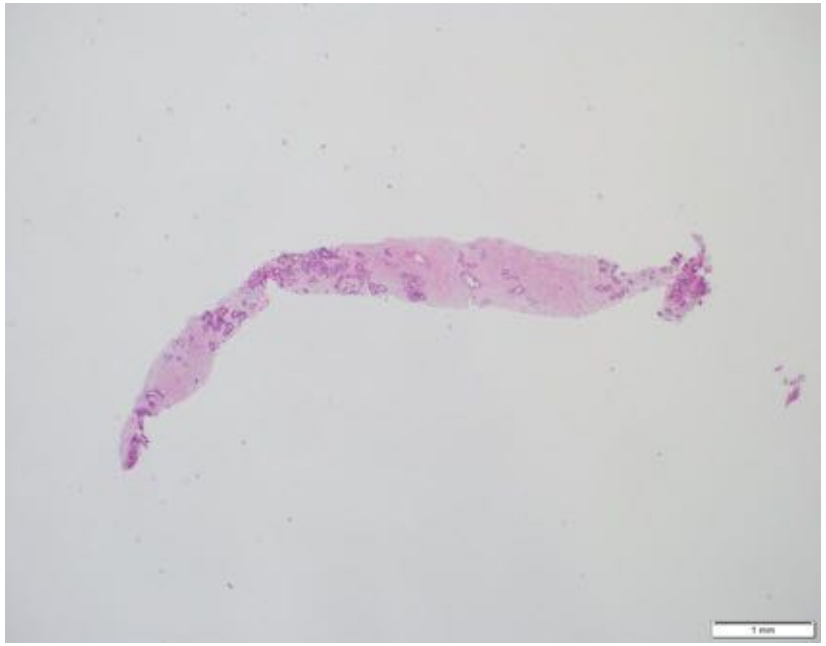

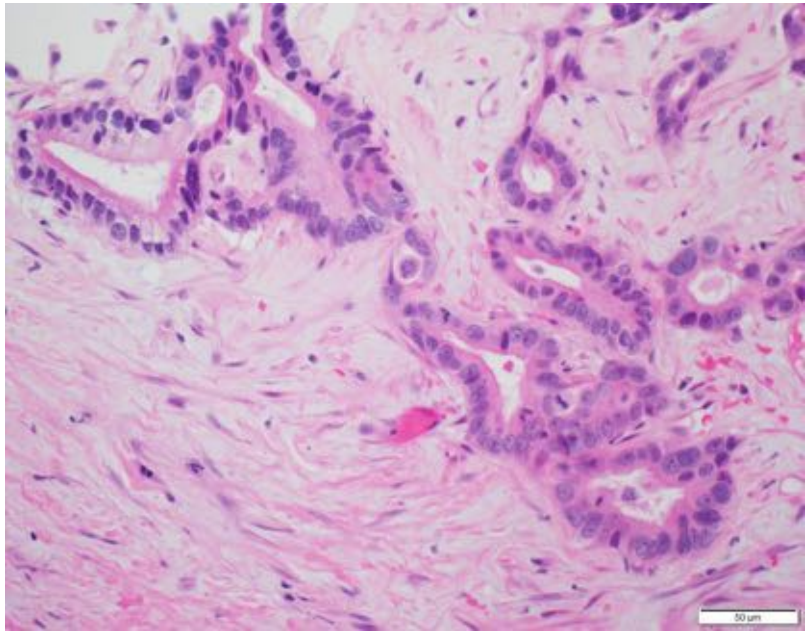

| | |
|---|---|
| **Patient Number** | P03 |
| **Age** | 81 |
| **Gender** | Male |
| **Stage at Diagnosis** | IV |
| **Treatment before tissue collection** | Yes |
| **Therapeutics** | FFX-based |
| **Tissue site** | Pancreas |
| **Procedure** | Resection |
| **Pathology** | Poorly differentiated adenocarcinoma |
| **Mutations** | WT |

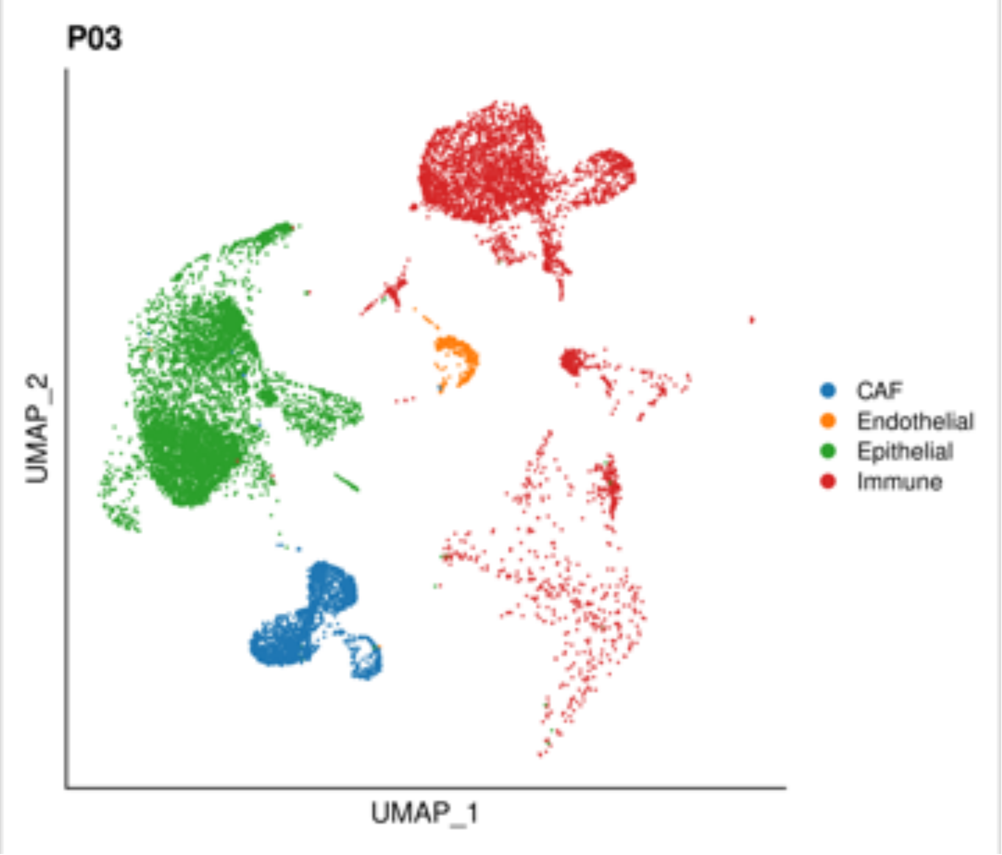


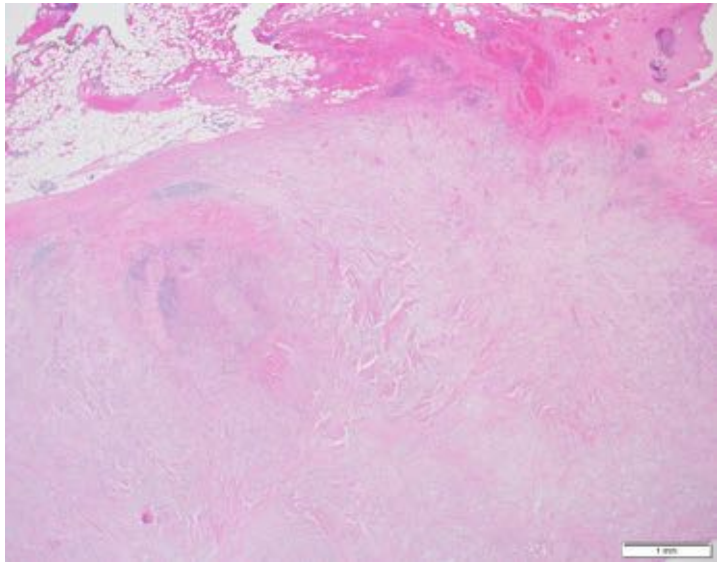

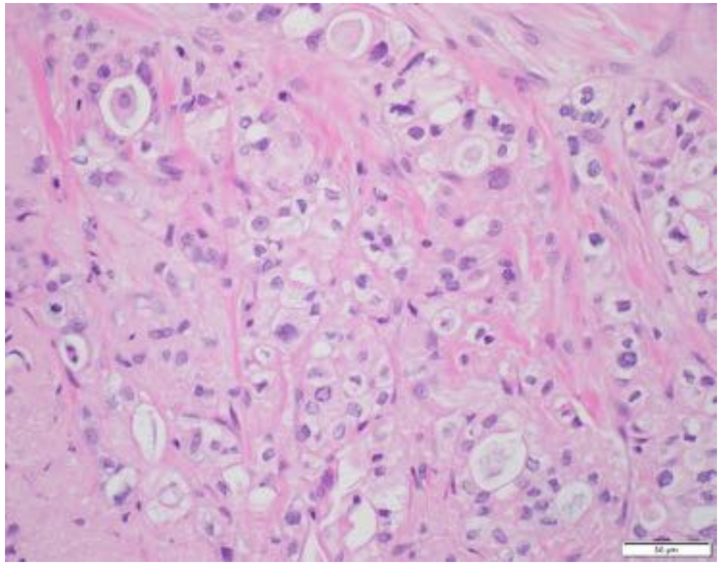

| | |
|---|---|
| **Patient Number** | P04 |
| **Age** | 85 |
| **Gender** | Female |
| **Stage at Diagnosis** | IB |
| **Treatment before tissue collection** | No |
| **Tissue site** | Pancreas |
| **Procedure** | Resection |
| **Pathology** | Moderately to poorly differentiated adenocarcinoma |
| **Mutations** | KRAS G12D, TP53 V147G |

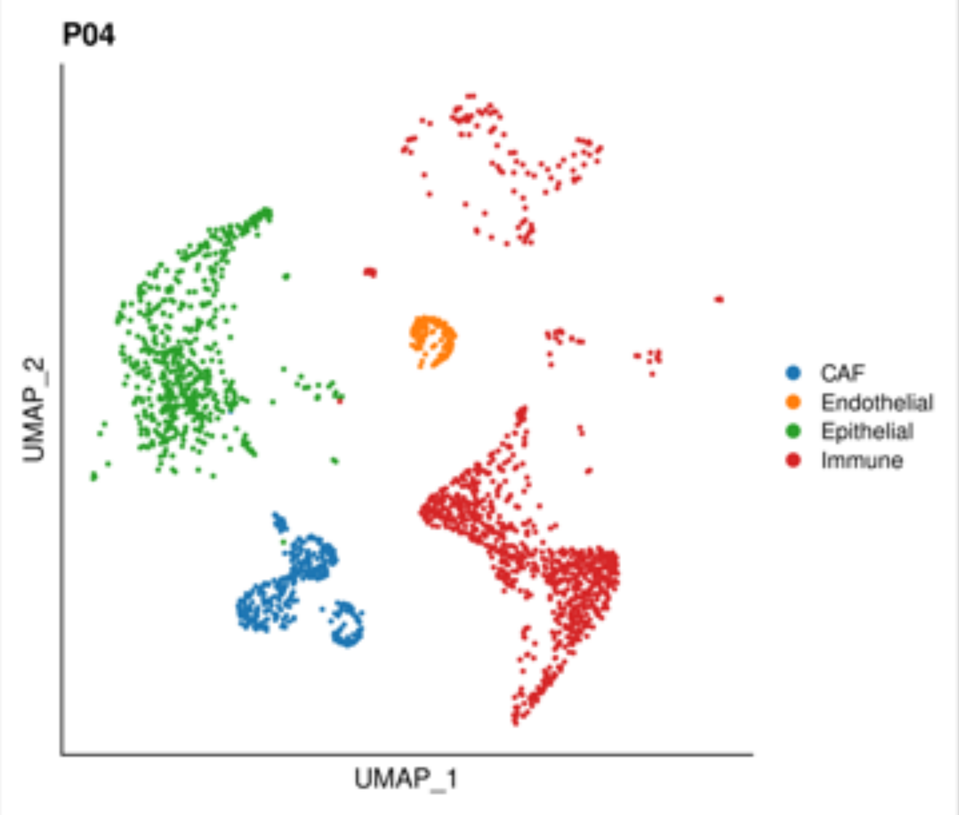


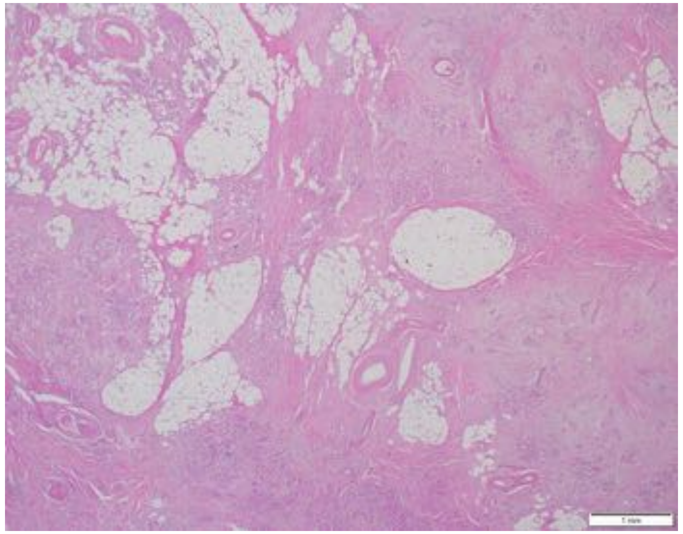

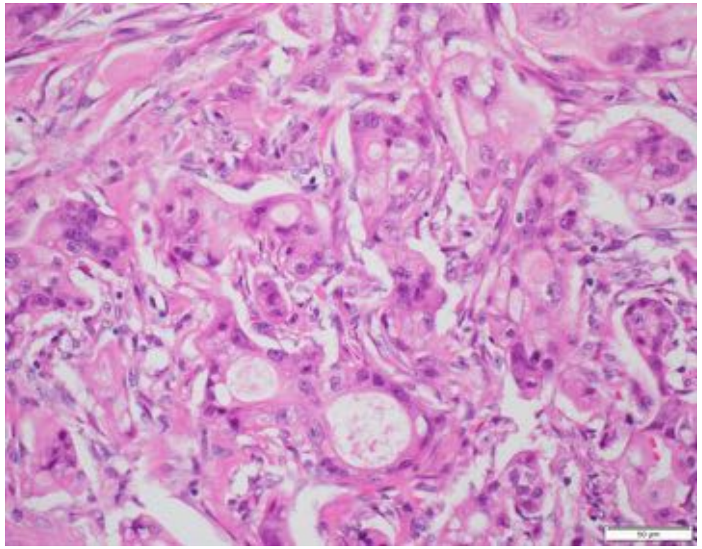

| | |
|---|---|
| **Patient Number** | P05 |
| **Age** | 69 |
| **Gender** | Female |
| **Stage at Diagnosis** | III |
| **Treatment before tissue collection** | No |
| **Tissue site** | Pancreas |
| **Procedure** | Resection |
| **Pathology** | Moderately differentiated adenocarcinoma |
| **Mutations** | KRAS G12D, TP53 H193L, SMAD4 R135* |

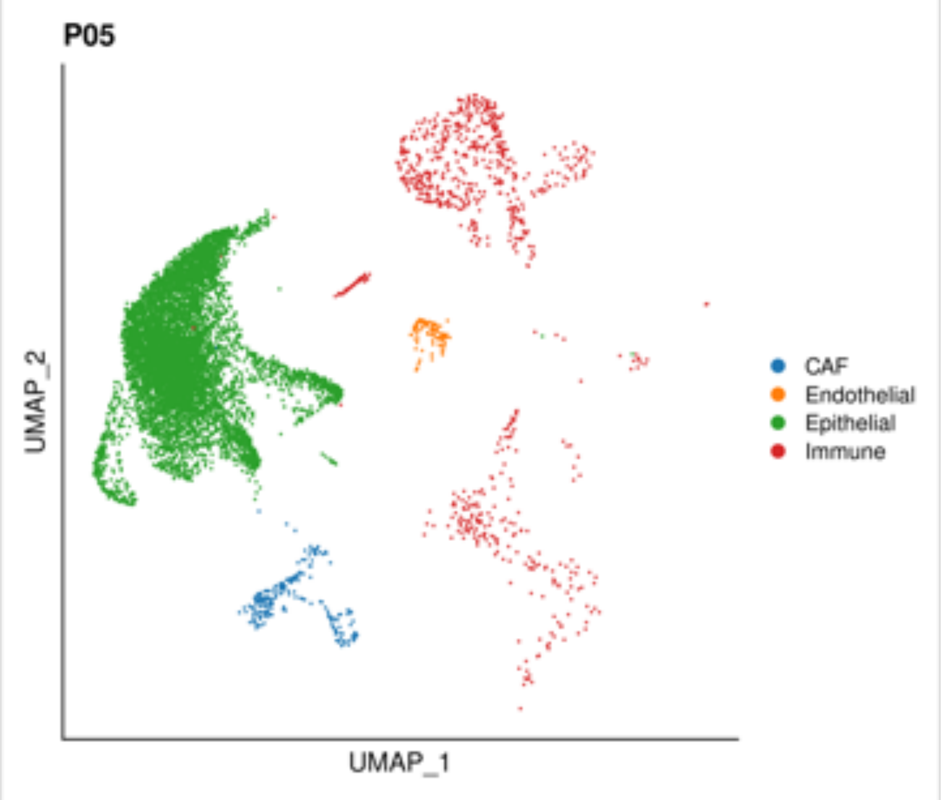


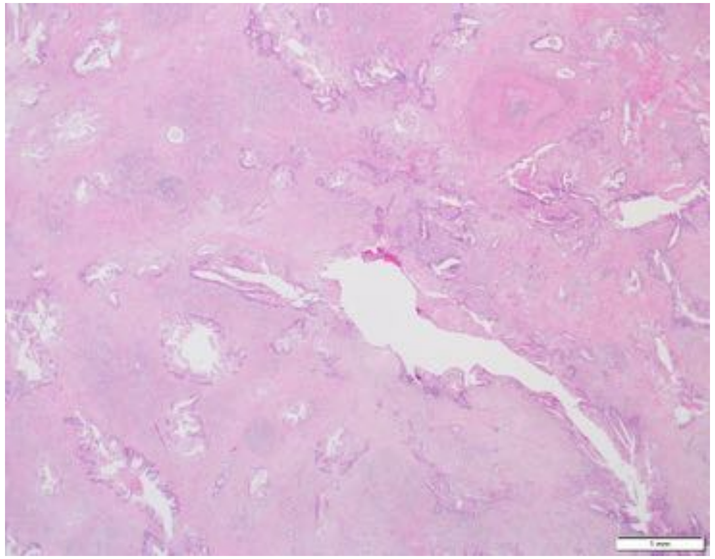

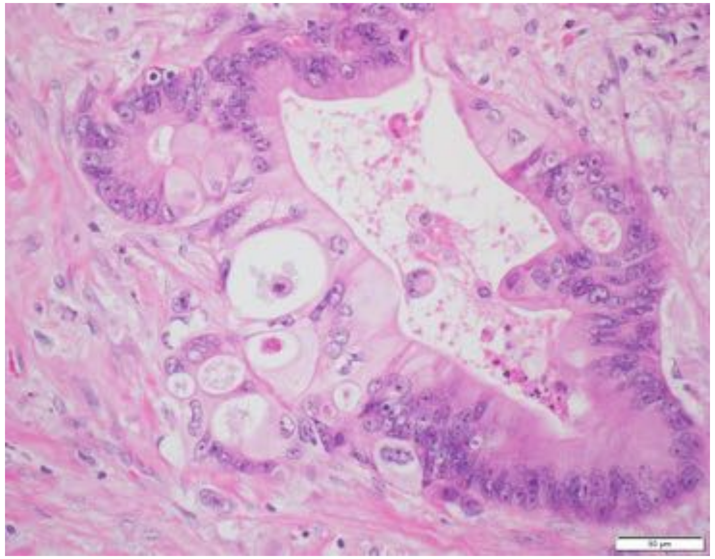

| | |
|---|---|
| **Patient Number** | P06 |
| **Age** | 55 |
| **Gender** | Male |
| **Stage at Diagnosis** | III |
| **Treatment before tissue collection** | Yes |
| **Therapeutics** | G/A |
| **Tissue site** | Pancreas |
| **Procedure** | Biopsy |
| **Pathology** | Moderately to poorly differentiated adenocarcinoma |
| **Mutations** | N/A |

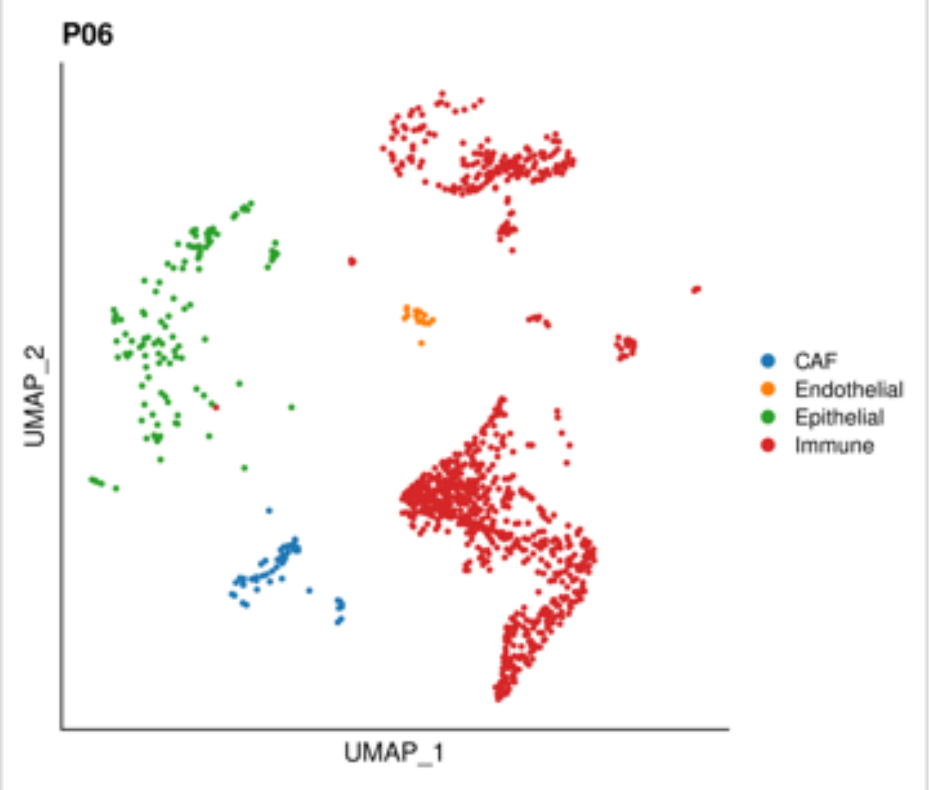


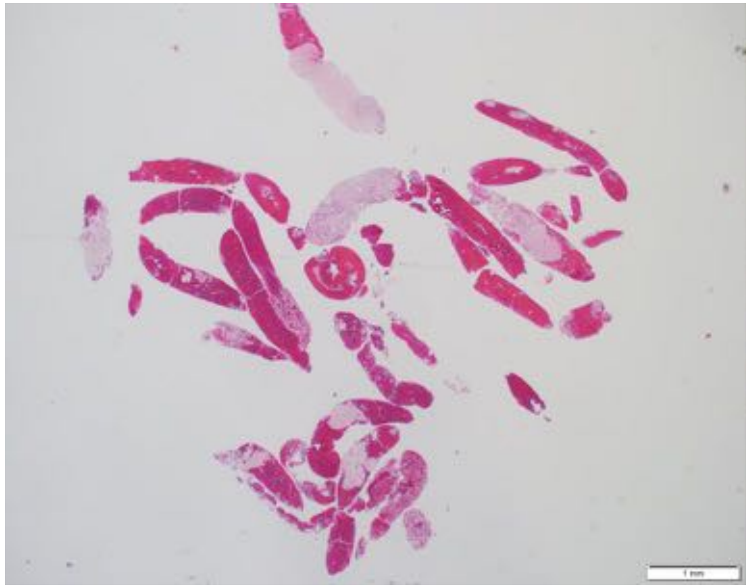

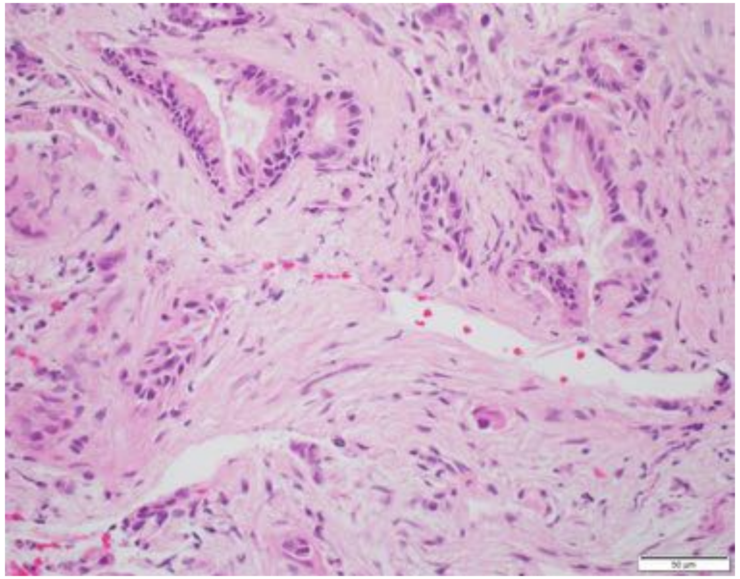

| | |
|---|---|
| **Patient Number** | P07 |
| **Age** | 70 |
| **Gender** | Female |
| **Stage at Diagnosis** | IB |
| **Treatment before tissue collection** | No |
| **Tissue site** | Pancreas |
| **Procedure** | Resection |
| **Pathology** | Well to moderately differentiated adenocarcinoma |
| **Mutations** | KRAS G12V |

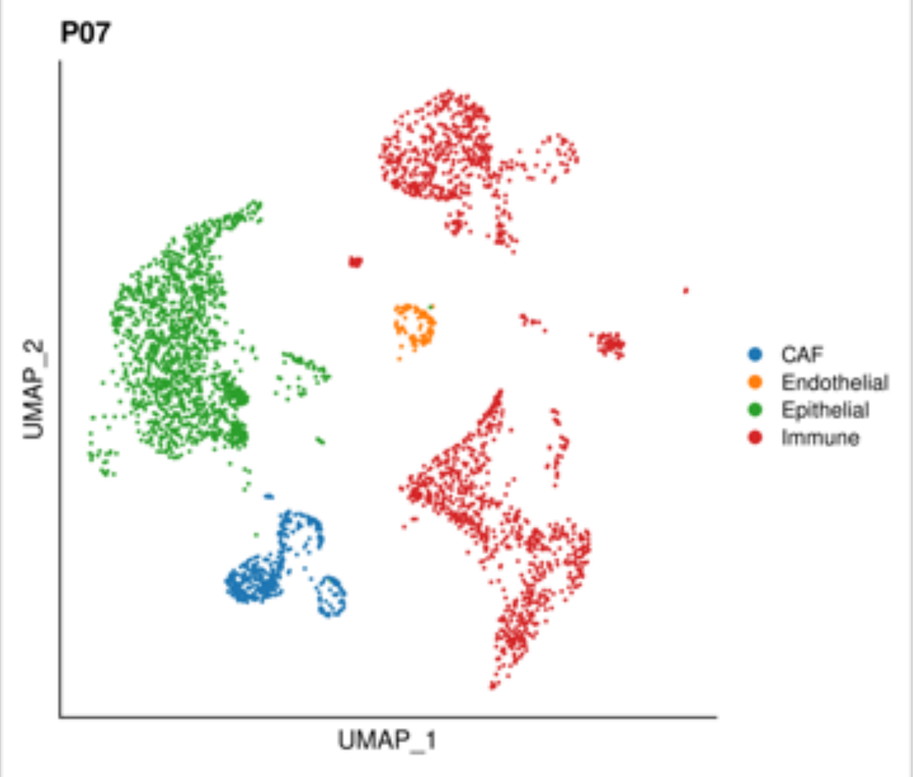


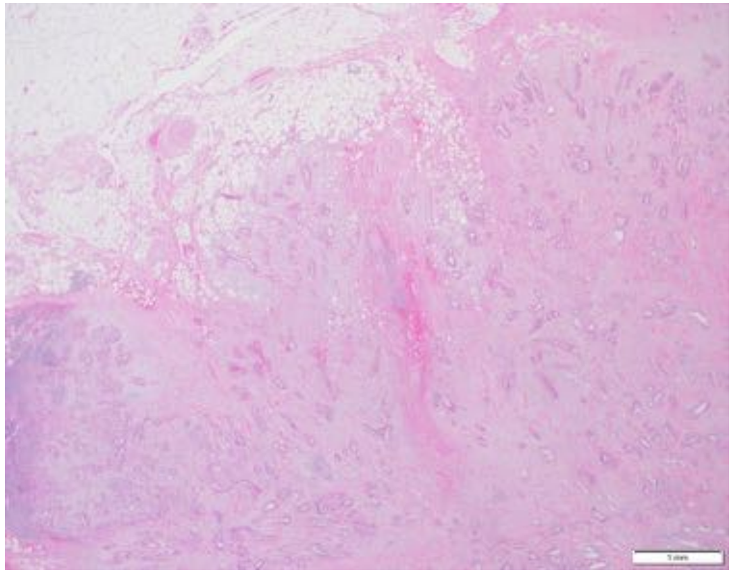

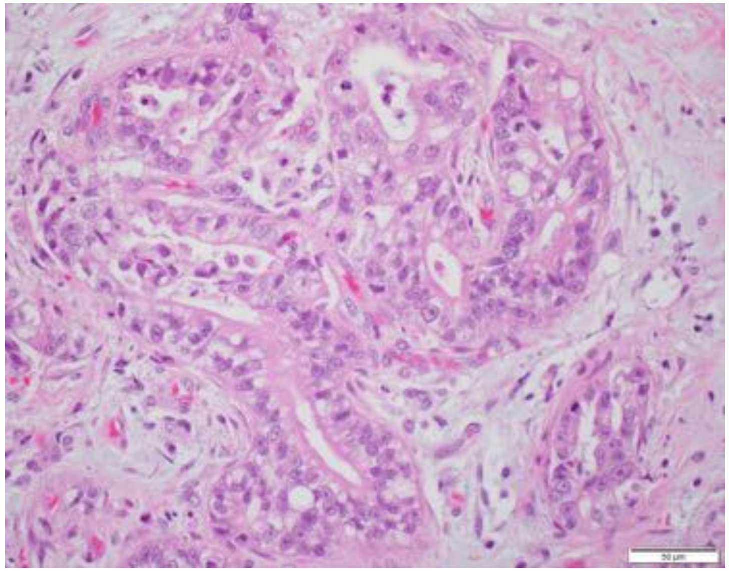

| | |
|---|---|
| **Patient Number** | P08 |
| **Age** | 87 |
| **Gender** | Female |
| **Stage at Diagnosis** | IIB |
| **Treatment before tissue collection** | Yes |
| **Therapeutics** | G/A |
| **Tissue site** | Pancreas |
| **Procedure** | Resection |
| **Pathology** | Moderately differentiated adenocarcinoma |
| **Mutations** | KRAS G12D, TP53 C141Y |

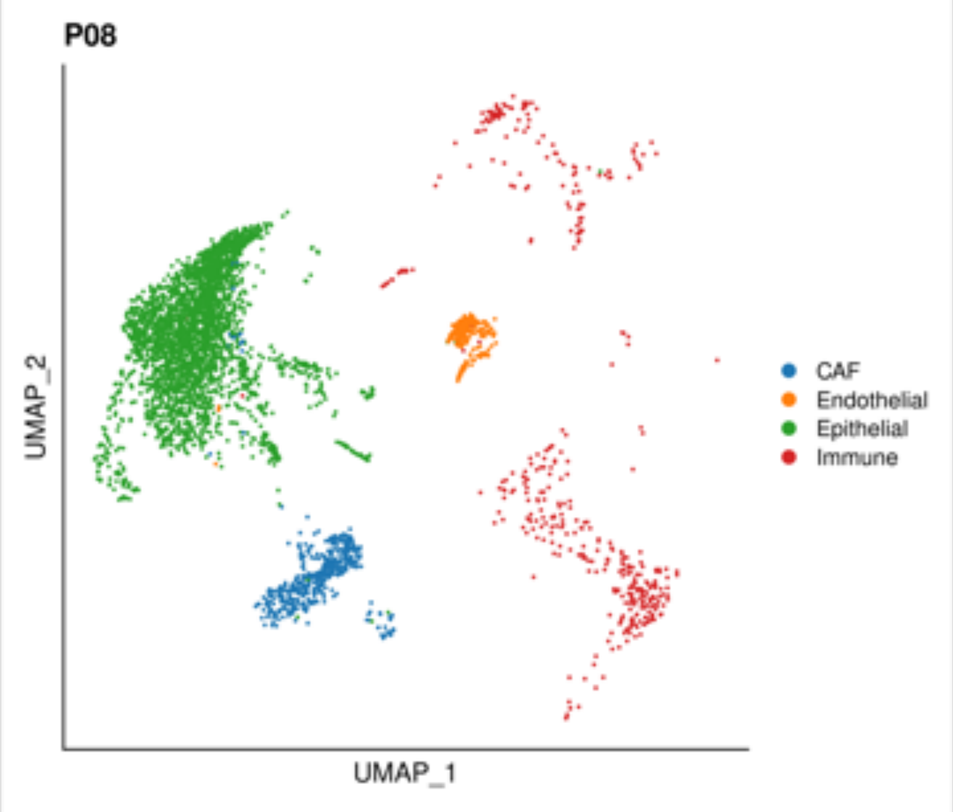


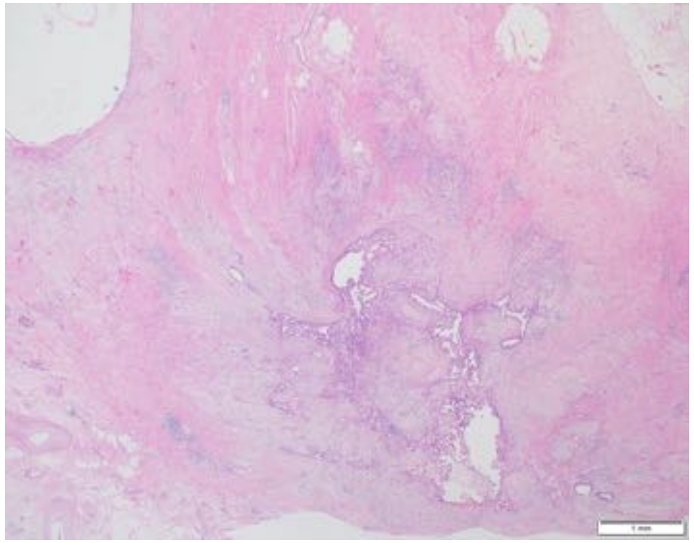

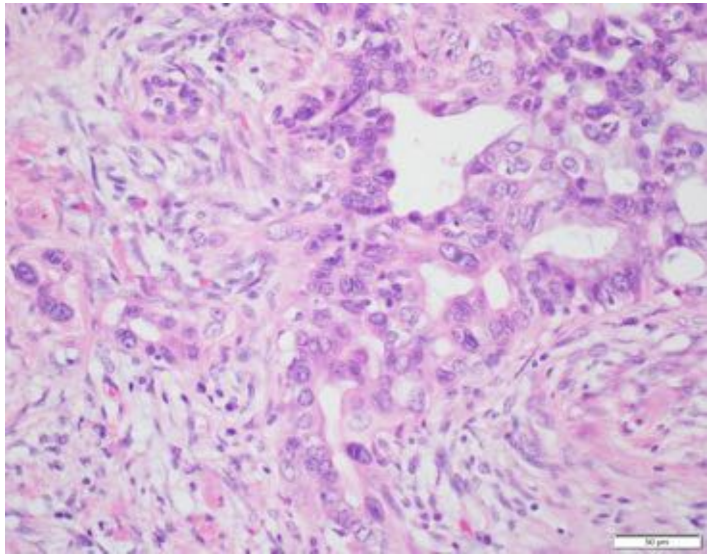

| | |
|---|---|
| **Patient Number** | P09 |
| **Age** | 63 |
| **Gender** | Male |
| **Stage at Diagnosis** | IV |
| **Treatment before tissue collection** | No |
| **Tissue site** | Pancreas |
| **Procedure** | Biopsy |
| **Pathology** | Well differentiated adenocarcinoma |
| **Mutations** | N/A |

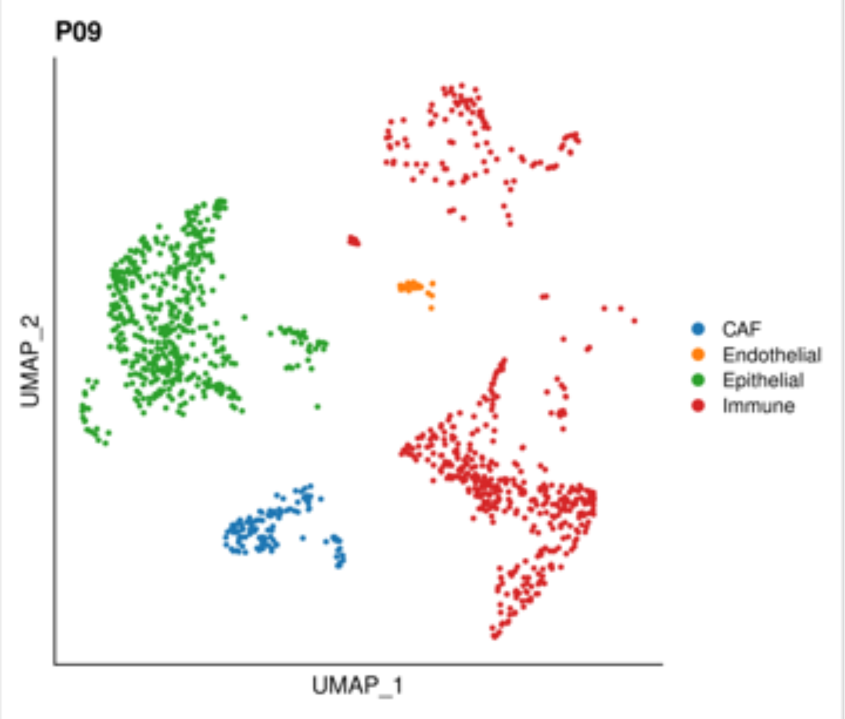


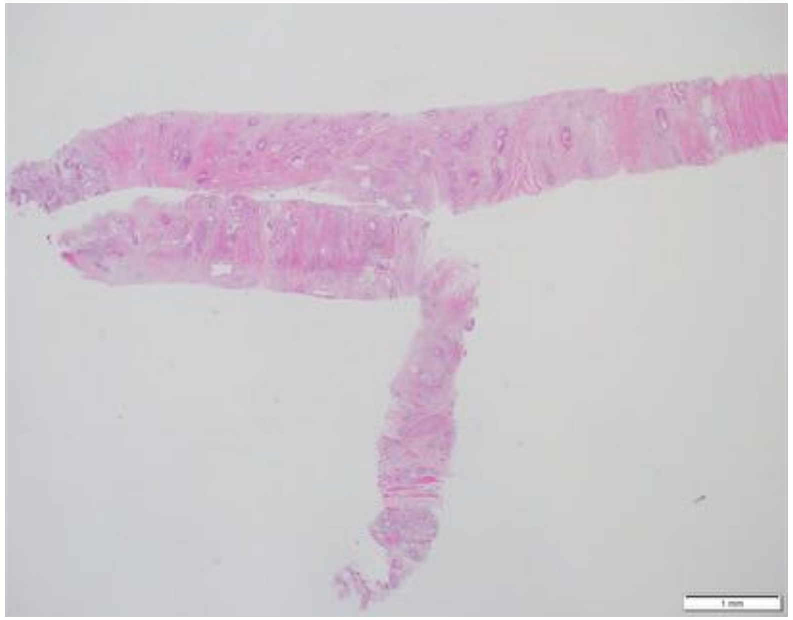

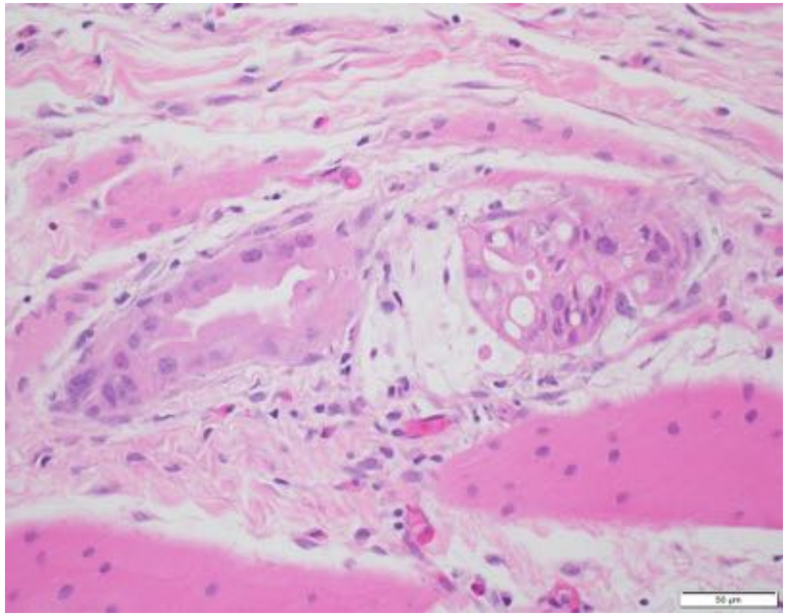

| | |
|---|---|
| **Patient Number** | P10 |
| **Age** | 48 |
| **Gender** | Female |
| **Stage at Diagnosis** | IIB |
| **Treatment before tissue collection** | Yes |
| **Therapeutics** | FFX-based |
| **Tissue site** | Pancreas |
| **Procedure** | Resection |
| **Pathology** | Well differentiated adenocarcinoma |
| **Mutations** | KRAS G12V |

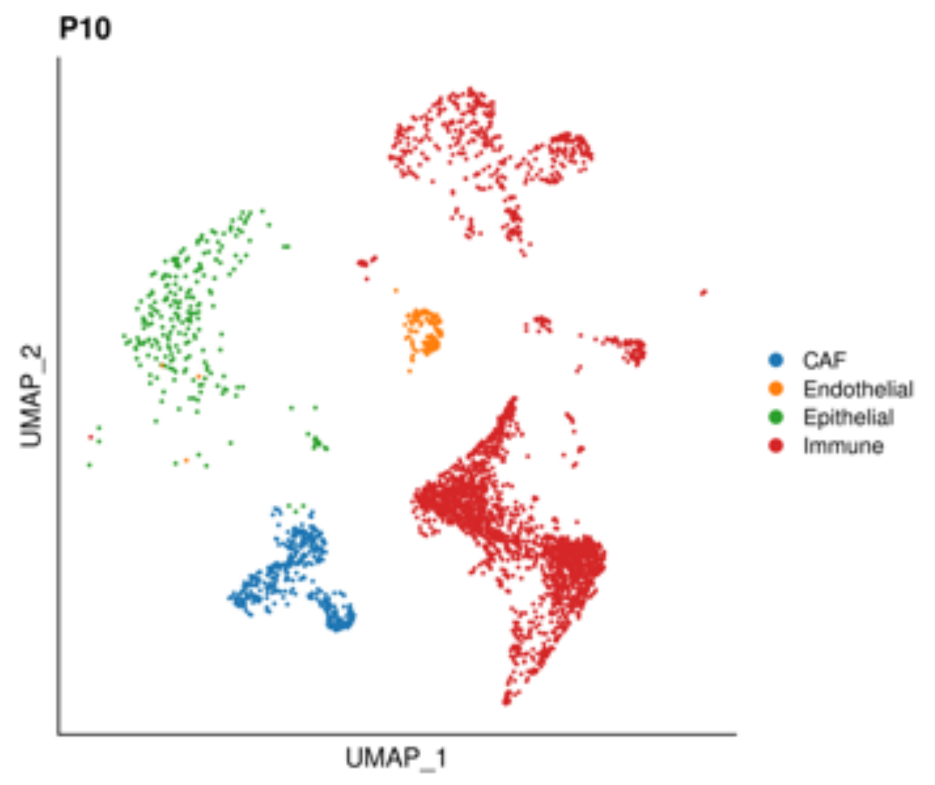


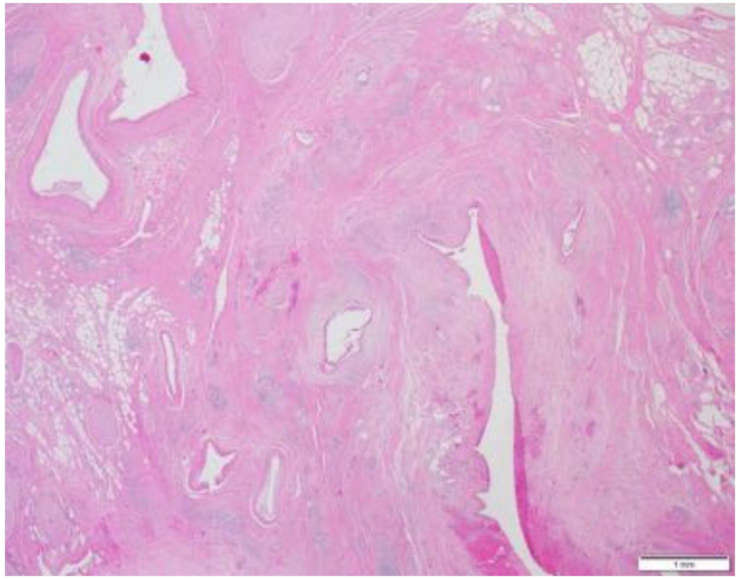

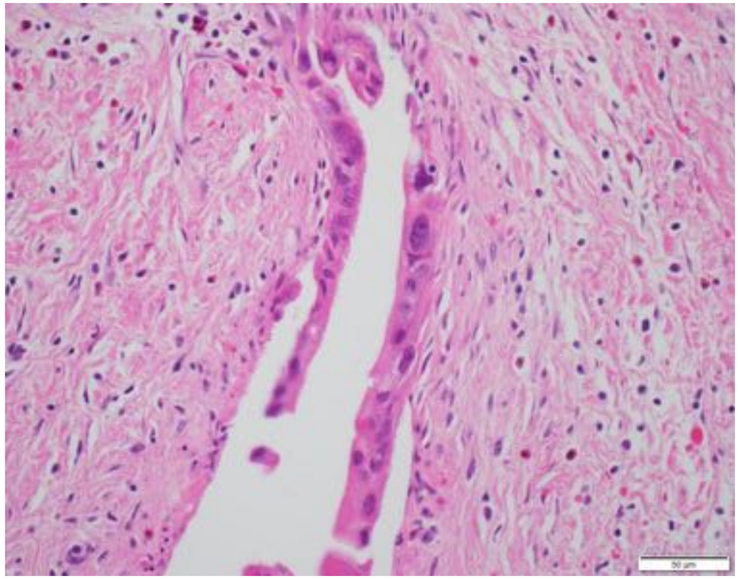

| | |
|---|---|
| **Patient Number** | P11 |
| **Age** | 73 |
| **Gender** | Female |
| **Stage at Diagnosis** | IV |
| **Treatment before tissue collection** | No |
| **Tissue site** | Liver |
| **Procedure** | Biopsy |
| **Pathology** | Moderately differentiated adenocarcinoma |
| **Mutations** | KRAS G12L, TP53 R175H, CDKN2A intron 1 rearrangement |

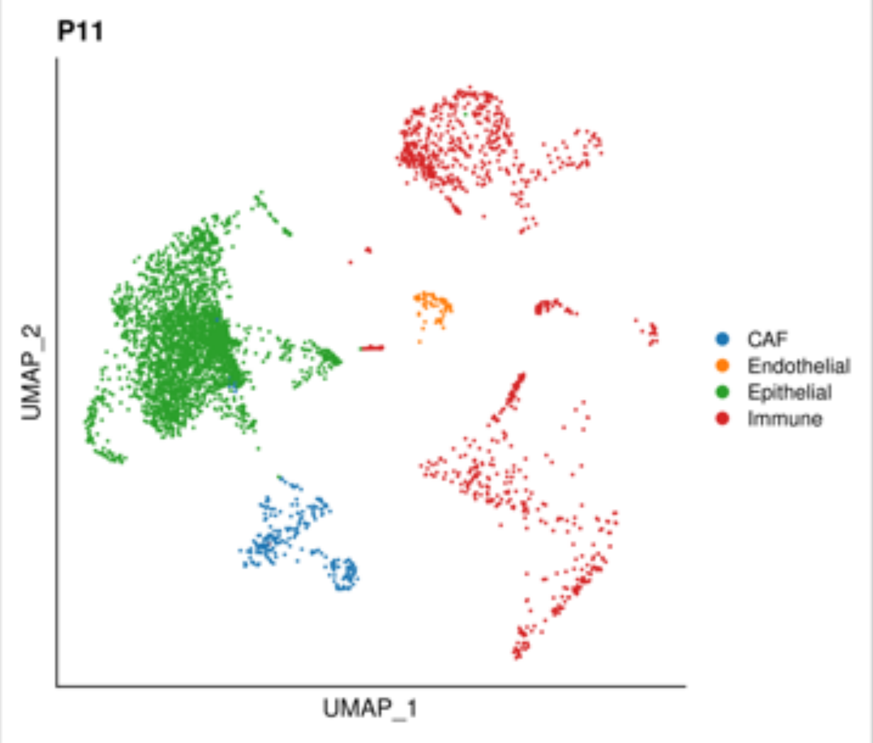


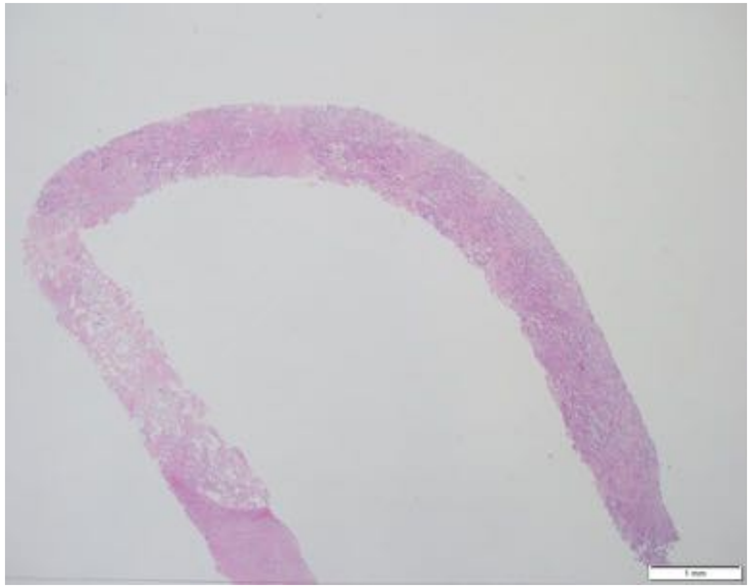

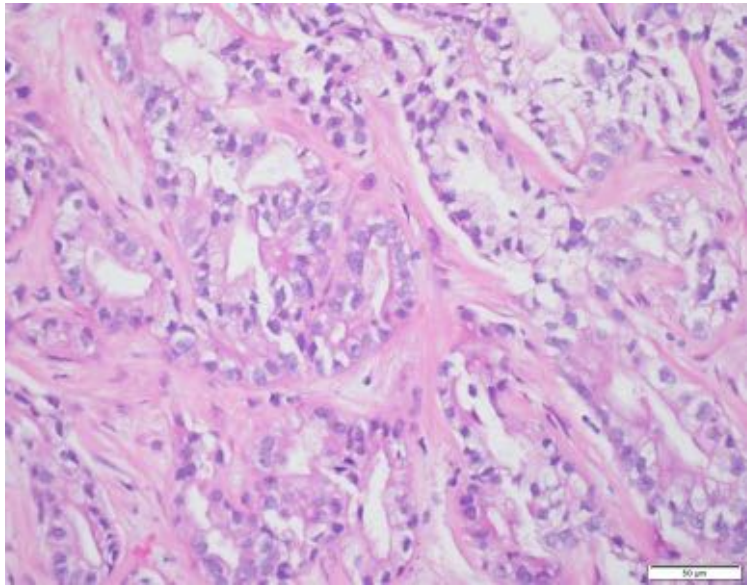

| | |
|---|---|
| **Patient Number** | P12 |
| **Age** | 68 |
| **Gender** | Male |
| **Stage at Diagnosis** | IV |
| **Treatment before tissue collection** | Yes |
| **Therapeutics** | GA |
| **Tissue site** | Pancreas |
| **Procedure** | Biopsy |
| **Pathology** | Moderately differentiated adenocarcinoma |
| **Mutations** | KRAS G12V, TP53 R306*, SMAD4 R361C, CDKN2A W110* |

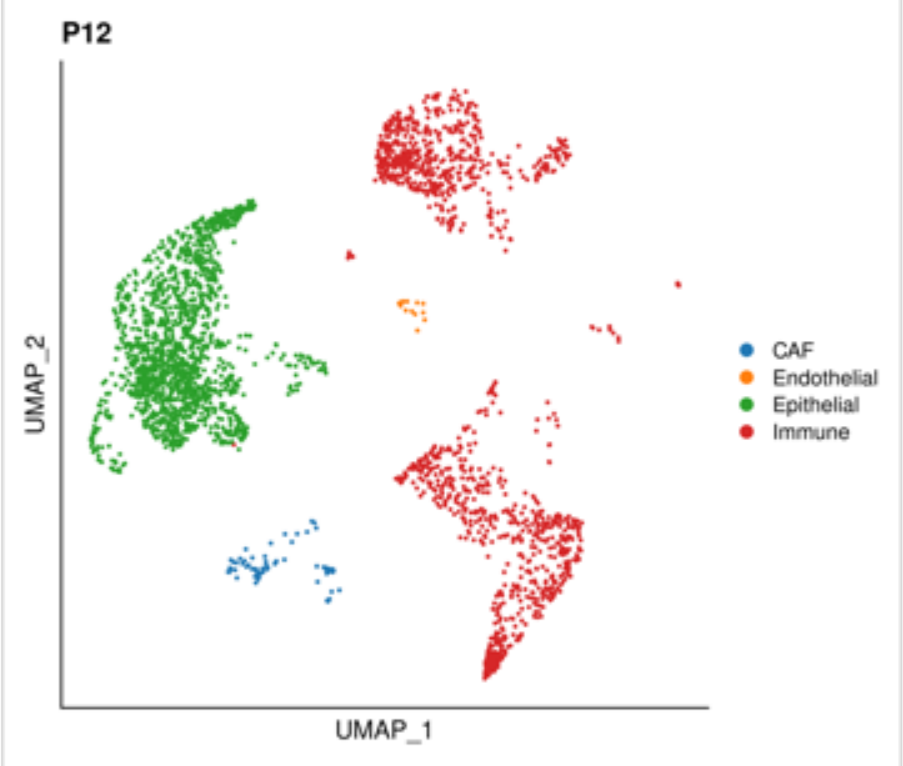


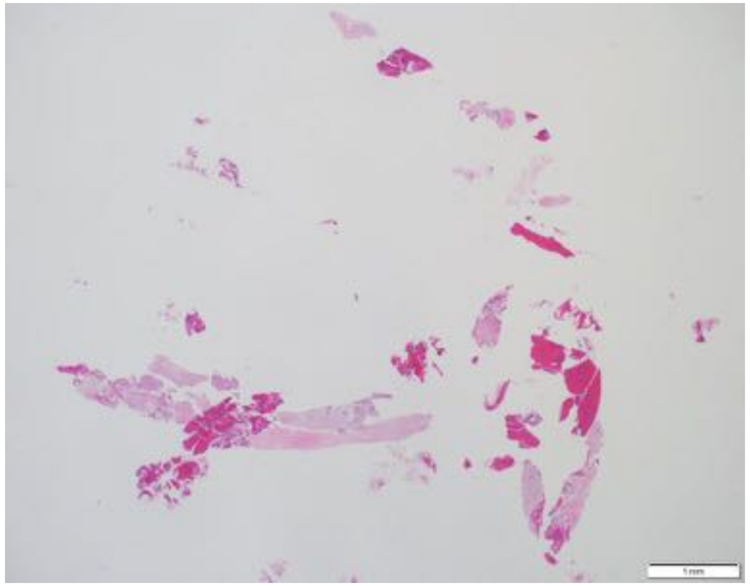

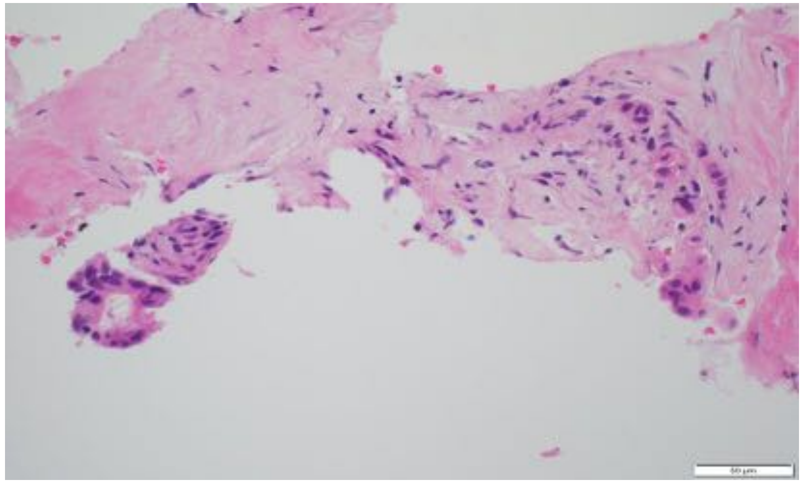

| | |
|---|---|
| **Patient Number** | P13 |
| **Age** | 78 |
| **Gender** | Male |
| **Stage at Diagnosis** | III |
| **Treatment before tissue collection** | No |
| **Tissue site** | Pancreas |
| **Procedure** | Biopsy |
| **Pathology** | Poorly differentiated adenocarcinoma |
| **Mutations** | TP53 c.880del |

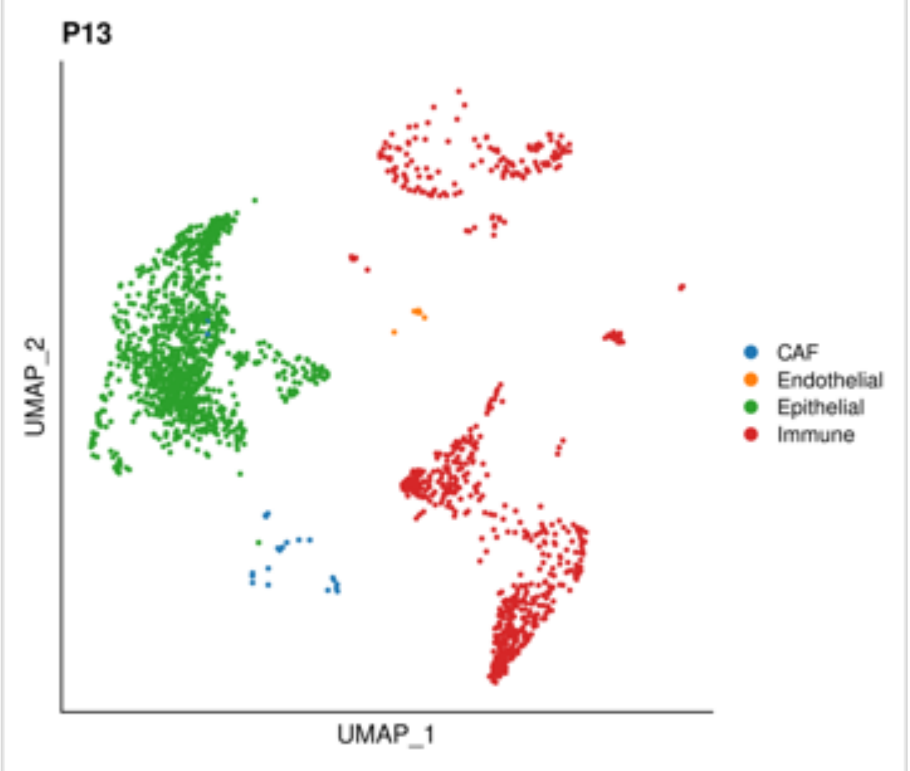


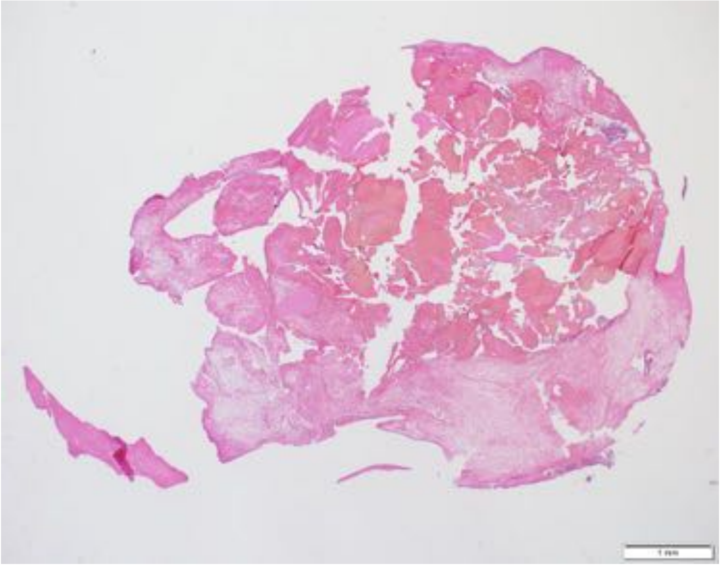

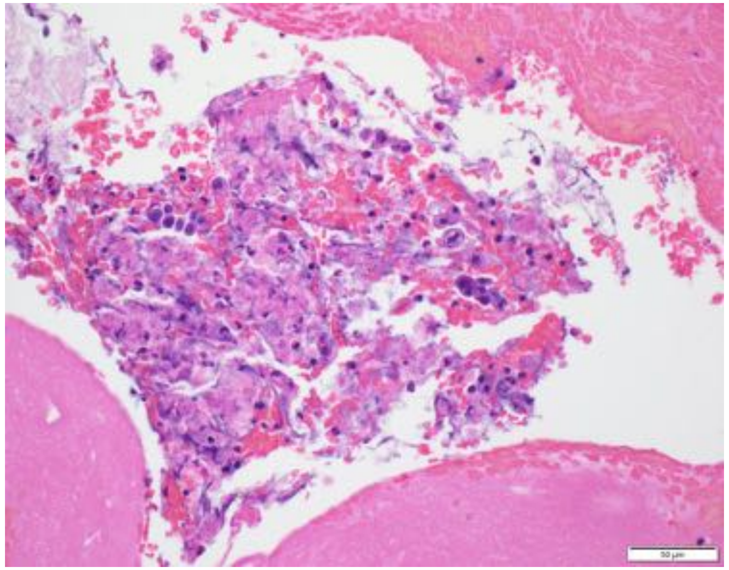

| | |
|---|---|
| **Patient Number** | P14 |
| **Age** | 67 |
| **Gender** | Female |
| **Stage at Diagnosis** | IB |
| **Treatment before tissue collection** | Yes |
| **Therapeutics** | FFX-based |
| **Tissue site** | Pancreas |
| **Procedure** | Resection |
| **Pathology** | Well differentiated adenocarcinoma |
| **Mutations** | WT |

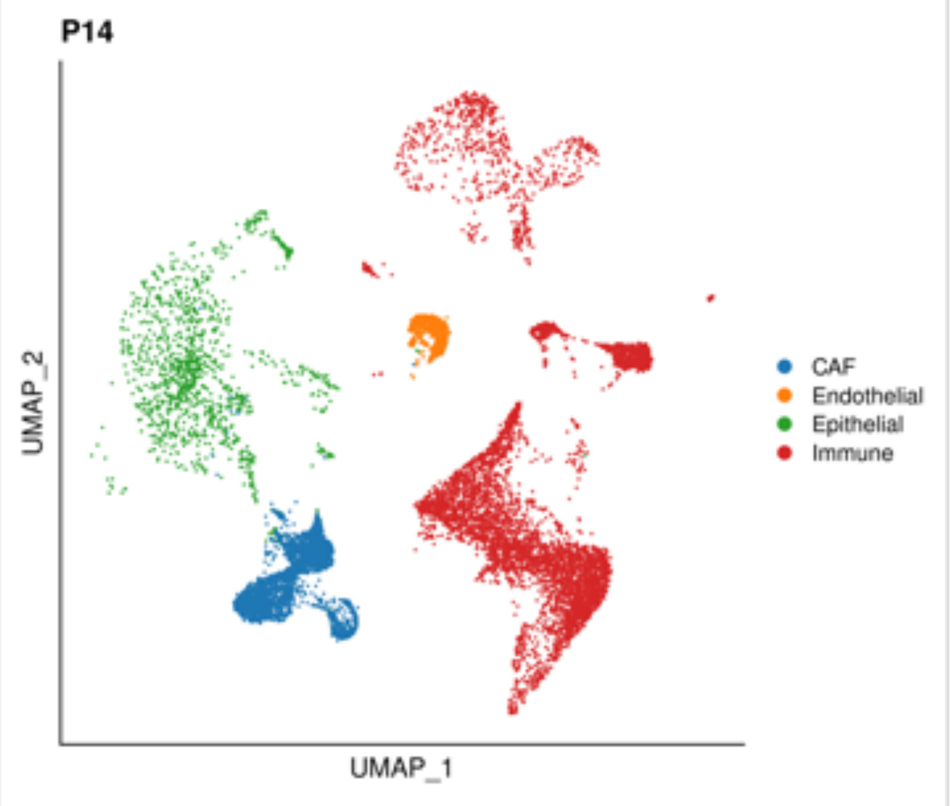


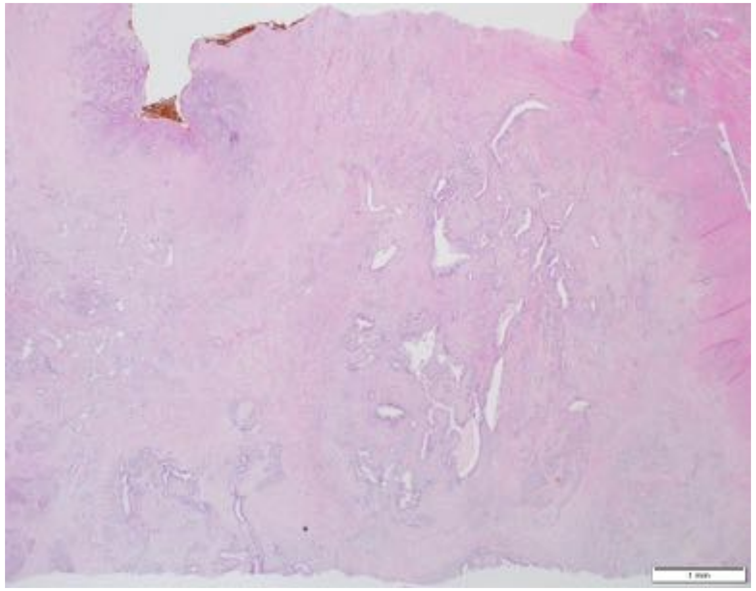

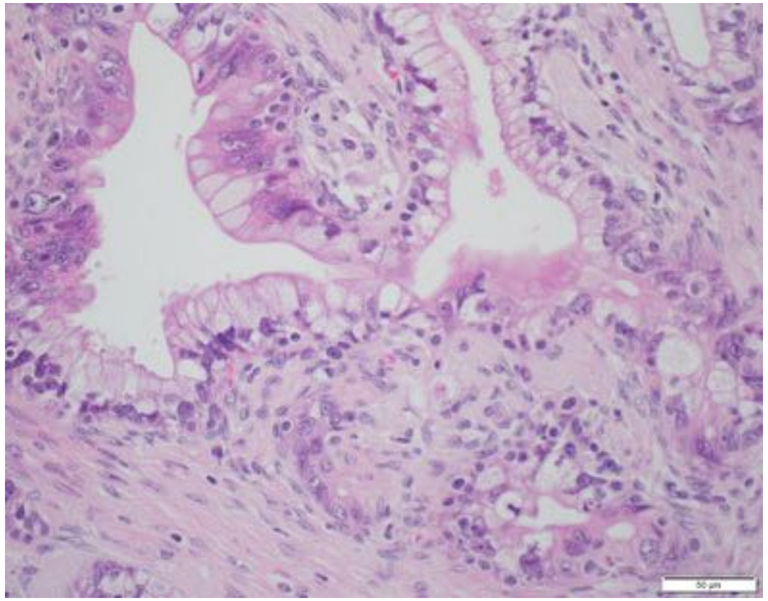

| | |
|---|---|
| **Patient Number** | P15 |
| **Age** | 58 |
| **Gender** | Female |
| **Stage at Diagnosis** | III |
| **Treatment before tissue collection** | No |
| **Tissue site** | Pancreas |
| **Procedure** | Resection |
| **Pathology** | Poorly differentiated adenocarcinoma |
| **Mutations** | KRAS G12V, TP53 C242Y, SMAD4 c.454+1G>A, CDKN2A R80* |

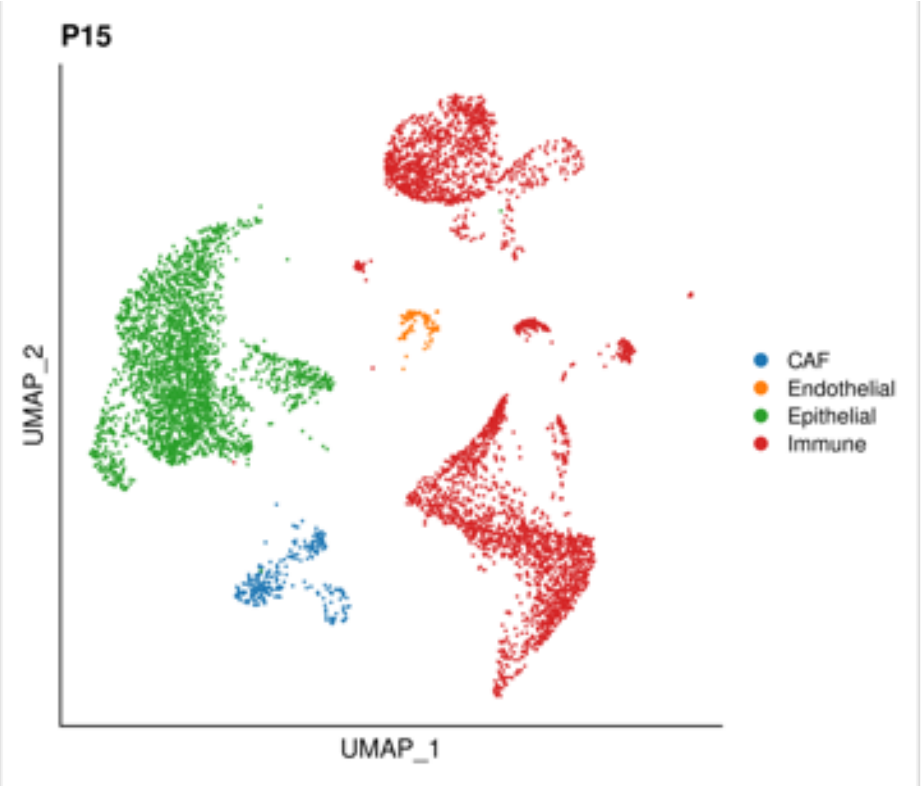


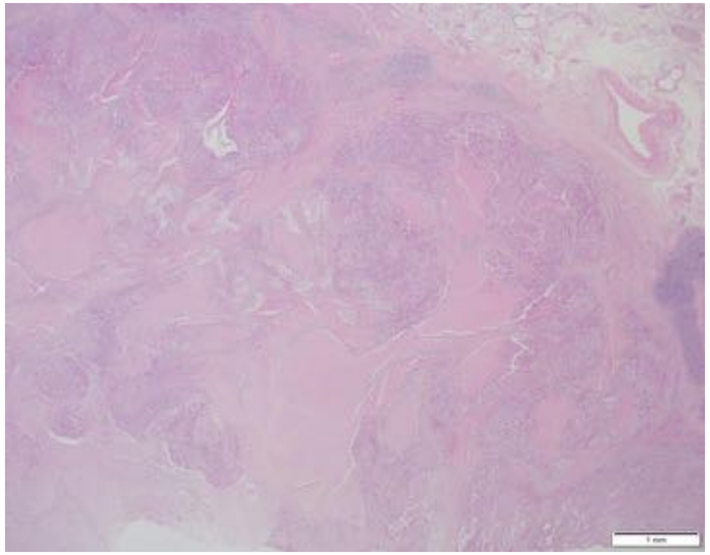

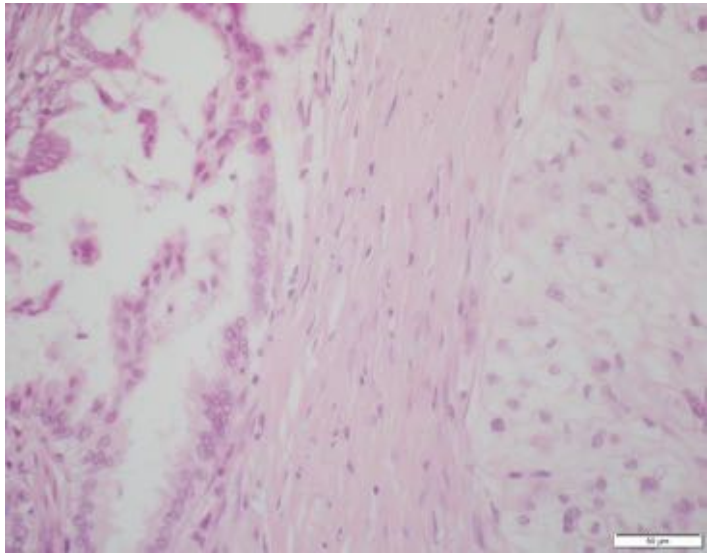

| | |
|---|---|
| **Patient Number** | P16 |
| **Age** | 69 |
| **Gender** | Female |
| **Stage at Diagnosis** | IV |
| **Treatment before tissue collection** | No |
| **Tissue site** | Liver |
| **Procedure** | Biopsy |
| **Pathology** | Moderately to poorly differentiated adenocarcinoma |
| **Mutations** | KRAS G12V, TP53 C135F |

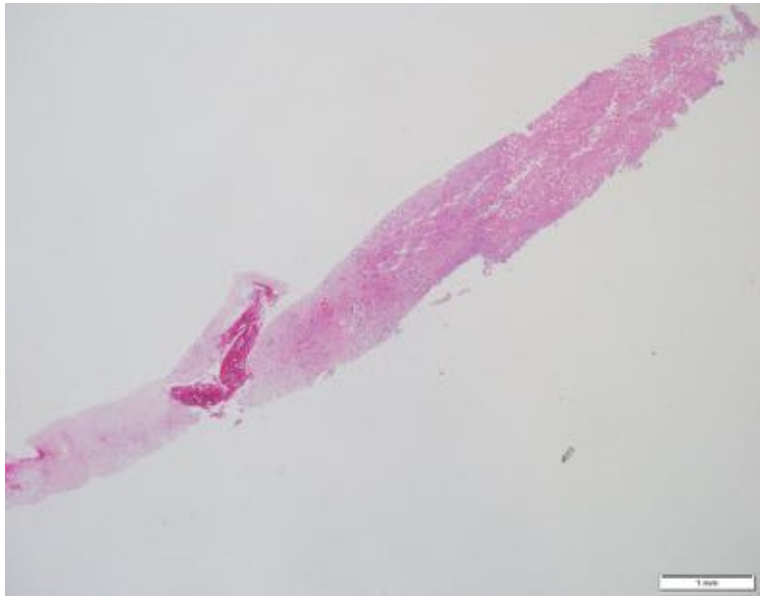

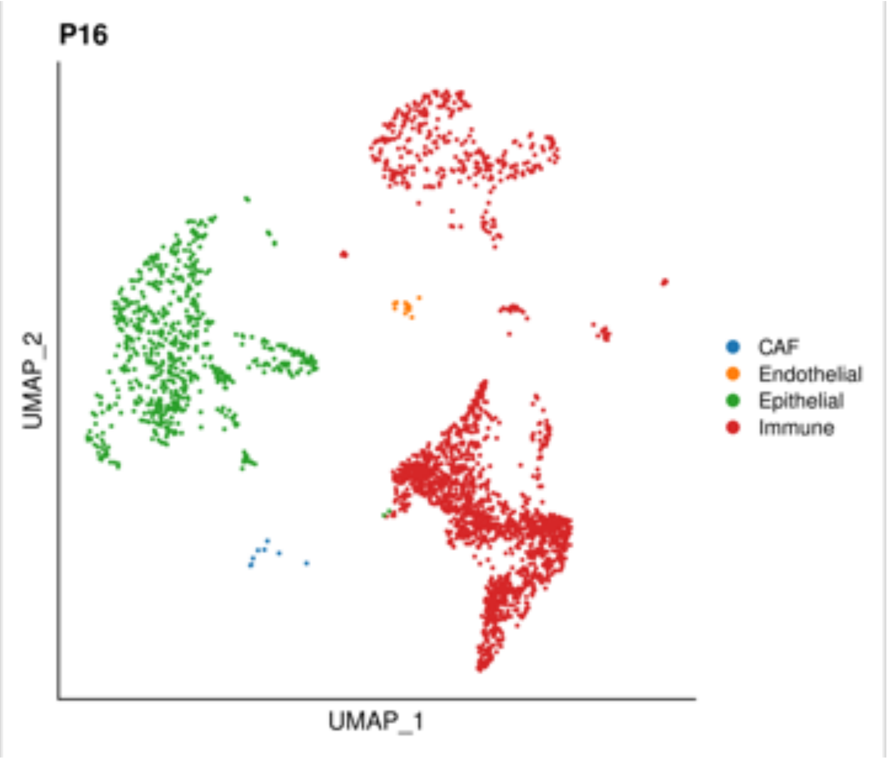


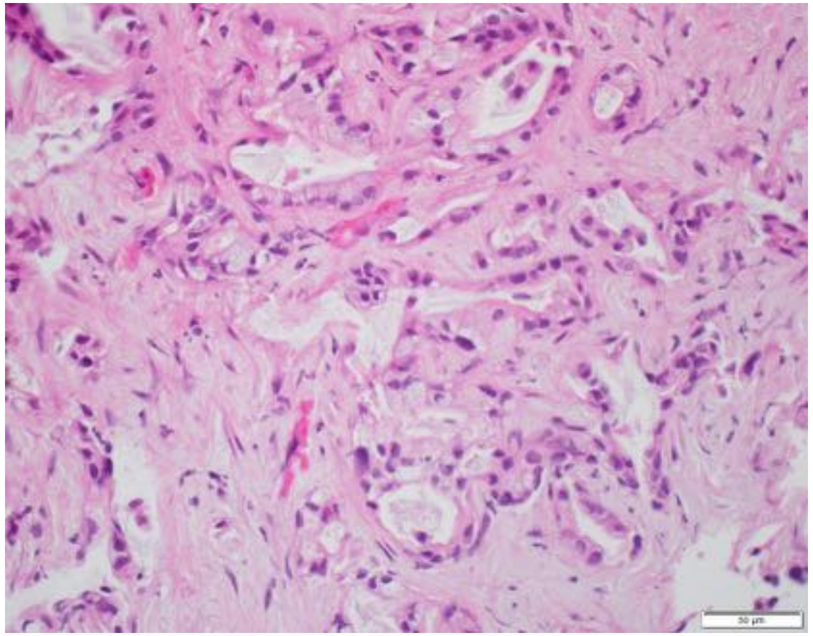

| | |
|---|---|
| **Patient Number** | P17 |
| **Age** | 64 |
| **Gender** | Male |
| **Stage at Diagnosis** | IV |
| **Treatment before tissue collection** | Yes |
| **Therapeutics** | FFX-based |
| **Tissue site** | Liver |
| **Procedure** | Biopsy |
| **Pathology** | Moderately differentiated adenocarcinoma |
| **Mutations** | N/A |

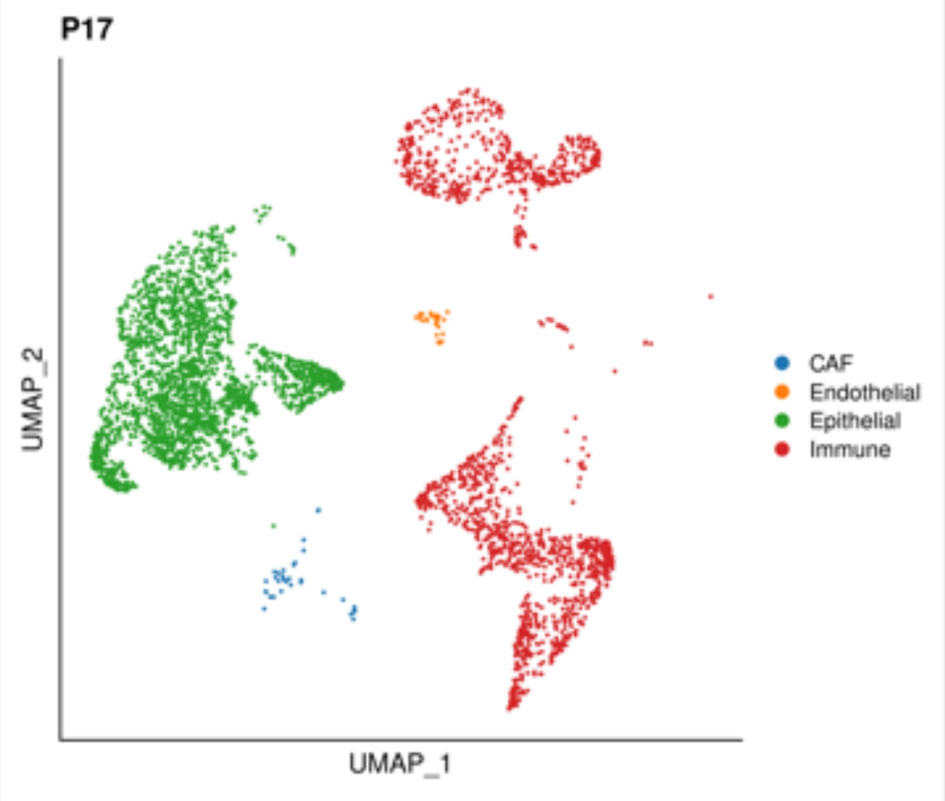


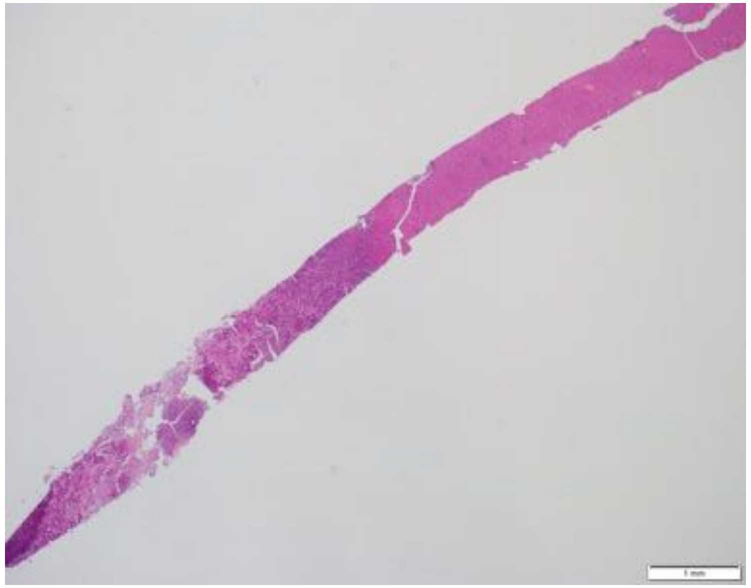

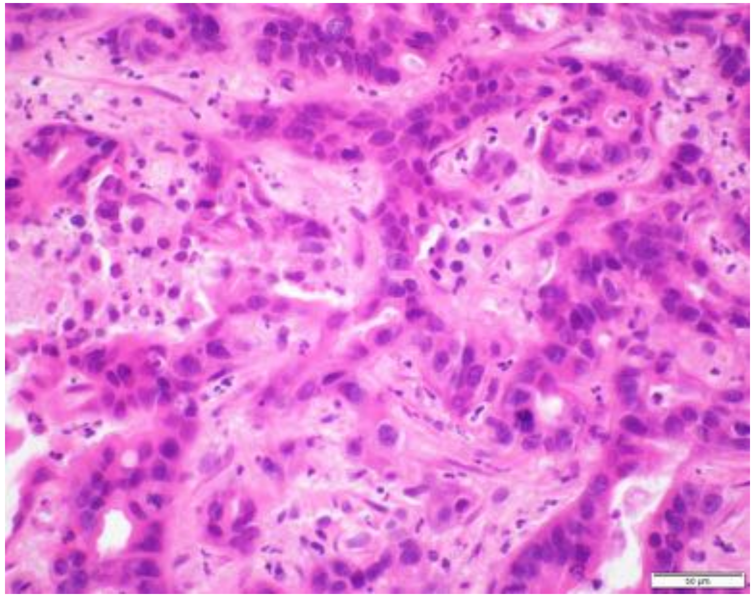

| | |
|---|---|
| **Patient Number** | P18 |
| **Age** | 75 |
| **Gender** | Male |
| **Stage at Diagnosis** | IV |
| **Treatment before tissue collection** | No |
| **Tissue site** | Liver |
| **Procedure** | Biopsy |
| **Pathology** | Poorly differentiated adenocarcinoma |
| **Mutations** | KRAS G12D, TP53 R248Q, CDKN2A loss |

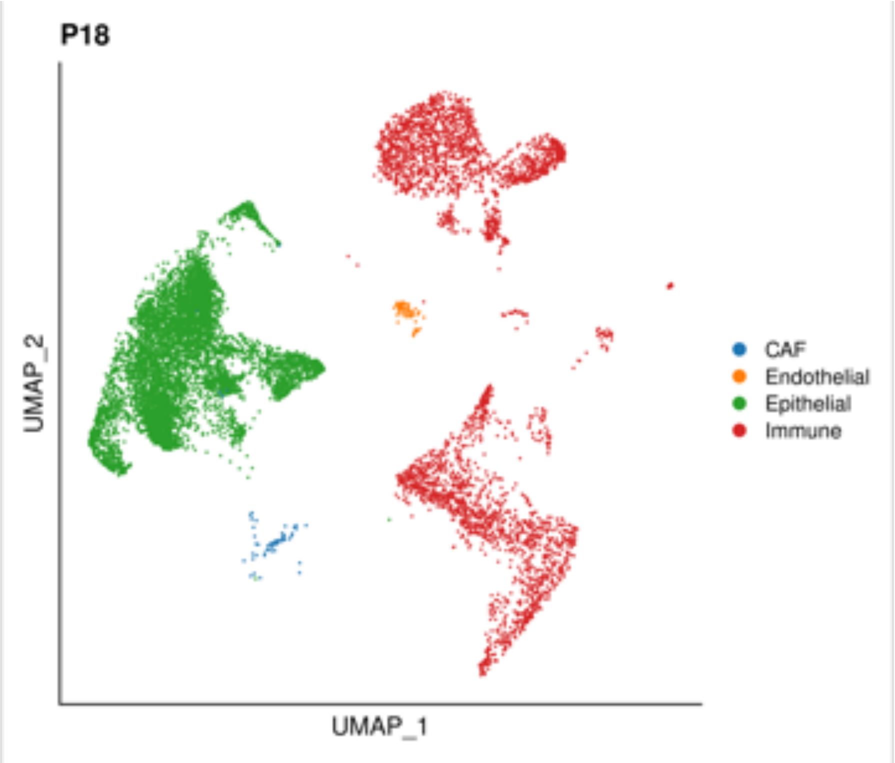


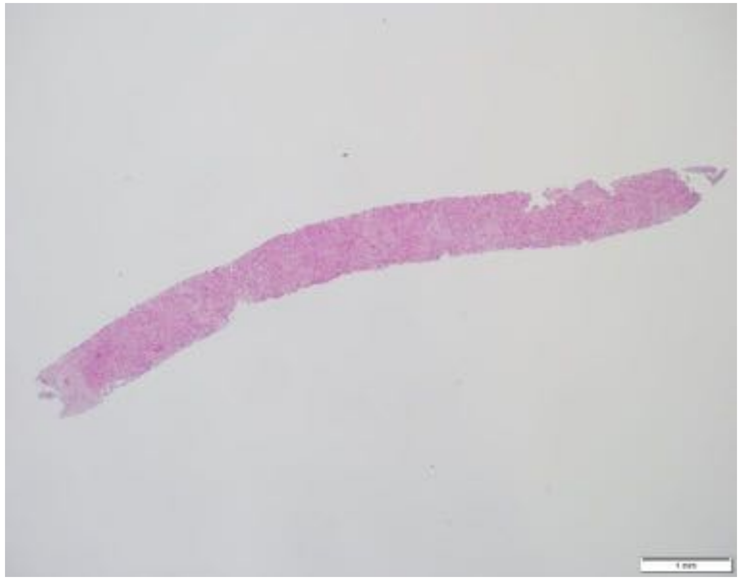

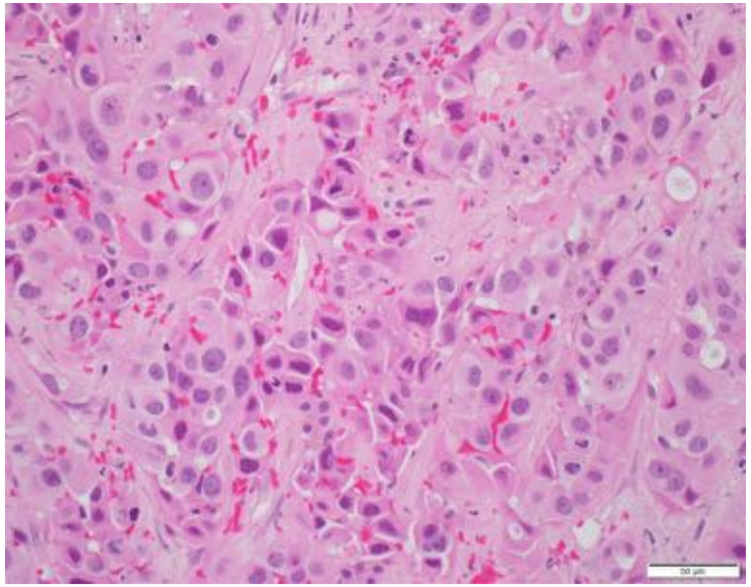

| | |
|---|---|
| **Patient Number** | P19 |
| **Age** | 60 |
| **Gender** | Female |
| **Stage at Diagnosis** | IB |
| **Treatment before tissue collection** | No |
| **Tissue site** | Pancreas |
| **Procedure** | Resection |
| **Pathology** | Moderately to poorly differentiated adenocarcinoma |
| **Mutations** | KRAS G12V, TP53 c.993+1G>A |

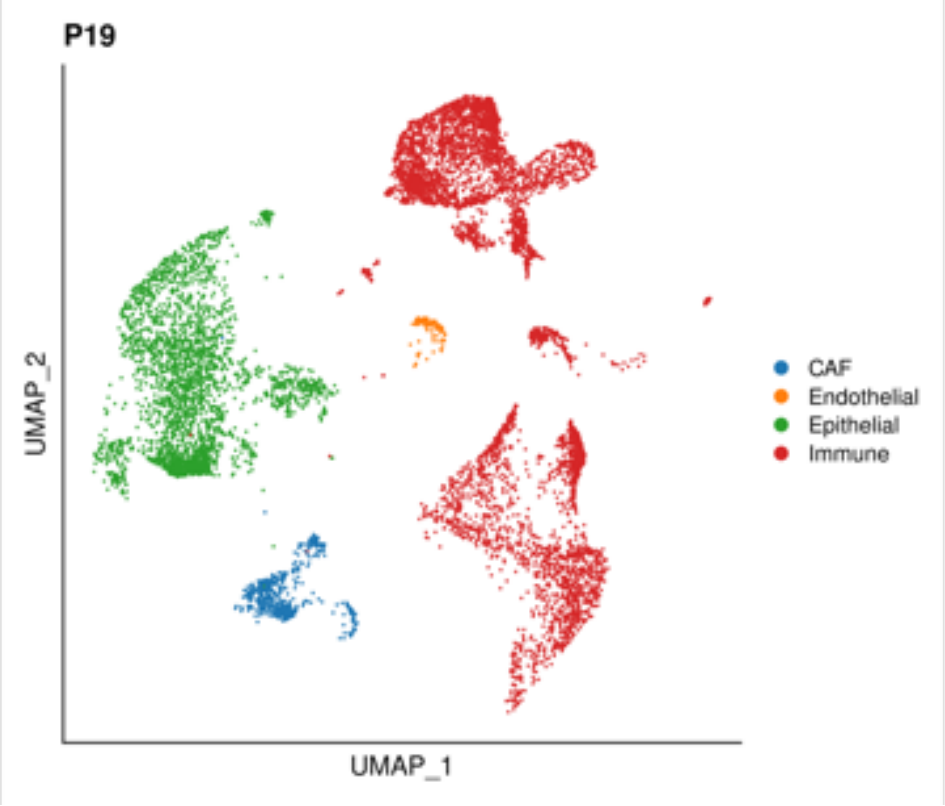


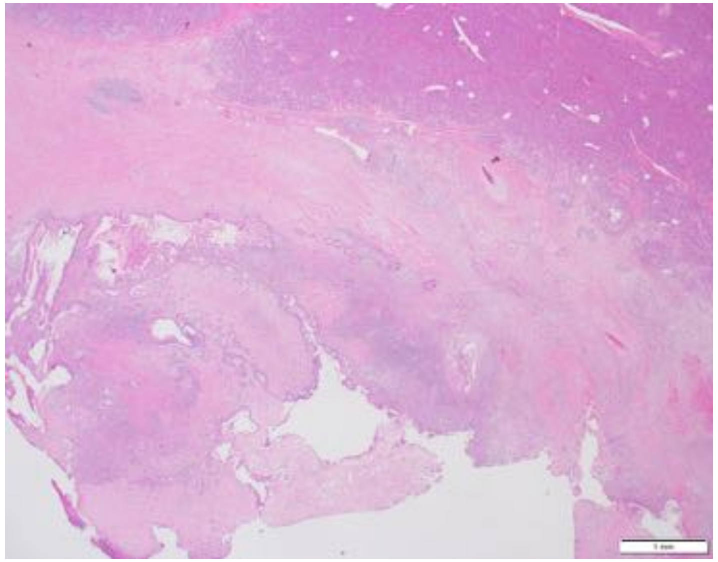

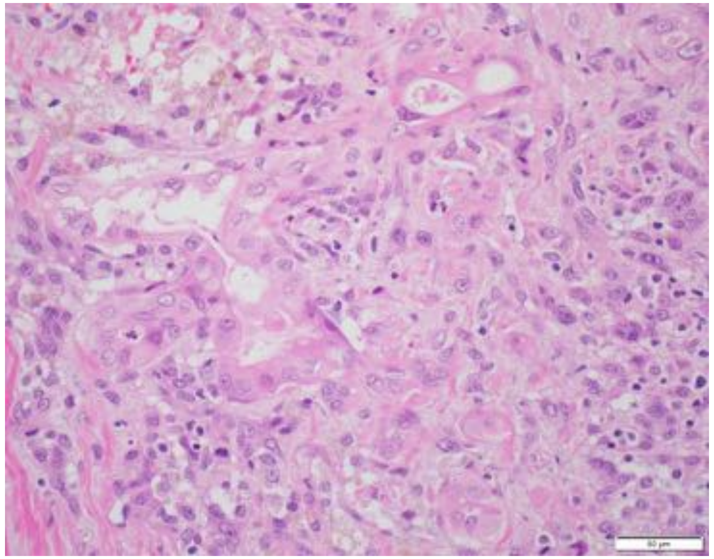

| | |
|---|---|
| **Patient Number** | P20 |
| **Age** | 66 |
| **Gender** | Female |
| **Stage at Diagnosis** | IV |
| **Treatment before tissue collection** | No |
| **Tissue site** | Pancreas |
| **Procedure** | Biopsy |
| **Pathology** | Moderately differentiated adenocarcinoma |
| **Mutations** | KRAS G12D, TP53 R249I |

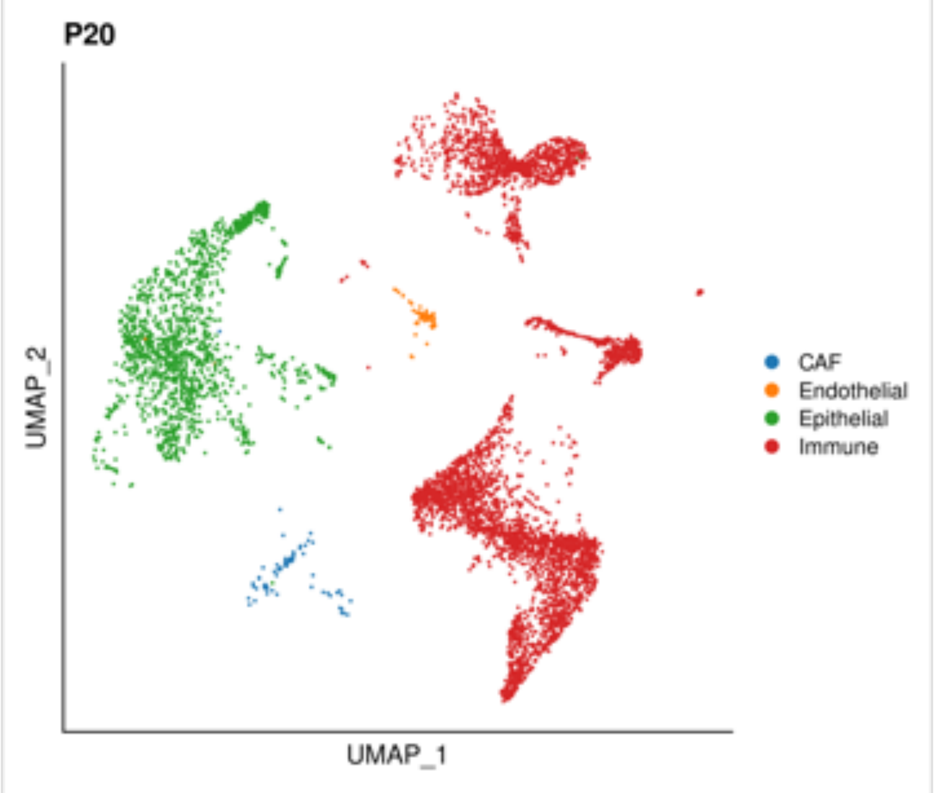


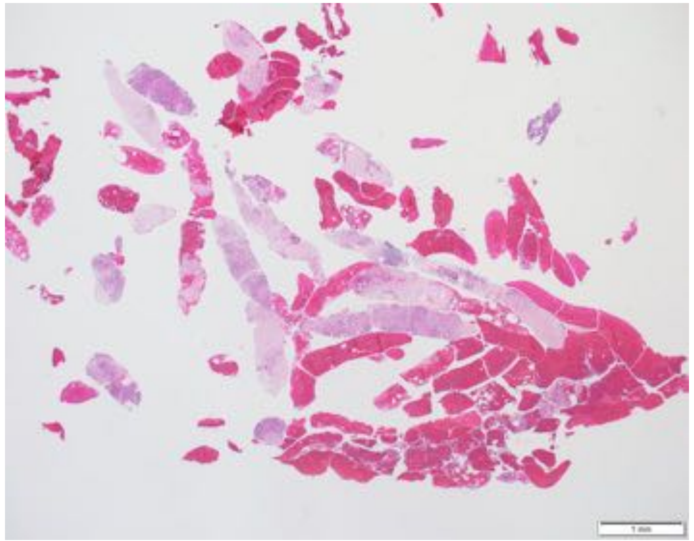

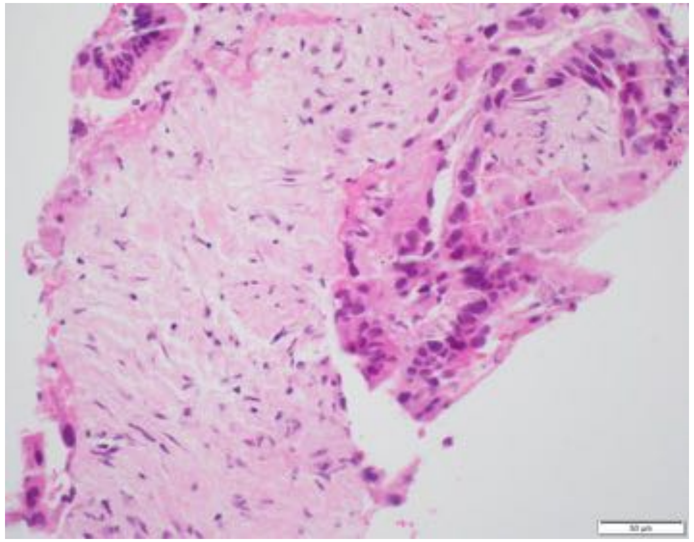

| | |
|---|---|
| **Patient Number** | P21 |
| **Age** | 69 |
| **Gender** | Male |
| **Stage at Diagnosis** | IV |
| **Treatment before tissue collection** | No |
| **Tissue site** | Liver |
| **Procedure** | Biopsy |
| **Pathology** | Moderately to poorly differentiated adenocarcinoma |
| **Mutations** | KRAS G12D, TP53 H168L |

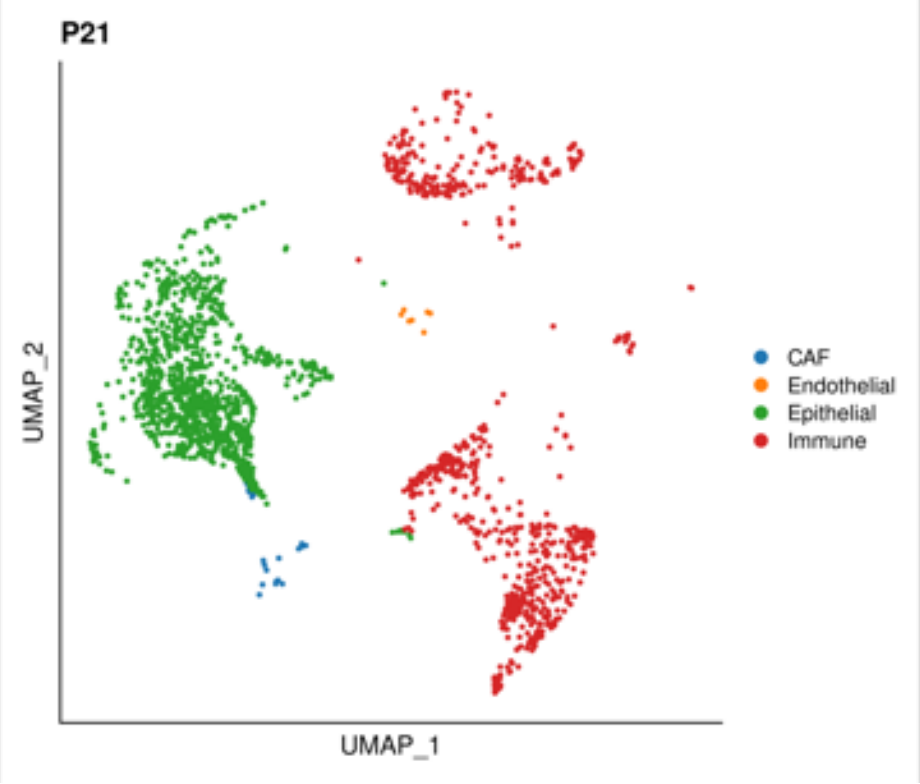


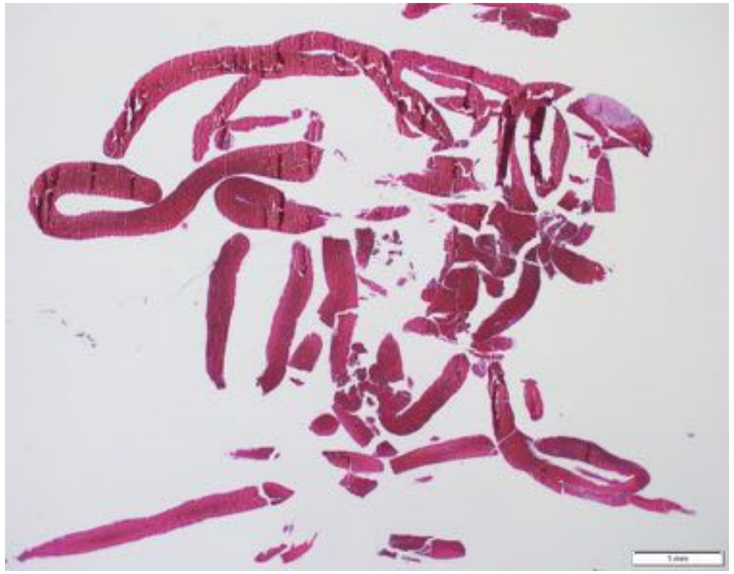

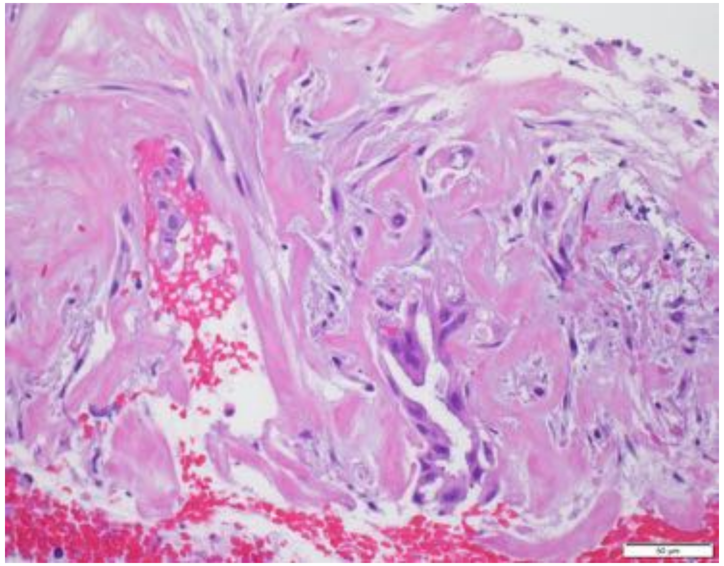

| | |
|---|---|
| **Patient Number** | P22 |
| **Age** | 78 |
| **Gender** | Male |
| **Stage at Diagnosis** | III |
| **Treatment before tissue collection** | No |
| **Tissue site** | Pancreas |
| **Procedure** | Biopsy |
| **Pathology** | Moderate differentiated adenocarcinoma |
| **Mutations** | N/A |

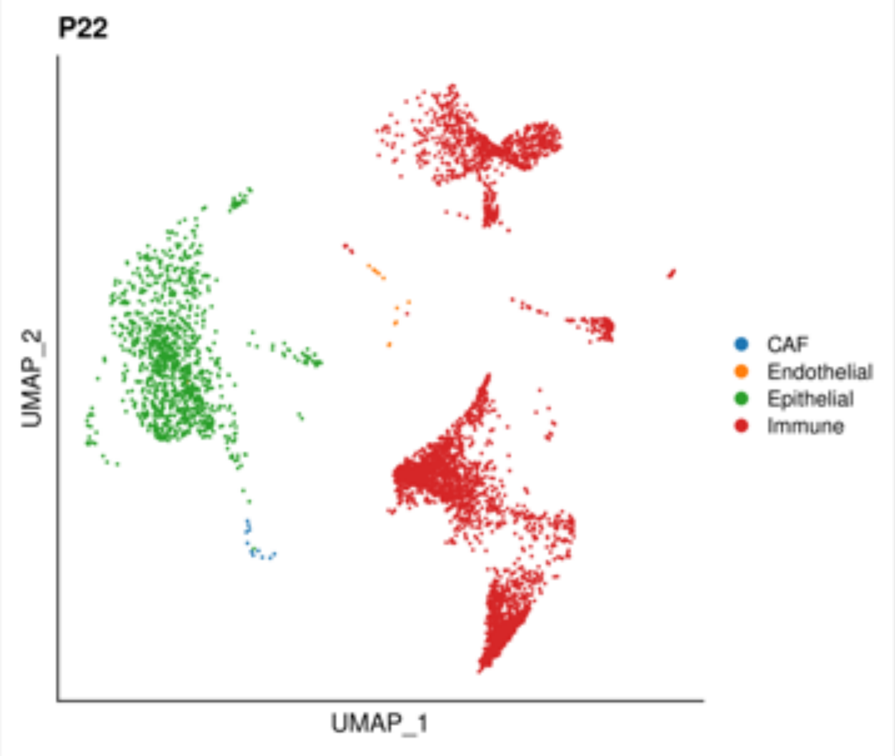


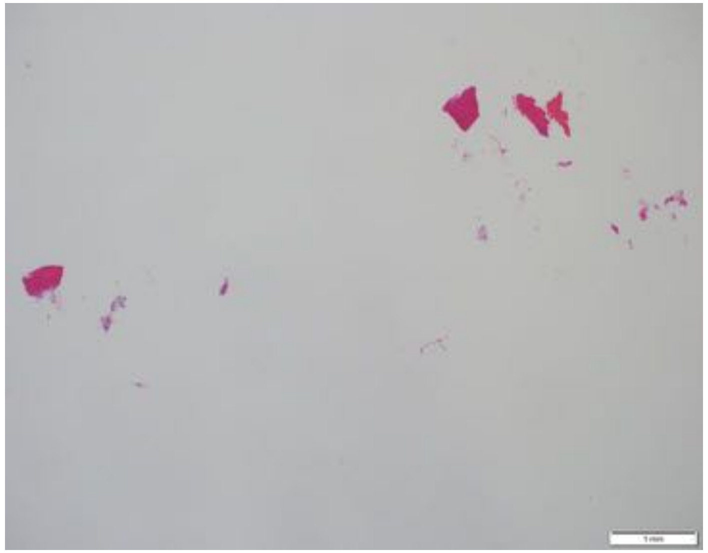

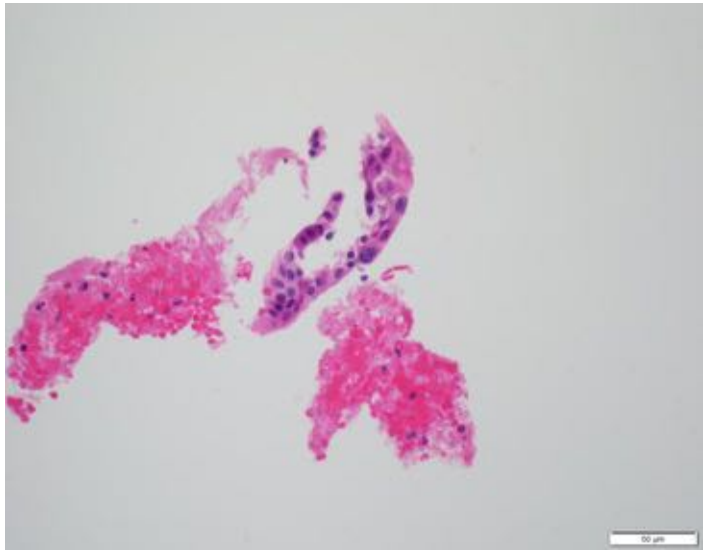

| | |
|---|---|
| **Patient Number** | P23 |
| **Age** | 66 |
| **Gender** | Female |
| **Stage at Diagnosis** | IB |
| **Treatment before tissue collection** | No |
| **Tissue site** | Pancreas |
| **Procedure** | Resection |
| **Pathology** | Well to moderately differentiated adenocarcinoma |
| **Mutations** | KRAS G12D, TP53 I195S |

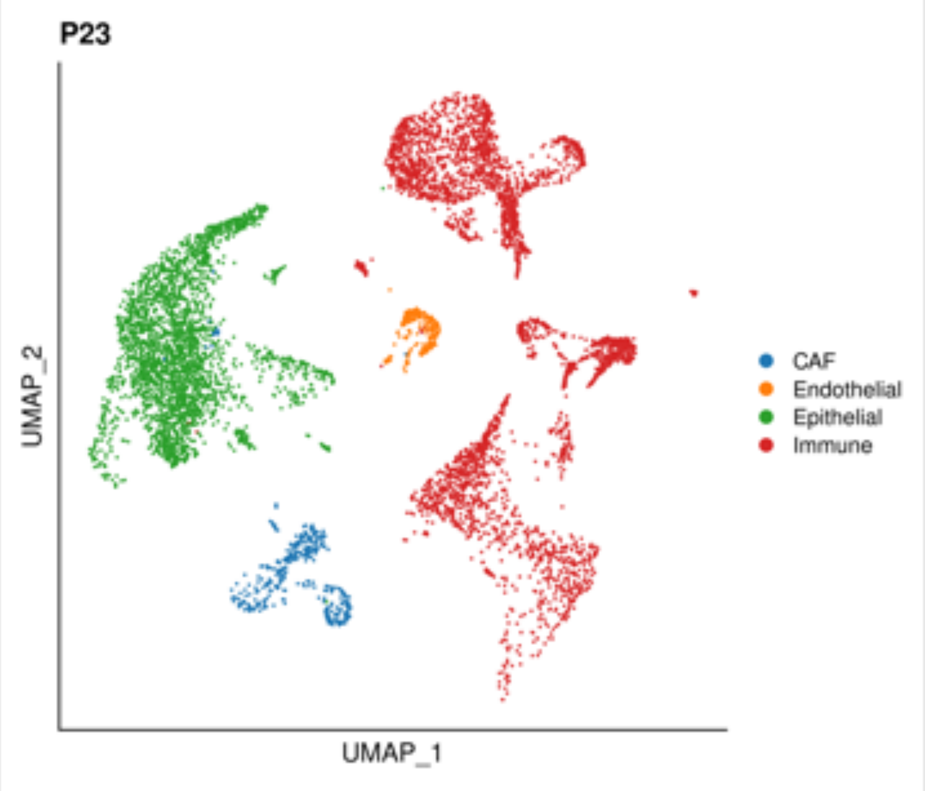


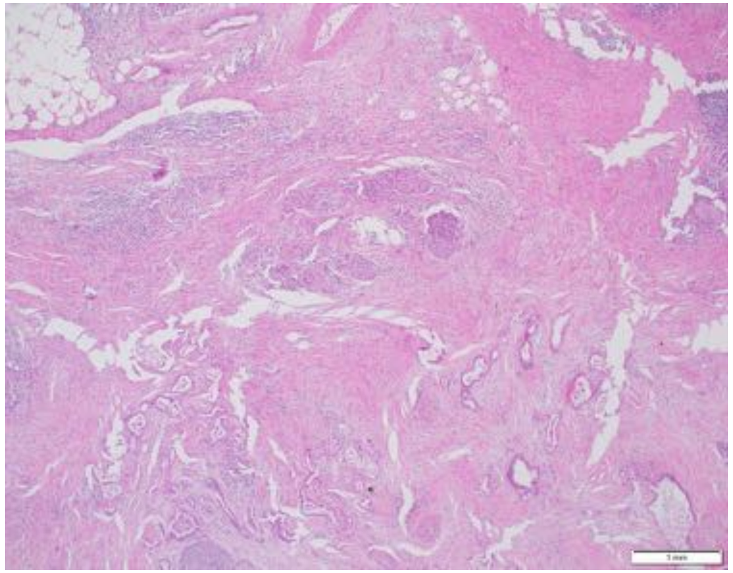

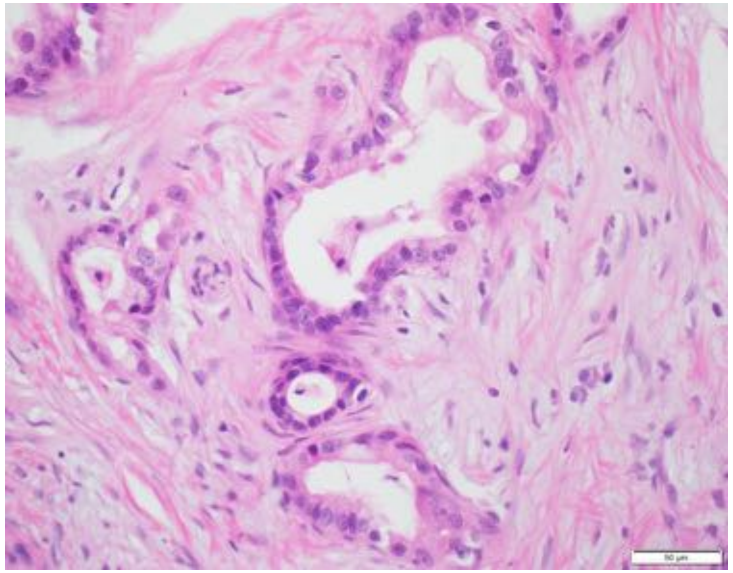

| | |
|---|---|
| **Patient Number** | P24 |
| **Age** | 65 |
| **Gender** | Female |
| **Stage at Diagnosis** | IV |
| **Treatment before tissue collection** | No |
| **Tissue site** | Liver |
| **Procedure** | Biopsy |
| **Pathology** | Moderately to poorly differentiated |
| **Mutations** | KRAS Q61K, TP53 C176Y |

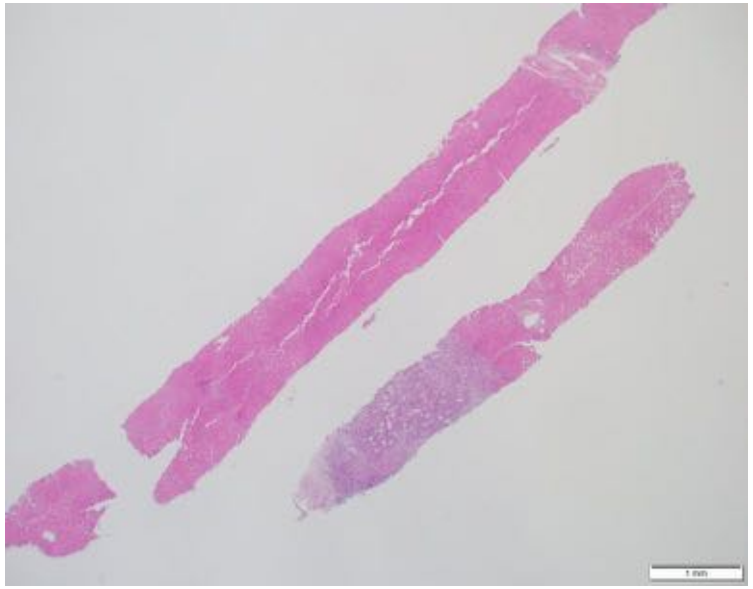

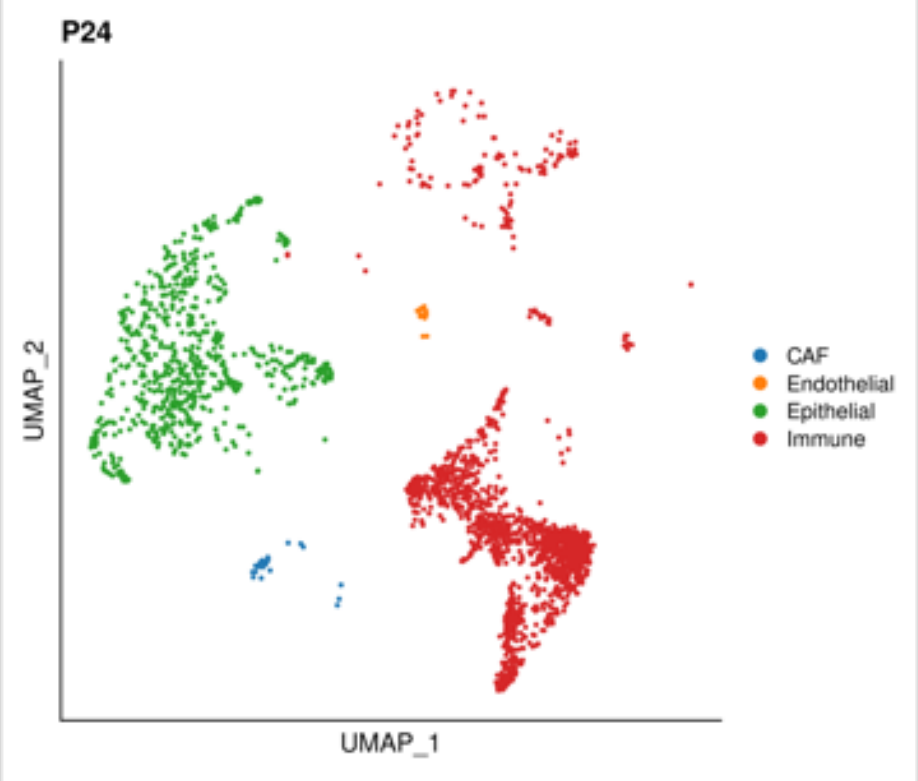


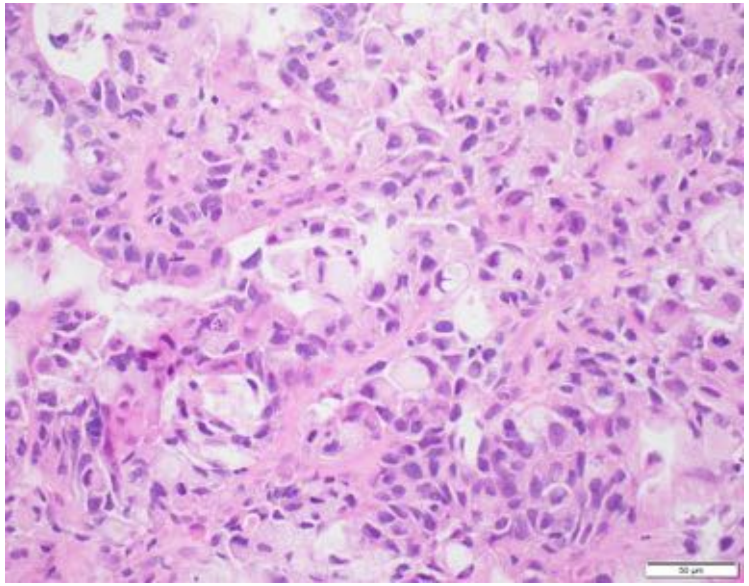

| | |
|---|---|
| **Patient Number** | P25 |
| **Age** | 48 |
| **Gender** | Male |
| **Stage at Diagnosis** | IV |
| **Treatment before tissue collection** | No |
| **Tissue site** | Liver |
| **Procedure** | Biopsy |
| **Pathology** | Poorly differentiated adenocarcinoma |
| **Mutations** | KRAS G12V |

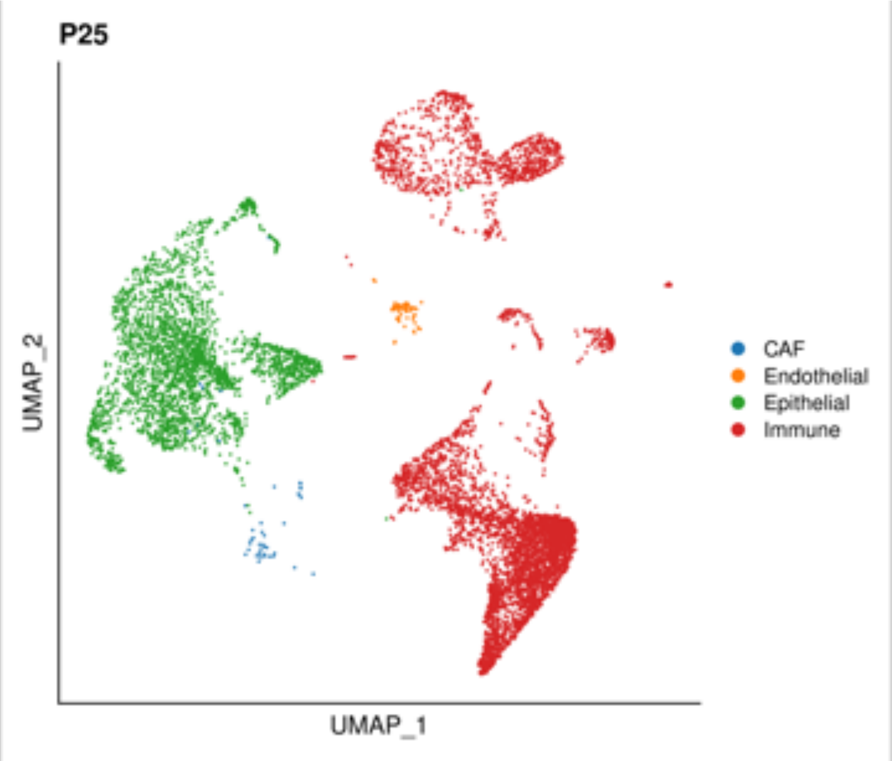


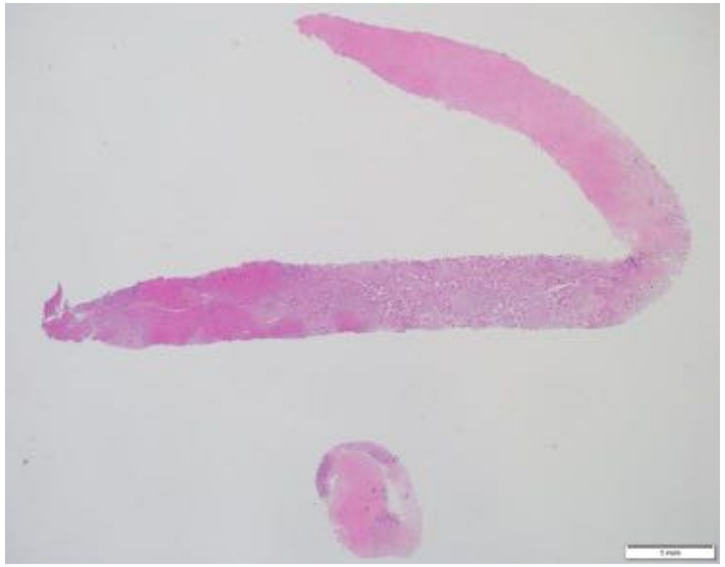

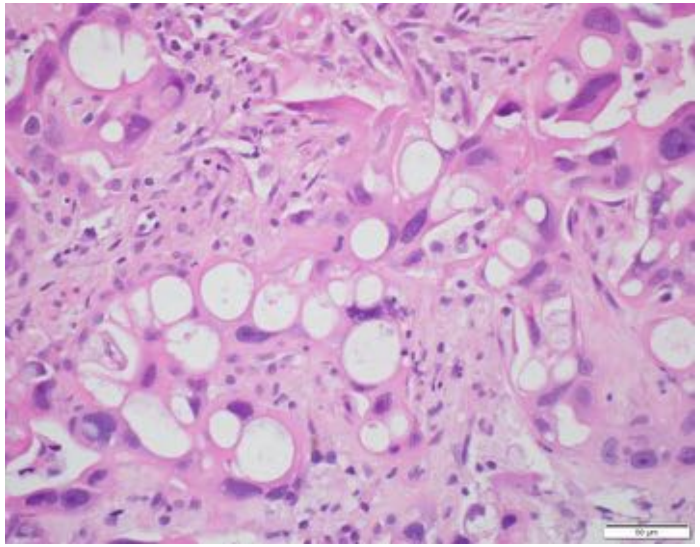

| | |
|---|---|
| **Patient Number** | P26 |
| **Age** | 64 |
| **Gender** | Male |
| **Stage at Diagnosis** | IV |
| **Treatment before tissue collection** | No |
| **Tissue site** | Pancreas |
| **Procedure** | Biopsy |
| **Pathology** | Moderately differentiated adenocarcinoma |
| **Mutations** | KRAS G12V |

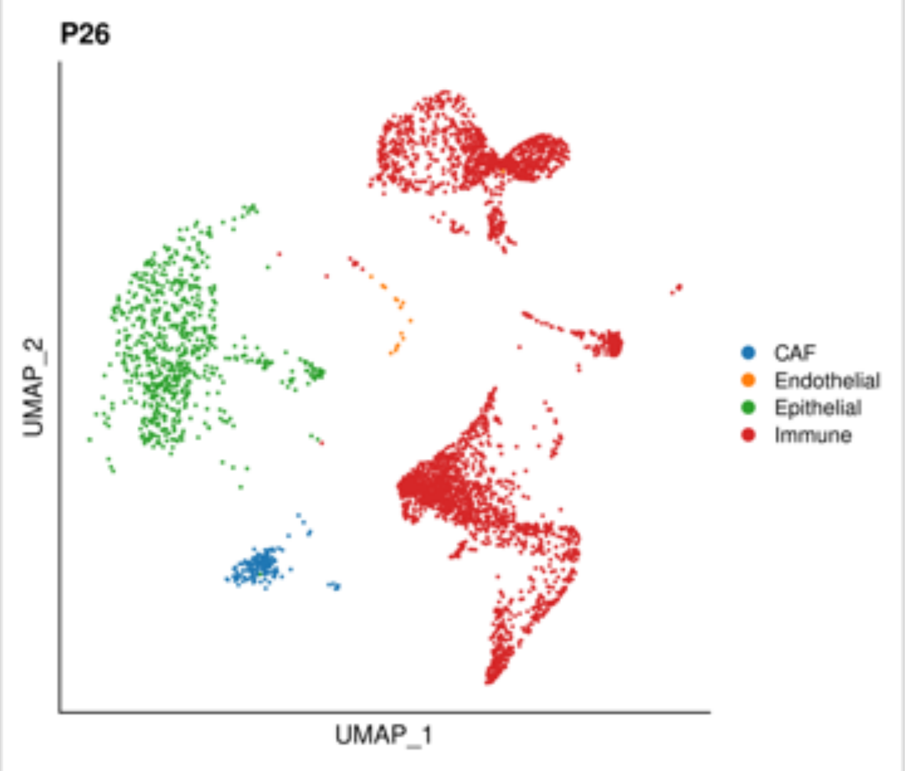


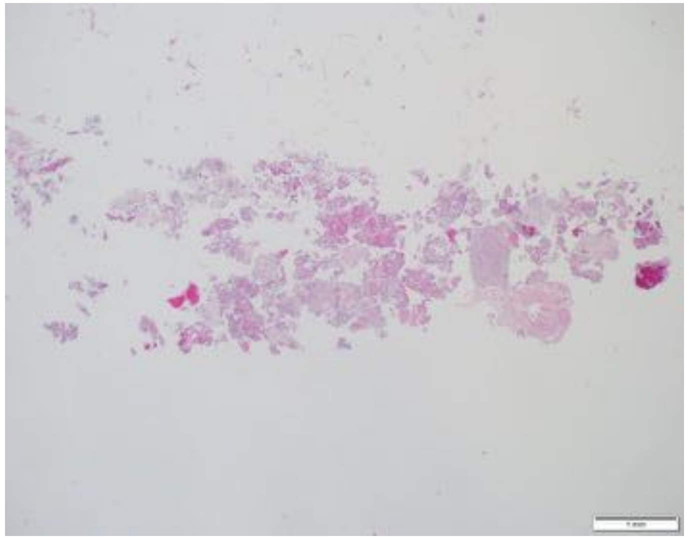

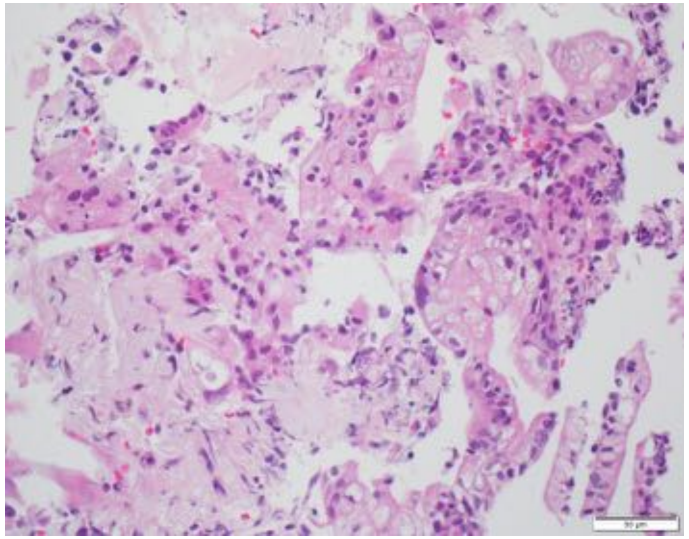

| | |
|---|---|
| **Patient Number** | P27 |
| **Age** | 72 |
| **Gender** | Female |
| **Stage at Diagnosis** | IV |
| **Treatment before tissue collection** | No |
| **Tissue site** | Liver |
| **Procedure** | Biopsy |
| **Pathology** | Moderately to poorly differentiated adenocarcinoma |
| **Mutations** | WT |

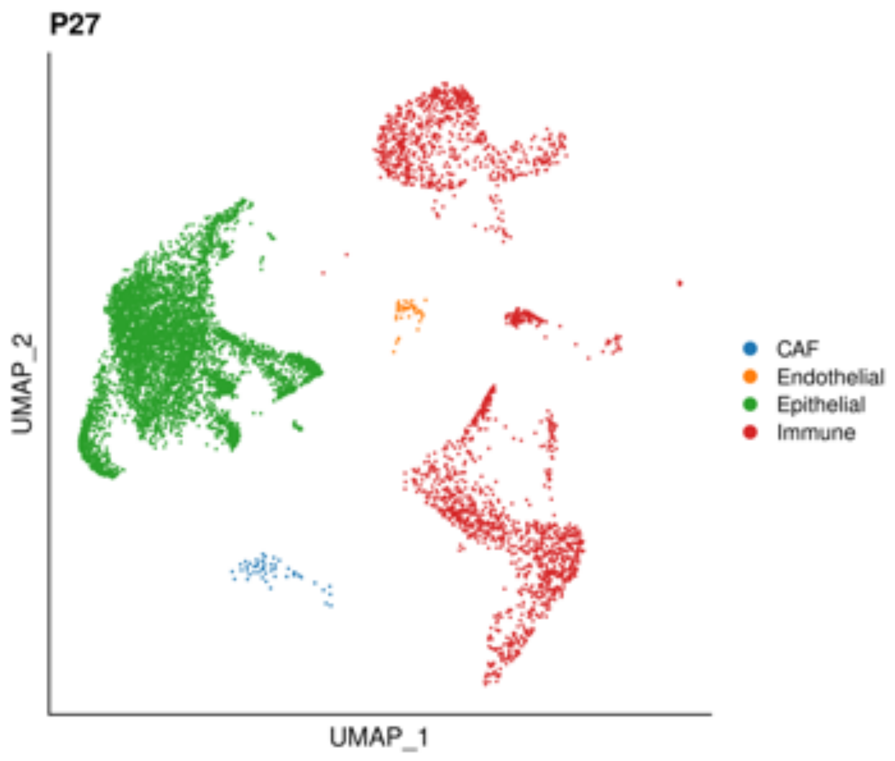


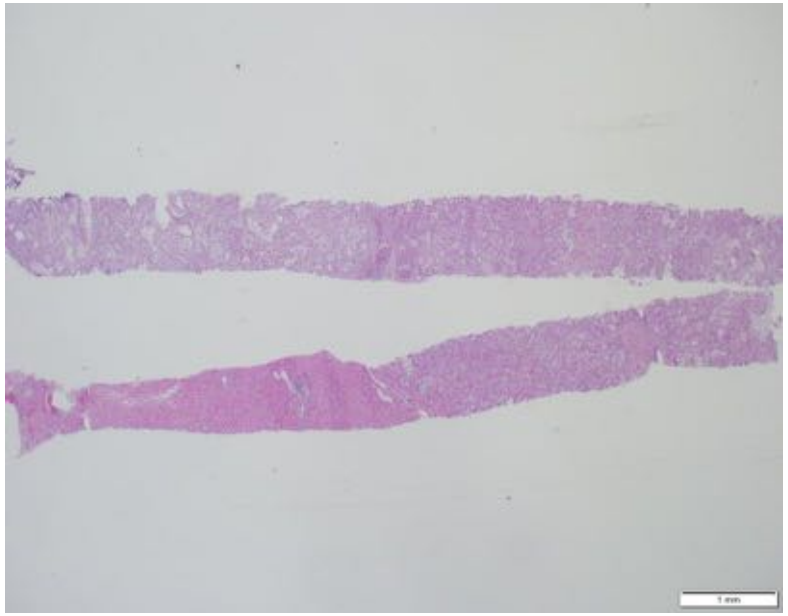

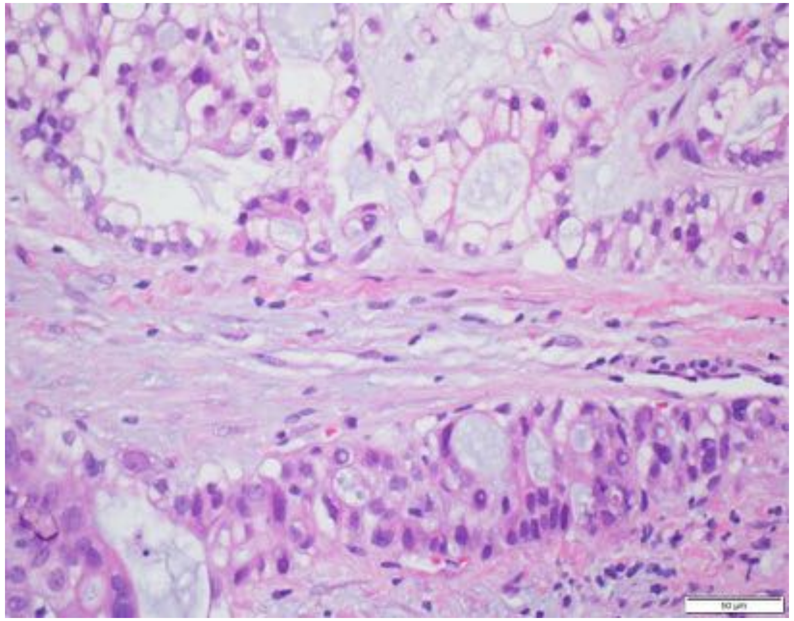
